# Charting the small-molecule universe from mass spectra with neuro-symbolic AI

**DOI:** 10.64898/2026.08.05.743095

**Authors:** Utku U. Acikalin, Dieqiao Feng, Aaron M. Ferber, Goncalo J. Gouveia, Tyler J. Schwertfeger, Di Chen, Delia Qu, Marissa A. Fontaine, Yingheng Wang, Richard A. Bernstein, Haofan Wang, Tae-Hyung Won, Christopher N. Parkhurst, Bart Selman, David Artis, Frank C. Schroeder, Carla P. Gomes

## Abstract

Mass spectrometry (MS) has revealed millions of small organic molecules across organisms, yet most remain uncharacterized, limiting progress in biology and medicine. Despite computational advances, MS workflows rely heavily on expert input and reference libraries that cover only a fraction of known chemical space. Here, we introduce AIMe (AI Molecule Explorer), a multi-agent neuro-symbolic AI framework that transforms the interpretation of unknown spectra into an omics-scale exploration across the known structural space, providing chemically interpretable annotations. At its core, AIMe combines chemical reasoning with structure-informed learning to predict MS^2^ spectra by modeling fragmentation as a sequence of actions, outperforming existing methods. AIMe dynamically constructs fragmentation pathways by assigning likelihoods to individual fragmentation actions, linking spectral peaks to explicit fragment molecular formulas and structures. At scale, AIMe predicted MS^2^ spectra for over 100 million small organic molecules in PubChem and organized them into MS^2^KOSMOS, a substructure-informed community resource comprising over 800 million predicted spectra that expands the searchable small-molecule universe by roughly three orders of magnitude relative to experimental libraries. Analogous to sequence homology-based searches in genomics and proteomics, AIMe maps unknown spectra to molecular neighborhoods in MS^2^KOSMOS. Exact- formula indexing enables ranked retrieval of candidates and related structures, with peak-level structural and fragmentation-pathway annotations. Applied to mouse microbiota-dependent metabolites, AIMe enabled putative annotation of knowns and guided structure elucidation of unknowns, revealing previously unreported types of microbiota-dependent polyamines that also occur in humans. At repository scale, AIMe enabled putative annotation of roughly a third of 7 million spectral clusters representing most of the unknowns in the GNPS database. By extending MS^2^ annotation beyond curated-library matching to interpretable search across the known small-molecule universe, AIMe accelerates discovery and large-scale exploration of small molecules across biomedicine, agriculture, and ecology.

## Main

The identification of the molecular structures of small organic molecules remains a major bottleneck to advancing chemistry, biology, and medicine^1–3^. The widespread adoption of high-resolution tandem mass spectrometry (MS^2^) has revealed that microorganisms, plants, and animals produce millions of small organic molecules that may play important biological functions, yet the chemical structures of the vast majority remain undetermined^4–9^. Similarly, structure elucidation of unknown compounds remains a critical challenge for drug discovery^10^, food safety^11^, and environmental analyses^12^. In a typical metabolomics study, samples are analyzed by combining liquid chromatography (LC) with MS^2^, followed by comparing the acquired spectra against MS^2^ reference libraries. However, available reference libraries contain spectra for fewer than 1% of known compounds^7,9,13^, representing only a tiny fraction of the astronomically larger space of all chemically possible small molecules^10,14^. As a result, for any given complex biological sample, more than 80% of detected metabolites remain usually unidentified^7,9,15^. In contrast to DNA or proteins, whose structures are constrained by a small number of building blocks, the diversity of small organic molecules is limited only by the basic chemical properties of the constituting elements. As a result, a single molecular formula can often correspond to thousands of distinct structures^15,16^. Therefore, expanding experimentally curated reference libraries alone is unlikely to solve the structure elucidation challenge.

Here we propose an alternative vision, in which structure elucidation is facilitated by the ability to predict, at scale and with high fidelity, the MS^2^ spectra of any previously detected or hypothetical compound, to then organize these predictions into a comprehensive spectral space that can be rapidly queried for spectral and structural similarities. Our approach builds on early efforts at the intersection of AI and mass spectrometry^6,17–22^, including the landmark Dendral expert system, which laid the foundation for computational approaches to molecular formula inference and structure elucidation through rule-based reasoning^23^. Although the field has advanced considerably since, current MS^2^-based workflows for small-molecule structure elucidation remain constrained by challenges in both accuracy and scalability^24–27^.

To address the structure elucidation challenge, we developed the *AI Molecule Explorer* (AIMe), a multi-agent, neuro-symbolic AI^28^ framework that integrates symbolic chemical reasoning with structure-informed neural learning^29^ to enable MS^2^ prediction, interpretation, and annotation at omics scale. AIMe consists of modular agents that collaboratively reason over MS^2^ fragmentation, molecular structure, and spectral relationships (**Fig. 1a-d**). At its core, AIMe’s *DeepMS^2^Reasoner* combines symbolic construction of chemically feasible fragmentation pathways with neural-network-based assignment of fragmentation-action likelihoods, enabling it to jointly predict MS^2^ spectra and fragmentation pathways **(Fig. 1b)**. By explicitly considering stepwise fragmentation actions during pathway construction, including bond-cleavage, excision, and hydrogen transfer, *DeepMS^2^Reasoner* produces interpretable fragmentation pathways and enables the representation of fragment ion structures that are inaccessible to fixed-maximum-depth, bond-cleavage-only models.

**Fig. 1:**
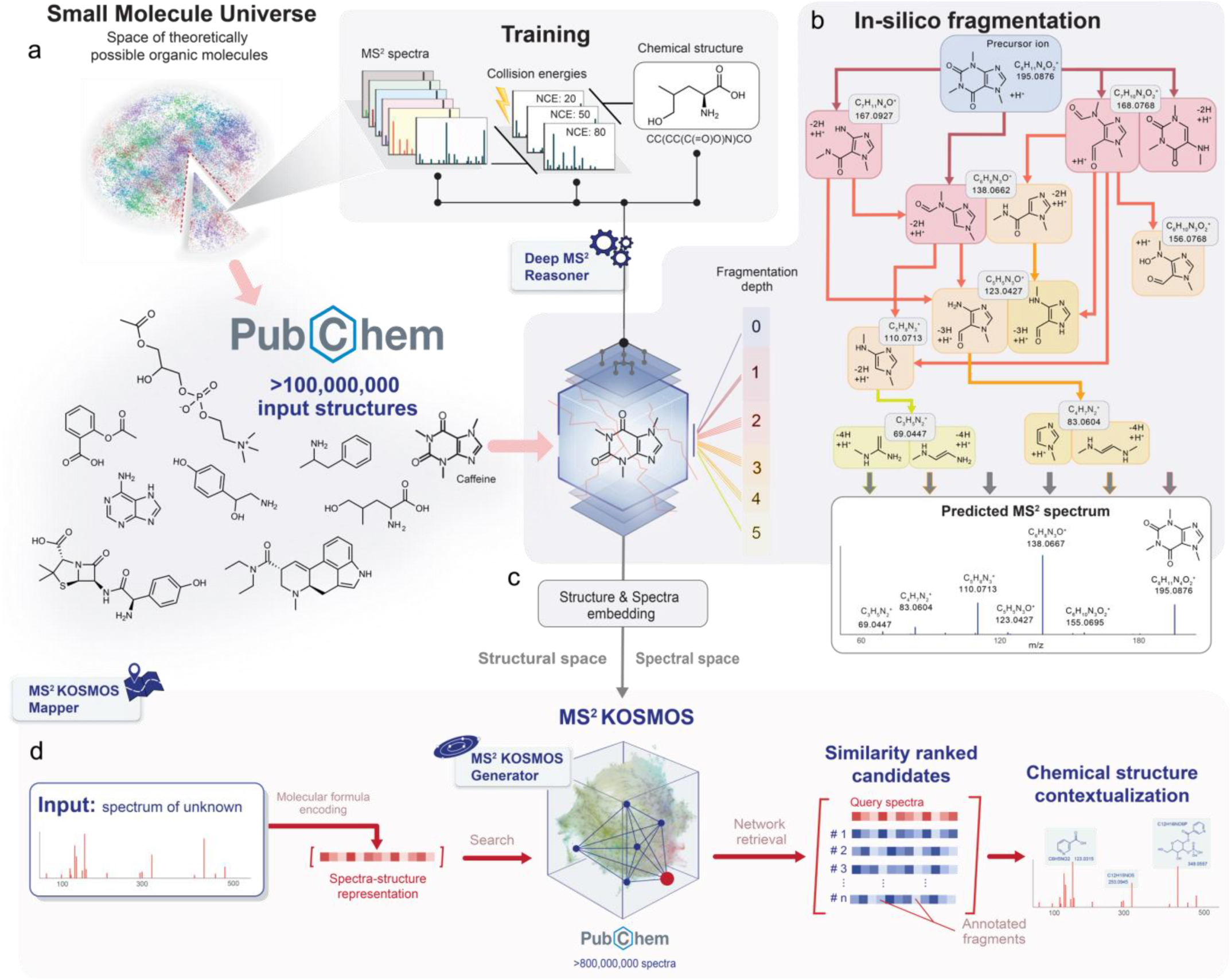
Overview of AIMe, a multi-agent neuro-symbolic AI framework for prediction, interpretation, exploration, and mapping of small-molecule MS data. AIMe comprises three modular agents, *DeepMS^2^Reasoner*, which predicts MS^2^ spectra and fragmentation pathways; MS^2^KOSMOSGenerator, which builds and indexes MS²KOSMOS, a searchable space of predicted MS^2^ spectra for known small organic molecules; and MS^2^KOSMOSMapper, which maps query spectra onto MS^2^KOSMOS. **a**, Schematic showing the *DeepMS^2^Reasoner* training set as a subset of the known chemical space, which in turn represents a subset of the universe of all theoretically possible small-molecule structures. The training set includes spectra acquired across multiple collision energies. **b**, *DeepMS^2^Reasoner* jointly predicts MS^2^ spectra and fragmentation pathways for any given precursor by modeling fragmentation as a sequence of symbolically enumerated chemically valid bond-breaking transitions with neural-assigned likelihoods. **c**, MS^2^KOSMOSGenerator orchestrates *DeepMS^2^Reasoner* to construct MS^2^KOSMOS, a searchable MS^2^ space of >800 million predicted MS^2^ spectra for >100 million small organic molecules in PubChem. **d**, *MS^2^KOSMOSMapper* maps query spectra onto the MS^2^KOSMOS to retrieve candidates and related structural neighborhoods taking into account the query’s exact experimental conditions.

Next, we constructed the MS^2^ Known Organic Small Molecule Space (MS^2^KOSMOS) as an efficiently queryable community resource comprising >800 million predicted MS^2^ spectra for >100 million compounds, covering essentially the entire set of known small organic molecules in PubChem.^30^ **(Fig. 1c)**. MS^2^KOSMOS was generated by another specialized AIMe agent that leverages *DeepMS^2^Reasoner* to orchestrate large-scale spectral prediction and organize the resulting spectra into a vast, searchable space. By indexing predicted fragments using symbolic fragment-formula representations, fast similarity queries across the entire MS^2^KOSMOS become feasible at a scale that would be intractable using raw spectra. Conceptually analogous to BLAST-based homology searches of nucleotide or protein sequences, AIMe’s *MS^2^KOSMOSMapper* maps any query spectrum to spectral neighborhoods within MS^2^KOSMOS, representing likely related chemical structures with analogous fragmentation patterns (**Fig. 1d**).

Finally, we demonstrate that AIMe can accelerate structure elucidation and repository-scale annotation. In a study of microbiota-dependent metabolites in mice, *MS^2^KOSMOSMapper* enabled rapid annotation of spectra likely representing known metabolites and prioritized plausible candidate substructures for unknowns, facilitating the discovery of unusual macrocyclic polyamines and other microbiota-dependent metabolites that were validated by synthesis and are also present in humans. At repository scale, applying *MS^2^KOSMOSMapper* to the GNPS database enabled putative annotation or identification of likely structural analogues for roughly a third of previously unannotated spectral clusters. Together, these applications position AIMe as a framework for scalable MS^2^-based structure elucidation, empowering researchers across disciplines to annotate unknown small molecules at scale.

## Results

### Fragmentation modeling: from structure to spectra

MS^2^ spectra are produced by fragmenting small molecules via collision with an inert gas at a defined kinetic energy (collision energy)^31^. Depending on the structural features of the parent molecule (the precursor), fragmentation can involve multiple consecutive bond breaks and rearrangements, producing a highly characteristic pattern of fragments with specific mass-to- charge ratios (*m/z*) and intensities that functions as a fingerprint of a compound’s molecular structure.^26,32,33^. At AIMe’s core is the *DeepMS^2^Reasoner*, a neuro-symbolic *AI* model that learns how the MS^2^ fragmentation process depends on molecular structure and collision energy for a given ionization mode. Starting from the precursor structure, *DeepMS^2^Reasoner* dynamically constructs a directed acyclic graph (DAG)^34^ representing fragmentation pathways, without imposing a predefined maximum depth. Symbolic chemical operators generate feasible fragmentation actions, while neural learning assigns their likelihoods, from which fragment intensities and the predicted MS^2^ spectrum are derived.

At each generated node (fragment) of the fragmentation DAG, *DeepMS^2^Reasoner* enumerates all permitted fragmentation actions, including halting (no further fragmentation), using three chemical operators; linear-bond cleavage, ring-bond cleavage, and excision **(Fig 2a)**. The bond cleavage operators model charge and hydrogen transfers between fragments, with the excision operator additionally modeling removal of internal substructures followed by reconnection of the retained boundary atoms^35^ **(Fig 2b)**.

**Fig. 2:**
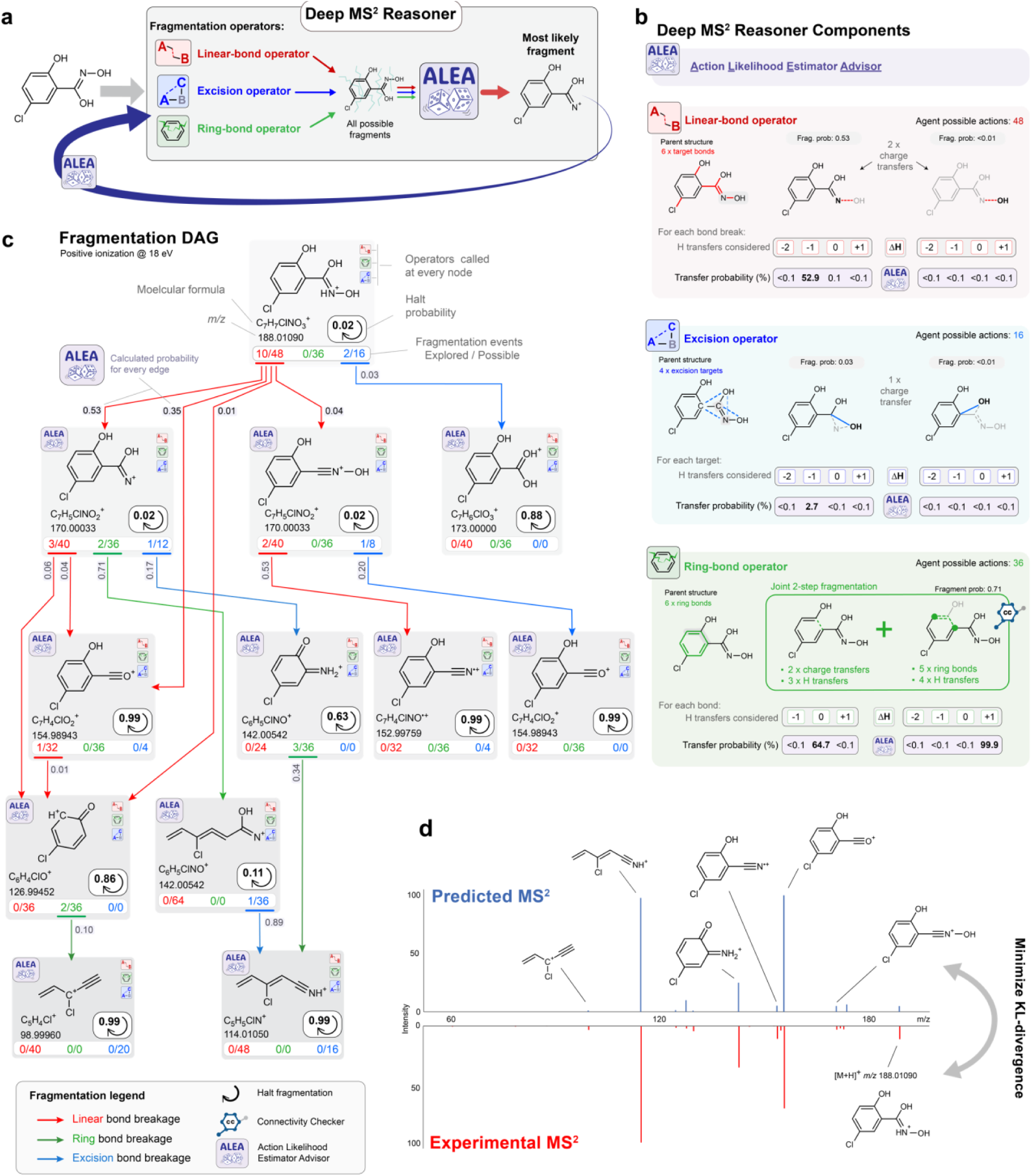
Fragmentation pathway generation and MS^2^ spectrum prediction in *DeepMS^2^Reasoner*. **a**, Overview of the roles of different components within *DeepMS^2^Reasoner*. ALEA, a structure-informed graph transformer, plays a central role by determining the relative likelihoods of all considered fragmentation outputs. **b,** Three different types of fragmentation mechanisms are considered by *DeepMS^2^Reasoner* in the form of chemical operators for (i) linear-bond cleavage, (ii) ring-bond cleavage, and (iii) excision, together with possible hydrogen transfer and respective learned probabilities by ALEA. Chemical feasibility checkers (e.g., connectivity and hydrogen balance) exclude invalid actions. **c,** Example DAG generated by *DeepMS^2^Reasoner* for a molecule under positive ionization at 18 eV. Nodes correspond to fragment ions annotated with molecular structure, formula, and *m/z*, while edges represent linear-bond cleavage (red), ring-bond cleavage (green), or excision events (blue). Note in the upper left example (C_7_H_5_ClNO_2_^+^ >> C_6_H_5_ClNO^+^), excision of the C-O moiety from the imide motif results in new C– N bond absent in the precursor ion. Color-coded annotations at the bottom of each node indicate, for each fragment, the number of possible bond-breaking and excision actions per fragmentation type and the number of actions actually expanded. Edge labels indicate transition probabilities assigned by ALEA. Loop arrows denote ALEA-calculated probability of halting fragmentation at a specific node, which contributes directly to the corresponding peak in the predicted spectrum. **d,** Experimental MS^2^ spectrum (bottom) and predicted MS^2^ spectrum (top), obtained by reasoning globally over all fragmentation pathways in the DAG and aggregating halting probabilities.

Conceptually, *DeepMS^2^Reasoner* implicitly searches the space of all possible fragmentation pathways, which increase exponentially with molecular size and complexity. However, guided by its Action Likelihood Estimator Advisor (ALEA), a structure-informed graph transformer^36^ that predicts the likelihood of candidate fragmentation actions, *DeepMS^2^Reasoner* typically expands only pathways with non-negligible likelihood, leaving unlikely branches unexpanded and thereby approximating the relative probabilities of different fragmentation events in the spectrometer.

To decide whether to expand a candidate action, *DeepMS^2^Reasoner* considers its global contribution to the spectrum, weighing ALEA’s action likelihood against the cumulative probability of reaching the current fragment, aggregated over all paths from the precursor. This decision is revisited dynamically: because a fragment can be reached by many routes, whenever a new path to a fragment or any of its ancestors is found, *DeepMS^2^Reasoner* updates the affected probabilities and reconsiders actions it had previously not expanded, refining the fragmentation DAG. ALEA plays a central role in guiding this symbolic pathway construction by ranking the large number of theoretically possible bond-breaking and excision actions according to their neural-assigned probabilities and identifying the most likely patterns of charge and hydrogen transfers between the resulting fragments **(Fig 2a,b)**.

*DeepMS^2^Reasoner* also tracks all atoms through each bond-breaking action, which is critical for the more complex fragmentation mechanisms defined by the excision and ring-breaking operators. Only fragments that satisfy symbolically enforced chemical feasibility constraints are retained as plausible and further expanded, ensuring interpretable, physically meaningful fragmentation pathways (see Methods). While chemical constraints are enforced at individual bond-breaking events, ALEA assigns a normalized probability distribution over the complete fragmentation action set, conditioned only on the current fragment state and experimental context, which are then aggregated to produce individual fragment intensities **(Fig 2c,d)**.

Training uses a Kullback-Leibler- (KL-)^37^ divergence-based loss that measures agreement between the predicted and experimentally observed spectra **(Fig. 2d)**, with additional regularization terms prioritizing simple linear-bond breaks over more complex actions. Memoization avoids redundant expansion by merging previously encountered fragments, allowing *DeepMS^2^Reasoner* to reason about exponentially many different pathways of fragmentation in time polynomial in the DAG size, taking advantage of the Markovian^38^ property of the fragmentation model. A direct consequence of this Markovian property is that the MS^2^ spectrum of a precursor can be expressed recursively as a convex combination of the spectra of its one-step fragments and its halting probability. This recursive decomposition naturally captures the observation that the fragmentation behavior of a molecular superstructure is highly informative about the fragmentation behavior of its constituent substructures. Together, these mechanisms enable *DeepMS^2^Reasoner* to efficiently explore the fragmentation space without being restricted to a fixed maximum fragmentation depth, while predicting chemically plausible fragmentation pathways and MS^2^ spectra. In addition, to accelerate learning, *DeepMS^2^Reasoner* is trained using a *curriculum learning strategy*^39^ that starts with simple molecules and gradually progresses to more complex structures. For simpler molecules, *DeepMS^2^Reasoner* explores a large number of fragmentation pathways, allowing ALEA to learn generalizable fragmentation patterns that in turn accelerate convergence and improve performance on larger, more complex compounds.

A central distinction from prior approaches is that *DeepMS^2^Reasoner* dynamically generates compound-specific fragmentation pathways without a predetermined maximum depth using an expanded set of fragmentation actions. In contrast, recent deep learning–based methods^19,20,40^, predict spectral intensities from fragment-level representations that are constructed independently from the learned intensity model, constrained to predetermined maximum depth fragmentation pathways, and a limited set of fragmentation actions. *DeepMS^2^Reasoner* instead predicts and explains spectral intensities at the level of individual fragmentation actions, dynamically constructing the fragmentation pathways step by step through symbolic chemical reasoning guided by ALEA’s neural likelihood estimates. Importantly, its action space includes excision, which creates product-ion bonds not present in the precursor and thus cannot be represented by bond-cleavage-only fragmentation models. Moreover, because *DeepMS^2^Reasoner* constructs pathways step by step, hydrogen transfer is represented as part of the evolving fragmentation state rather than as a post hoc scoring step, which is important because alternative hydrogen-shift outcomes generate distinct fragment states that can undergo different downstream fragmentation. Thus, subsequent fragmentation decisions are conditioned on the specific sequence of bond-cleavage, excision, and hydrogen-transfer events that produced the current fragment (**Fig 2c)**. Together, pathway construction without a predetermined fixed maximum depth and explicit consideration of excision and sequential hydrogen transfer events enable *DeepMS^2^Reasoner* to represent fragmentation sequences and product-ion connectivities that are systematically inaccessible to fixed maximum depth fragment enumeration approaches and bond-cleavage-only action spaces^19,20,40^.

### *DeepMS^2^Reasoner* significantly improves prediction accuracy

For training *DeepMS^2^Reasoner* we used the NIST 2020 dataset^41^, which was split into three parts, with 80% reserved for training, 10% for validation, and 10% for testing (see Methods). Evaluation of the trained model yielded predicted MS^2^ spectra with average cosine similarity scores to the experimental data of 0.83, with 50% of the predicted spectra having a cosine similarity greater than 0.89 **(Fig. 3a,b)**. This is comparable to, and surpasses, the minimum thresholds commonly used for annotation via library matching^42^. Other evaluated models achieved average cosine similarity scores between 0.61 and 0.75 for the same test dataset^20,40,43–45^ (see **Supplementary Table 1** for additional metrics). *DeepMS^2^Reasoner* performed best across the full molecular-weight range **(Fig. 3c)**, although performance of all models generally declined with molecular weight increase across all benchmarks (**Extended Data Figs. 1-3)**, likely owing to a greater number of possible fragmentation actions and sparser training coverage of larger-molecule structure space.

**Fig. 3:**
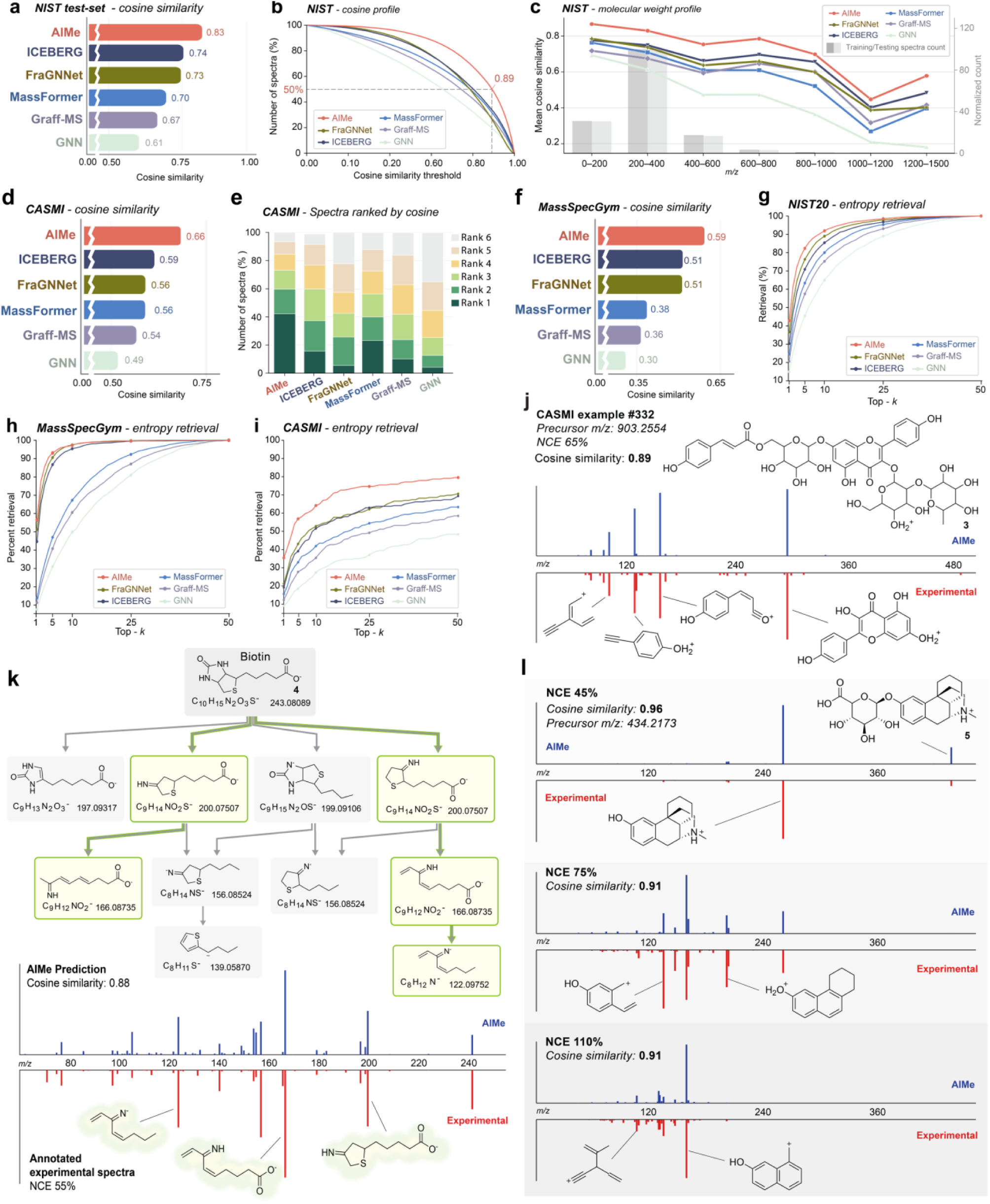
MS^2^ prediction by *DeepMS^2^Reasoner* outperforms existing approaches. a, Aggregate cosine similarity between experimental and predicted spectra for the NIST20 test set, comparing *DeepMS^2^Reasoner* with other models. All models were trained and tested with the same data splits, in both positive and negative modes (see Methods). **b,** Cosine similarity profile of all tested models, representing the number of spectra above or equal to any given cosine threshold. The 50% line denotes the median cosine similarity (0.89 for *DeepMS^2^Reasoner*). **c**, Performance of all models with increasing molecular weight. The total number of spectra within each bin in the training set is represented by gray bars. **d**, Cosine similarity of all predicted spectra for the CASMI2022 challenge dataset. **e,** Comparison of spectral prediction quality by different AI models for the CASMI set. Different colors in the stacked bars represent percentages of predicted spectra from a specific model that, when compared with the experimental data by cosine, are most similar, second-most similar etc. among the six evaluated models. **f**, Aggregate cosine similarity between experimental and predicted spectra for the MSG test set, comparing *DeepMS^2^Reasoner* with other models. **g**, NIST20 test set retrieval using entropy. The top 50 candidates from PubChem, ranked by Tanimoto similarity, were predicted for each model and compared against the corresponding experimental spectrum in the NIST20 test set, generating top-k retrieval of the correct matching structure. **h**, Using the MSG predefined test set, the top 50 candidates from PubChem were predicted, and retrieval was calculated for each model using entropy (additional performance with MSG-defined criteria can be found in Supplementary Table 6). **i**, CASMI test set retrieval using entropy to all isomers present in PubChem for each spectrum in the CASMI test set. **j**, Out-of-training-distribution example spectra from the CASMI dataset. **k**, *DeepMS^2^Reasoner*-generated fragmentation DAG for biotin, whose predicted spectrum achieved a cosine similarity of 0.88 (ICEBERG 0.39, FraGNNet 0.59). **l**, Experimental MS^2^ spectra (bottom) and predicted MS^2^ spectra (top) for a morphinan alkaloid glucuronide at 3 different collision energies.

Next, we tested AIMe on two other datasets that have been used extensively for evaluating the performance of MS^2^ prediction algorithms: the CASMI 2022 challenge dataset^46^ and the community-developed benchmark dataset, MassSpecGym (MSG)^47^. Both encompass a diverse range of organic molecules selected specifically for testing MS^2^ prediction models^46,47^. AIMe’s *DeepMS^2^Reasoner*, achieved the highest average cosine similarity on both test sets (CASMI: 0.67 MSG: 0.59, **Fig. 3d,f**), producing the closest predictions to the experimental spectra in >40% of cases, compared with the average cosine similarity runner-up ICEBERG^20^ in <20% of the cases **(Fig. 3e, Extended Data Figs. 2b,3b** and **Supplementary Tables 2,3)**.

Similarly, *DeepMS^2^Reasoner* outperformed all other evaluated tools across all benchmark datasets in retrieval of the correct structure from a set of predicted decoy spectra, using either the entropy **(Fig. 3g,h,i)** or cosine similarity metrics **(Extended Data Fig. 1-3** and **Supplementary Tables 4-6)**. The decoy sets were generated following previous examples^19,20,40^, and include spectra of the target molecule and up to 49 PubChem candidates selected within 10 ppm of the target mass, prioritized by Tanimoto similarity^48^. In addition, for the much smaller CASMI dataset, we designed more challenging, and perhaps more realistic, decoy sets that for each target compound include all of its isomers in PubChem. *DeepMS^2^Reasoner* achieved top-1, top-10, and top-50 accuracies of 35.6%, 64.4%, and 79.8%, respectively, consistently outperforming the runner-up, FraGNNet^40^, with >10% in top-1 and top- 10 retrievals **(Fig. 3i** **and Supplementary Table 5b)**.

*DeepMS^2^Reasoner*’s strong performance across all tested benchmarks and decoy-set comparisons (**Supplementary Tables 1-6**) validates its core architecture, in which pathway construction and spectral-intensity derivation are tightly coupled to capture multistep fragmentation cascades even in challenging chemical structures. For example, in the case of biotin, a cofactor commonly detected in metabolomic analyses **(Fig. 3k)**, *DeepMS^2^Reasoner* predicts the spectrum, matched to the experimental collision energy, with an overall cosine similarity of 0.88. Importantly, the most prominent fragment ion at *m/z* 166.0874 is derived from a multi-step fragmentation cascade, including loss of CONH from the ureido ring, opening of the tetrahydrothiophene ring, and elimination of H_2_S. Furthermore, the fragment at *m/z* 122.0975 cannot be explained by a single fragmentation event of the precursor and instead can be derived only via multi-step fragmentation. These deep fragmentation cascades are uncovered by *DeepMS^2^Reasoner* via exploration of the very large number of theoretically possible fragmentation pathways, including for larger and more complex molecules **(Fig. 3j, Extended Data Fig. 4** for additional examples).

The generalizable fragmentation patterns learned during training further position *DeepMS^2^Reasoner* to predict spectra with high fidelity across a range of collision energies **(Fig. 3l)**. The underlying fragmentation is modulated by varying collision energy, reflecting deeper and more extensive fragmentation cascades at higher energies^31,32^. This capability is particularly valuable in practice, where MS^2^ spectra are routinely acquired at multiple and often non-standardized collision energies across different instruments and thus energy-matched experimental references may not be available.

## Predicting mass spectra for the small-molecule universe

A major first goal in the process of identifying any unknown molecule from MS^2^ spectra is its chemical contextualization, which relies on identifying candidates and related molecules that share some of the unknown’s structural characteristics. This serves two purposes: (i) structural features of related molecules can provide inspiration for proposing candidate structures for the unknown, and (ii) the likely presence of specific structural features as well as the biochemical origin and/or biological properties of related molecules can assist with prioritization of unknowns for more detailed characterization. Given that the identities of most peaks in untargeted HPLC-MS analyses are usually unknown, such prioritization is highly desirable.

Inspired by genomic and proteomic annotation strategies, where gene or protein sequences are contextualized via homology search (e.g., BLAST), we envisioned that MS^2^ spectra could analogously be mapped onto a space organized by shared MS^2^ fragment formulas. As a first step toward developing such a tool for the chemical contextualization of MS^2^ spectra, AIMe’s *MS^2^KOSMOSGenerator* orchestrates *DeepMS^2^Reasoner* to predict mass spectra across the entirety of organic compounds with *m/z* <1000 in the PubChem database (see Methods for details), encompassing >105 million chemical structures representing the vast majority of known organic molecules from any source, predicting their mass spectra in positive and negative ionization modes, at four different collision energies, resulting in > 800 million predicted spectra.

These predictions are then organized into MS^2^KOSMOS, a fragment-formula-informed space in which molecules are embedded according to their shared predicted fragments (**Fig. 4a**). Each spectrum is represented in a discrete *Formula Coordinate System* derived from predicted fragmentation pathways, which spans roughly 30 million dimensions in positive-ion mode and 26 million in negative and is indexed by exact fragment formulas rather than *m/z* values, so that spectra with shared fragment formulas lie close together. This representation supports scalable vector-based similarity search and neighborhood-based reasoning for both structural and spectral queries, in the spirit of homology-based approaches such as BlastGraph^49^ or BlastAlign^50^.

**Fig. 4:**
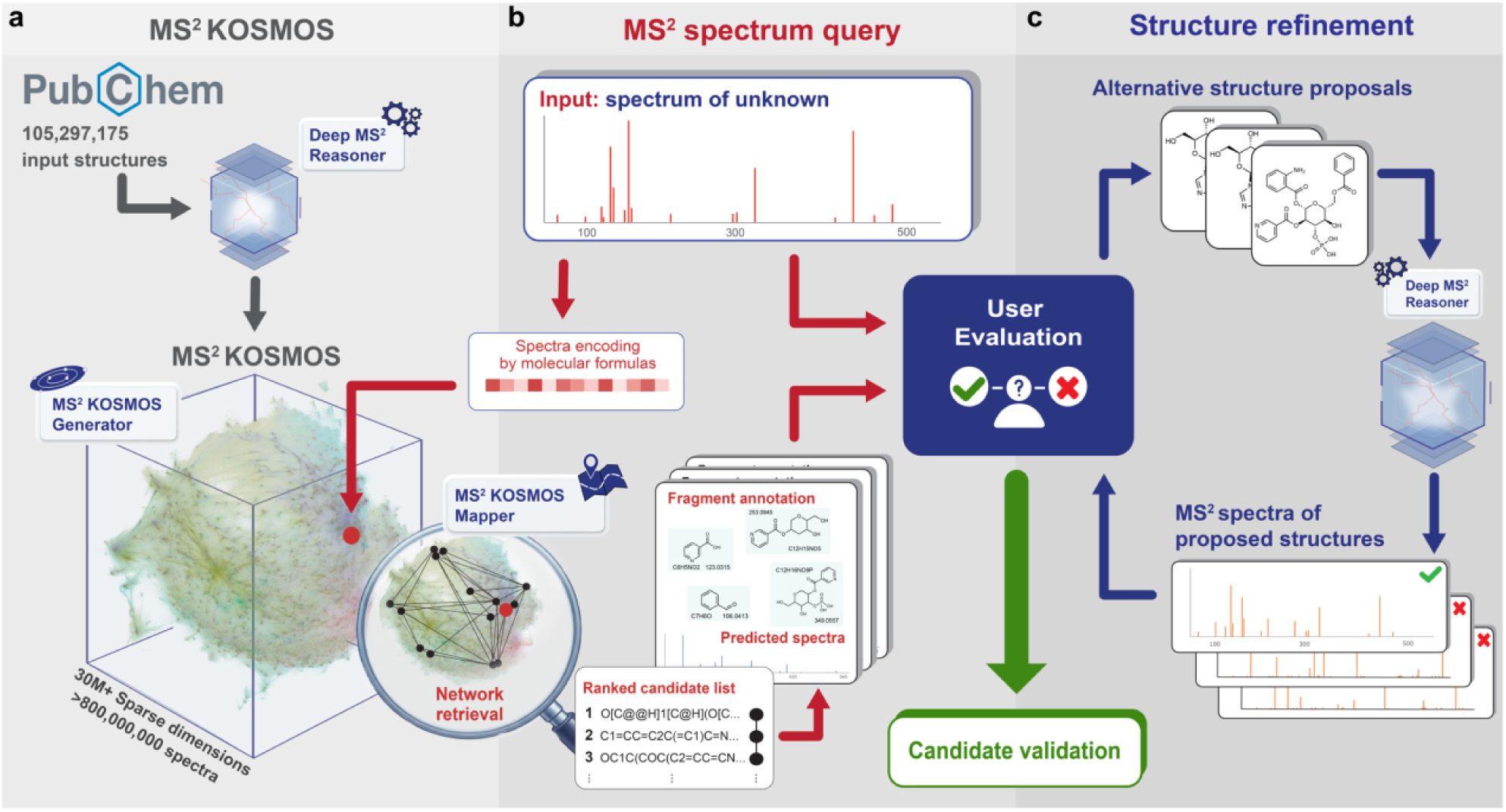
Contextualizing unknowns within MS^2^KOSMOS. **a**, *MS^2^KOSMOSGenerator* orchestrates *DeepMS^2^Reasoner* to predict MS^2^ spectra for 105.3 unique organic molecules with *m/z* <1,000 in PubChem. These predictions are organized into MS²KOSMOS, in which each predicted spectrum is encoded as a vector in a >30-million-dimensional fragment-formula space for positive-ion mode (∼26 million in negative), with each coordinate corresponding to a unique fragment or neutral-loss molecular formula across all structures. **b**, Experimental query spectra are encoded in the same fragment-formula embedding and matched against MS^2^KOSMOS using the inverted index to retrieve initial candidates, which *MS^2^KOSMOSMapper* then re-ranks using *DeepMS^2^Reasoner* spectra predicted under the query’s exact experimental conditions. Each candidate is accompanied by its predicted MS^2^ spectrum and a corresponding fragmentation DAG annotated with fragment formulas and putative chemical structures, enabling direct comparison with the query (see Supplementary Interface Guide). **c**, The retrieved candidates provide a basis for selecting structures for validation and serve as inspiration for proposing alternative structures. Proposed structures can be evaluated by predicting their MS^2^ spectra with *DeepMS^2^Reasoner*, enabling iterative refinement of structural hypotheses for novel molecules beyond the boundaries of existing chemical databases.

## Localizing unknowns in MS^2^KOSMOS

MS^2^KOSMOS can now be queried with any unknown spectrum, not only to identify realistic spectral and structural candidates but also, importantly, to define likely chemical neighborhoods in which matched fragments carry structural assignments shared with or related to those of the query spectrum **(Fig. 4b)**. Existing scoring functions, however, are imperfect proxies for chemical similarity and are limited when used as stand-alone metrics for spectral retrieval.^33,51–53^ This limitation becomes more acute as the search space expands from a few hundred thousand metabolites in a single library to more than 100 million compounds in MS^2^KOSMOS.

To address this challenge, we developed the *MS^2^KOSMOSMapper*, which contextualizes queries through a coarse-to-fine workflow. It first queries the fragment-formula inverted index, built by MS^2^KOSMOSGenerator, to retrieve an initial set of candidate molecules that share fragments with the query, scoring only molecules with at least one shared fragment rather than all of MS^2^KOSMOS. It then re-ranks these candidates using *DeepMS^2^Reasoner* spectra predicted under the query’s exact experimental conditions, scoring fine-grained, energy-dependent agreement. The result is a set of ranked candidates, which may span several disjoint structural neighborhoods in MS^2^KOSMOS, reflecting different aspects of the query. For each candidate spectrum, the results include a detailed fragmentation DAG in which every fragment is annotated with its mass, molecular formula and putative chemical structure (**Fig. 4b**).

Mapping query spectra onto MS^2^KOSMOS enables their contextualization across a wide range of use cases. For each query, *MS^2^KOSMOSMapper* outputs within seconds a ranked list of candidates supported by shared fragments that may suggest related structures or inform about shared structural features (**Fig. 4b**). If the query compound is present in PubChem, i.e., if it represents a known molecule, the correct structure is highly likely to be found among the top candidates, based on the retrieval performance of our model (**Supplementary Tables 4-6**). If the query compound is unknown, candidates from one or more spectral neighborhoods can provide priors for alternative structure proposals (**Fig. 4c**). The proposed structures can then be evaluated by predicting their MS^2^ spectra with *DeepMS^2^Reasoner* and comparing them to the query spectrum, as part of an iterative workflow that refines structural hypotheses and prioritizes candidates for further characterization, e.g., via synthesis of authentic samples for direct comparison and validation (**Fig. 4c**). This approach has the potential to dramatically accelerate the structure elucidation process and enable systematic annotation of the large numbers of unknowns typically detected in metabolomics.

## AIMe enables annotation of microbiota-dependent metabolites in mouse

To illustrate the potential of AIMe for compound discovery, we applied this framework to investigate metabolic differences between germ-free (GF) mice and mice with a replete gut microbiota (specific pathogen free (SPF) mice). Gut-microbiota-derived metabolites play a central role in animal physiology, including the development and maintenance of the immune system, regulation of fat metabolism, and anti-tumor responses^54–56^. Comparative HPLC-MS analysis of fecal samples from GF and SPF mice revealed several 1000 features that are significantly depleted or enriched more than 4-fold in GF mice relative to SPF mice, most of which could not be identified using available standards and reference libraries^41^. We then mapped MS^2^ spectra of the 111 most abundant unidentified microbiota-dependent features (**Supplementary Table 7**) onto MS^2^KOSMOS to retrieve spectral neighborhoods that would enable contextualized interpretation at both the fragment and structure levels (**Fig. 5a**). Roughly a third of the unidentified metabolites had close matches with MS^2^KOSMOS spectra suggesting that they represent known compounds or closely related isomers (for examples, see **Fig. 5b** and **Supplementary Fig. 1**). The remaining microbiota-dependent metabolites did not map to isomers with highly similar spectra in MS^2^KOSMOS, indicating that they likely represent yet undescribed metabolites. These unknowns included some of the most abundant microbiota- dependent metabolites in our dataset. Two examples are shown in **Fig. 5c**.

**Fig. 5:**
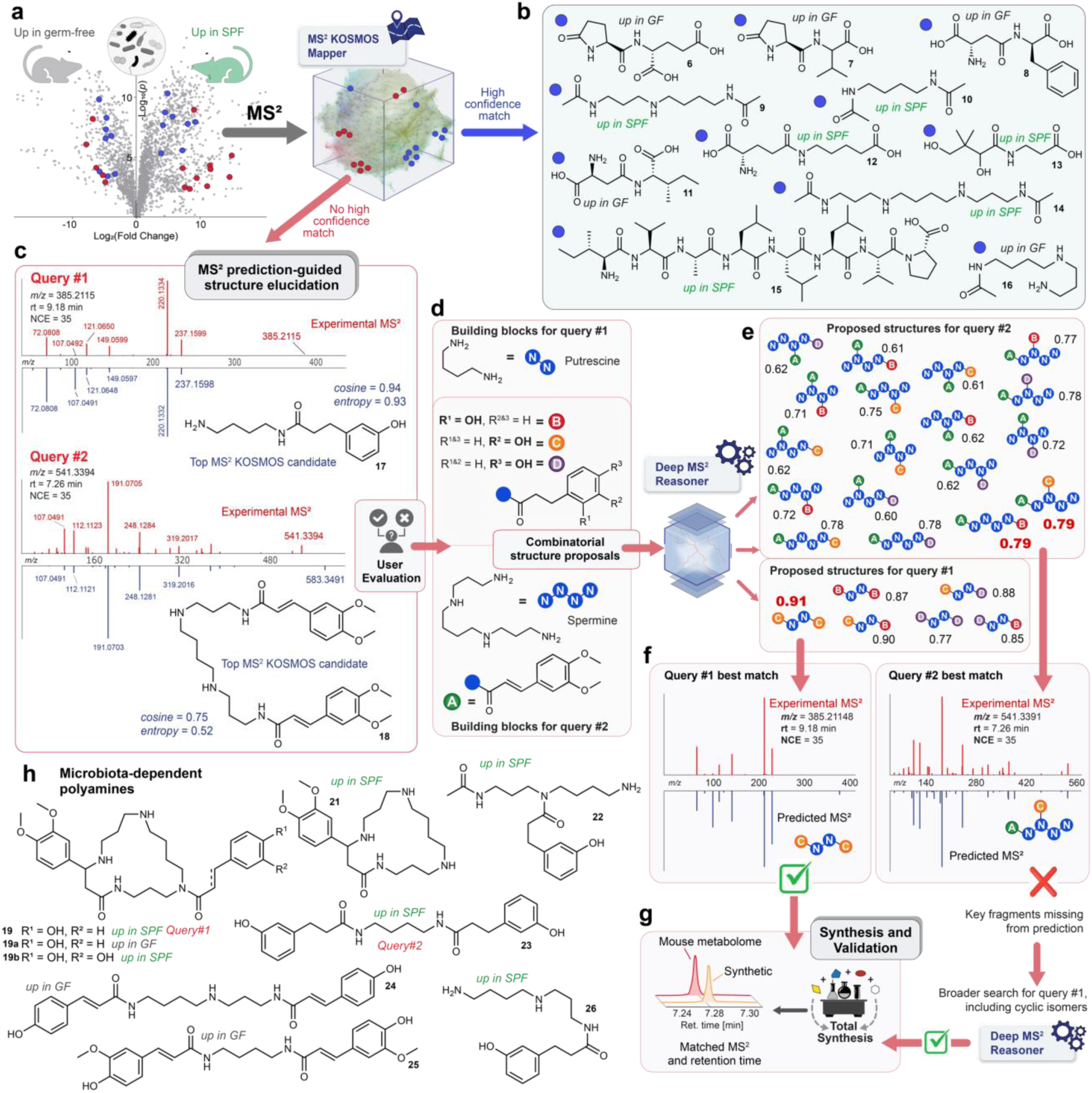
**AIMe-powered identification of gut-microbiota-dependent metabolites**. **a,** Unidentified microbiota dependent metabolites detected via comparative metabolomics of feces from germ-free (GF) and specific pathogen-free (SPF) mice are mapped onto spectral neighborhoods in MS^2^KOSMOS. **b,** A subset of metabolites had close matches with MS^2^KOSMOS spectra suggesting that they represent known compounds or related isomers. **c,d,** Many other microbiota-dependent metabolites did not map to isomers with highly similar MS^2^ spectra, indicating that they likely represent yet undescribed metabolites. Shown are two examples, query #1 (m/z 385.2115) and query #2 (*m/z* 541.3394). For these unknowns, analysis of the most similar MS^2^ spectra from MS^2^KOSMOS (**c**) suggested possible fragment structures and connectivities (**d**). **e**, Combinatorial assembly of inferred fragments allowed proposing arrays of potential candidate structures. Shown are a selection of most intuitive possibilities (6 for query#1 and 18 for query#2). For all proposed structures, MS^2^ spectra were predicted at query-matched collision energies and then compared with the query spectra. Numbers adjacent to the proposed structures reflect calculated cosine scores in positive ion mode (best matches highlighted in red). **f,** Experimental positive- ion MS^2^ spectra for queries #1 and #2 and predicted MS^2^ spectra for the best-matching proposed structures. Closer inspection of the predicted spectra for query#2 revealed the lack of key fragments in all predicted candidates, prompting an expanded search. **g,** Validation of proposed structures via synthesis and subsequent direct comparison of retention times and MS^2^ spectra (see Extended Data Fig. 5-8). **h,** Previously undescribed microbiota-dependent polyamine metabolites related to queries#1 and #2 identified via MS^2^ prediction-guided structure elucidation.

Toward elucidating the structures of these unknowns, we aimed to first develop viable hypotheses about the structures of the fragments in their MS^2^ spectra, based on examination of the predicted MS^2^ spectra and DAGs of their respective best matches from MS^2^KOSMOS (**Fig. 5d**). Here we took advantage of the chemically interpretable DAGs that AIMe provides for each predicted spectrum, which include explicit structures for all fragments and provide a rationale for their sequential origin from the parent structure (for DAGs of the spectra shown in **Fig. 5c**., see **Supplementary User Interface Guide**).

Next, we generated arrays of potential candidates for each query via combinatorial assembly of building blocks derived from the inferred fragment structures, e.g., 6 candidates in the case of query#1, a putative putrescine derivative, and 18 structures in the case of query#2, putatively assigned as a spermine derivative (**Fig. 5e,f)**. For all proposed structures, MS^2^ spectra were predicted at query-matched collision energies and then compared with the query spectra (see **Extended Data Figs. 5-7**) to prioritize candidates for further validation (**Fig. 5g**). In case of query#2, even the best-matching candidate spectra (cosine 0.79) lacked several key fragments prominent in the experimental spectra, which led us to expand our search to consider a wider range of candidates, including cyclized variants, for which AIMe predicted production of the fragments that were missing in the spectra the initial 18 candidates (**Extended Data Fig. 6-8**). We then synthesized reference samples for the best matching candidates for both queries (see **Supplementary Information**). Retention times and MS^2^ spectra of the highest ranked (by cosine) candidate for query#1 (**Extended Data Fig. 5**) and the cyclic candidate for query#2, a highly unusual macrocyclic spermine derivative that was predicted to produce the key fragments absent from the MS^2^ spectra of other high-ranking candidates, were identical to those of the corresponding query metabolites, confirming their chemical identities (**Extended Data Figs. 6- 8**).

Identification of the macrocyclic spermine and putrescine derivatives representing queries#1 and #2 enabled putative assignments for additional previously undescribed polyamine - derivatives whose MS^2^ spectra appeared to be closely related (**Fig. 5h** and **Supplementary Fig. 2**). These include metabolites enriched or exclusively present in SPF mice, which could be directly microbiota-derived, as well as polyamines that are enriched in GF mice, e.g., because of lack of microbial degradation or altered host metabolism, highlighting the multi-layered effects of microbiota deletion on host metabolic networks. Search for the MS^2^ spectra across the GNPS database indicates that many of the identified microbiota-dependent metabolites, including the macrocyclic spermine derivatives, are also present in samples of human origin (**Extended Data Fig. 9**).

The structures of the identified macrocyclic polyamine derivatives are unlike any previously identified from mouse or human, but are reminiscent of a family of macrocyclic alkaloids of plant and microbial origin that exhibit diverse bioactivities^57^. In the context of human biology, polyamines are increasingly recognized to sit at the nexus of diet, microbiota, epithelial biology, and immune signaling, and thus identification of new families of microbiota-dependent polyamines may contribute to understanding the multifaceted physiological roles of the microbiota. From a metabolomics perspective, our results show that combining search across MS^2^KOSMOS with combinatorial generation of candidate structures followed by MS^2^ prediction-guided ranking of candidates provides a powerful resource for structure elucidation, even in cases of unusual or unexpected scaffolds, such as the macrocyclic spermine derivatives.

## AIMe supports repository-scale annotation

To explore the potential of utilizing AIMe for spectral annotation at repository-scale, we employed a large previously published MS^2^ dataset derived from the GNPS repository^42,58^, a critical community resource that enables researchers to host, share, organize and analyze MS^2^ data. Reflecting the typically very low annotation level of metabolomics datasets, the chemical structures of the vast majority of the 521 million experimental MS^2^ spectra hosted at GNPS are unknown. Previous efforts to increase annotation levels organized these MS^2^ data into 8,543,020 spectral clusters, with each cluster representing a group of related spectra across several source datasets **(Fig 6a)**.

**Fig. 6:**
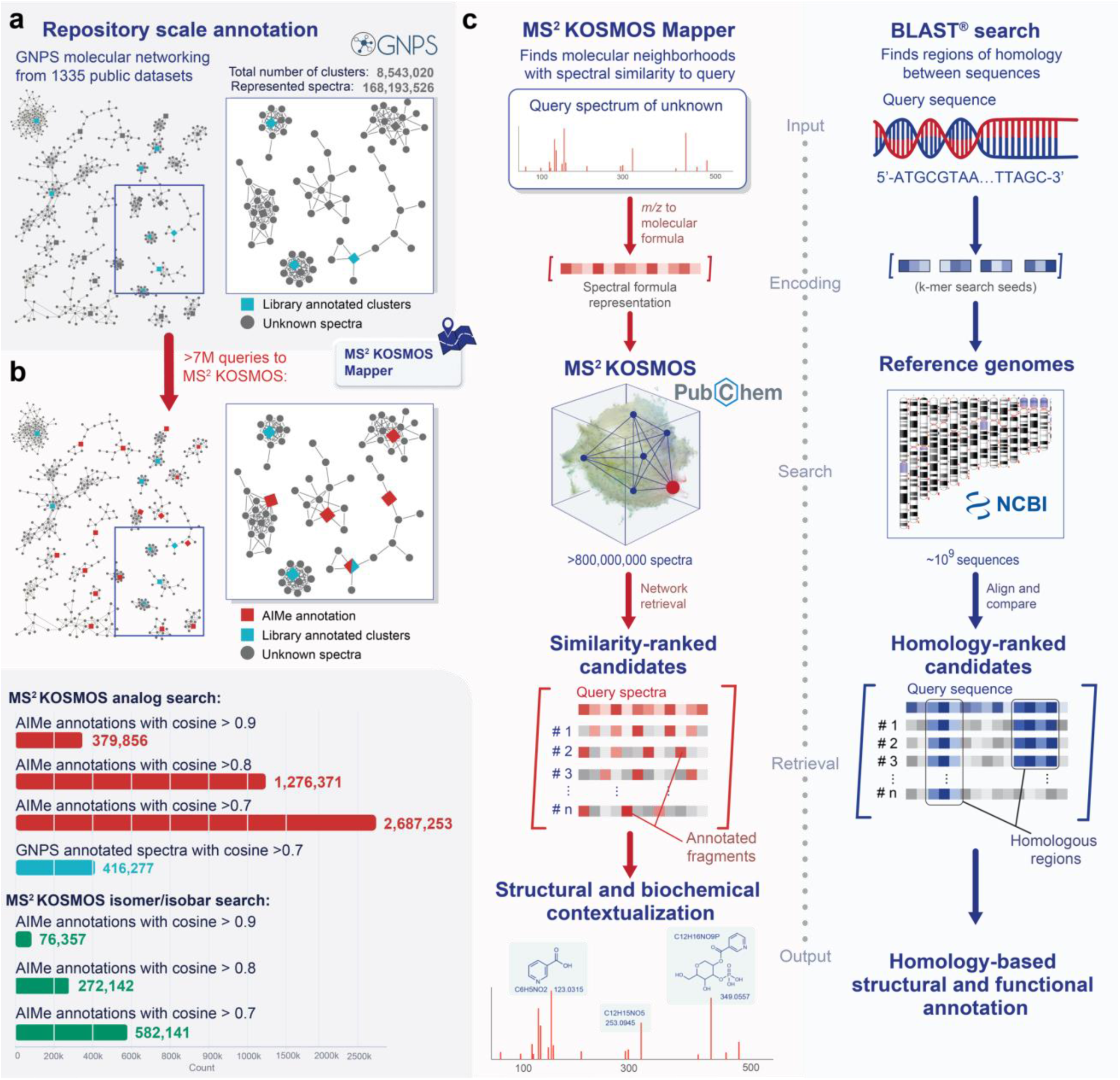
**AIMe supports repository-scale annotation**. **a,** GNPS derived network from a previous study^42^ consisting of 8,543,020 clusters, of which 7,144,480 were searched after filtering. **b**, AIMe retrieved 1,276,371 annotations, considering top-ranked candidates with a spectral similarity to the query of 0.8 and above. For similarity matches of 0.7 and above, 2,687,253 annotations, of which 379,856 were above 0.9 cosine similarity to the experimental spectra. Restricting MS^2^KOSMOS search space to candidates within 20 ppm or 0.01 Da of the query precursor, a stricter criterion than used previously^42^, yields subsets of annotations representing likely isomers. **c**, Bioinformatic tools for sequence searching such as BLAST inspired AIMe’s design for mapping of experimental MS^2^ spectra onto the PubChem-derived MS^2^KOSMOS, providing chemical contextualization as a basis for structure elucidation. Analogous to retrieving nucleotide or protein sequences based on homology, the *MS^2^KOSMOSMapper* ranks candidate structures from MS^2^KOSMOS based on similarity of molecular formula-encoded spectral embeddings.

To determine the extent to which querying MS^2^KOSMOS could further increase annotation levels, we filtered this dataset to eliminate multiply charged species and large molecules above *m/z* 1000, resulting in a test set of 7,144,480 consensus spectra. We then used *MS^2^KOSMOSMapper* to retrieve spectral neighborhoods for each of these consensus spectra in MS^2^KOSMOS. For roughly a third (∼2.69 million) of the queried experimental spectra we obtained annotations supported by a cosine similarity above 0.7 to predicted spectra of known compounds in PubChem, compared to 416,277 previous annotations at cosine 0.7^42^. At stricter cosine similarity above 0.8, querying MS^2^KOSMOS added approximately 1.27 million annotations for the consensus spectra, representing a total of >26 million previously unannotated spectra in GNPS. Restricting candidates to within 20 ppm or 0.01 Da of the query precursor mass yielded 272,142 annotations enriched for likely isomers **(Fig 6b).** Detailed procedures for linking GNPS cluster representatives to raw spectra and searching MS^2^KOSMOS are provided in the Supplementary AI methods.

These results demonstrate that AIMe provides a scalable route to transforming the vast and rapidly expanding space of publicly available but unannotated MS^2^ spectra into interpretable outputs with meaningfully annotated DAGs available for each retrieved candidate, facilitating the interpretation and contextualization of MS^2^ spectra at the scale of large metabolomics datasets.

## Discussion

The majority of small organic molecules detected across biology remain unannotated, primarily because of their essentially infinite structural diversity, only a small fraction of which is covered by experimental reference libraries. Consequently, metabolomic data interpretation and follow-up typically focus on known metabolites, whereas potentially important but unidentified molecules are not pursued because elucidation of their structures would require specialist expertise and often prohibitive manual effort.

AIMe extends MS^2^ spectral annotation beyond searches of experimental libraries toward fragment-informed molecular neighborhood retrieval across most of known structure space as well as user-specified sets of hypothetical structures. At the core of this framework lies *DeepMS^2^Reasoner*, which bidirectionally couples symbolic chemical reasoning with structure-informed neural learning to model MS^2^ fragmentation. Learned probabilities guide the construction of chemically plausible sequences of bond-breaking and excision events, while the resulting fragmentation pathways provide spectral feedback that refines those probabilities. This coupling, incorporating a rich set of fragmentation actions, advances spectral prediction capabilities beyond those of other models **(Fig. 3)** and enables the joint prediction of spectra and fragmentation DAGs without a predetermined fixed maximum depth, across collision energies. Importantly, these outputs not only complement the sparse coverage of experimental libraries but also provide putative mechanistic rationalizations for the fragmentation events leading to every node in the fully annotated DAG.

Inspired by genomic and proteomic annotation workflows, in which novel genes or proteins are functionally characterized by sequence homology searches, we envisioned that MS^2^ spectra could be interpreted against a chemically meaningful space of shared fragments, analogous to BLAST homology searches (**Fig. 6c**). Toward this goal, we predicted MS^2^ spectra for the entirety of small organic molecules in PubChem, representing most of the known organic small-molecule universe, and organized these predictions into MS^2^KOSMOS as an efficiently searchable community resource. By indexing fragments through symbolic fragment-formula representations MS^2^KOSMOS makes similarity searches across this vast spectral space computationally feasible. *MS^2^KOSMOSMapper* uses this index to map any experimental MS^2^ spectrum onto MS^2^KOSMOS and retrieve molecular neighborhoods within seconds, with shared putative fragment structures supporting candidate ranking and structural-neighborhood discovery. An important distinction is that gene or protein sequence searches operate on database entries represented using a linear, well-defined alphabet and can exploit exact or near-exact *k-mer* matches, whereas the candidate search space for small molecules is combinatorial and embedded in a vast, extremely high-dimensional chemical space. Thus, while a BLAST match provides a deterministic entry point into local homology, an MS^2^ match is chemically informative but not definitive: an individual fragment may arise from different fragmentation pathways and parent structures, occur at varying intensities, and correspond to many chemically plausible substructures. Moreover, the same structure can produce different spectra depending on the fragmentation conditions^31,59–61^.

AIMe addresses these challenges by combining high-fidelity MS^2^ spectrum prediction and interpretable fragmentation pathways with rapid, fragment-informed molecular-neighborhood mapping across MS^2^KOSMOS, shifting the objective from identifying a single best-matching candidate to mapping unknowns into structurally similar molecular neighborhoods within MS^2^KOSMOS. We demonstrated the utility of this approach with the identification of previously unreported microbiota-dependent metabolites in mice. For each query metabolite, mapping onto MS^2^KOSMOS revealed sets of previously described compounds that appeared structurally related through matched fragments, enabling the proposal of sets of candidate structures (**Fig. 5c,d**). Predicting MS^2^ spectra for the proposed structures allowed to prioritize among the many candidates for validation via chemical synthesis **(Fig. 5f,g)**, which revealed previously unreported families of polyamines, including highly unusual macrocyclic structures that nonetheless represent some of the largest peaks in our mouse metabolomics datasets and are also present in samples of human origin. This example demonstrates that even for metabolite classes such as polyamines, which are generally considered well annotated^62,63^, chemical characterization remains highly incomplete and that interactive use of AIMe to map unknowns, develop structural hypotheses, and predict MS^2^ spectra of putative matches, can greatly accelerate structure elucidation. The metabolite identification challenge further demands approaches that reach beyond a single molecule, test set, or study and can complement community efforts to improve current annotation rates. Application of AIMe to MS^2^ spectra deposited at the GNPS repository enabled putative annotation of a substantial fraction of the queried spectra, significantly increasing annotation levels at GNPS and demonstrating that AIMe supports small-molecule contextualization at repository scale.

*DeepMS^2^Reasoner* and other spectrum-prediction models remain constrained by limited chemical diversity and uneven molecular-weight representation in their training data^47^. Larger molecules often fragment through more complex and diverse pathways, and their sparse representation in training data likely contributes to the decline in performance observed with increasing molecular weight across the evaluated models (**Fig. 3c**). More chemically diverse, higher-quality, and better-curated training spectra should therefore improve prediction accuracy. Evaluation also remains sensitive to the choice of spectral comparison metric (**Supplementary Tables 1-6**), as existing metrics exhibit structure-dependent biases and emphasize different spectral properties^59,60,64,65^. AIMe’s Formula Coordinate System partially mitigates this by enabling efficient comparison of spectra in a shared fragment-formula space, but further gains will likely require closer integration of fragmentation-aware similarity functions with retrieval objectives.

Without additional orthogonal information, AIMe outputs correspond most closely to level 3 annotations under the Metabolomics Standards Initiative framework^66^, providing a foundation for validation, as demonstrated in the mouse microbiota-dependent structural elucidation. Nonetheless, the interpretable framework for prioritizing candidates, organizing spectral neighborhoods, and generating mechanistic hypotheses when direct matches are unavailable provides valuable information beyond a confidence score, establishing AI-enabled Computer-Assisted Structural Elucidation^26^ as a practical approach. Because AIMe provides explicit fragment structures within a detailed fragmentation DAG, it could also serve as a testbed for MS^3^ and MS^n^ experiments. Conversely, including MS^3^ and MS^n^ data in model training sets may improve spectral prediction accuracy.

AIMe integrates specialized reasoning agents that combine symbolic, chemistry-aware operators, and validation modules with neural learning to (i) predict MS^2^ spectra and interpretable fragmentation pathways and (ii) map experimental spectra to contextualize unknowns, prioritize candidate structures, and guide structure elucidation. Future versions of AIMe could augment its specialized agents with language-model-based and multimodal reasoning capabilities, enabling integration of orthogonal sources of evidence, including chromatographic retention times, NMR spectra, biochemical knowledge bases, scientific literature, and experimental metadata. More broadly, AIMe illustrates how neural learning, symbolic reasoning, and explicit chemical domain knowledge can be integrated within a unified framework to accelerate discovery across vast, sparsely explored scientific spaces, with a wide range of applications in biology, medicine, and environmental sciences.

## Methods

### AIMe: a multi-agent framework, combining neural learning with symbolic reasoning and search over chemical space

AI Molecule Explorer (AIMe) is a multi-agent neuro-symbolic AI^28^ framework. AIMe couples three goal-directed agents around a shared resource, MS^2^KOSMOS: DeepMS^2^Reasoner, a forward fragmentation agent that predicts MS^2^ spectra by combining neural learning with symbolic and combinatorial reasoning over sequential fragmentation events in chemical space; *MS^2^KOSMOSGenerator*, a generation-and-curation agent that builds and indexes MS^2^KOSMOS by orchestrating *DeepMS^2^Reasoner*; and *MS^2^KOSMOSMapper*, an on-demand query-mapping agent that searches MS^2^KOSMOS to contextualize unknown spectra.

A central design principle of AIMe is the separation of large-scale, amortized pre-computation from on-demand, query-specific inference. Expensive neuro-symbolic reasoning over molecular structure and fragmentation is performed once at scale, producing predicted spectra, fragment annotations, fragmentation pathways, and an inverted index over fragment formulas. At query time, *MS^2^KOSMOSMapper* encodes experimental spectra into MS^e^KOSMOS’ Formula Coordinate System and uses this index to search MS^2^KOSMOS efficiently. Initial candidate rankings are then refined by targeted *DeepMS^2^Reasoner* predictions under the query-specific experimental conditions and contextualized into interpretable structural neighborhoods.

In this setting, the forward problem is to predict the fragmentation pathway and resulting MS^2^ spectrum for a molecule under a specified experimental configuration. AIMe approaches the inverse problem through two tractable tasks: constructing MS^2^KOSMOS as an efficiently searchable space of predicted spectra, and retrieving and re-ranking candidates for an experimental query. AIMe therefore couples forward neuro-symbolic fragmentation reasoning with inverse, query-driven retrieval over MS^2^KOSMOS. Its output is not only a ranked candidate list, but a fragment-resolved molecular neighborhood in which shared peaks can be traced to predicted fragment formulas, structures and the fragmentation pathways that generate them. Results are returned in seconds. Formal notation, problem definitions and complete agent specifications are provided in Supplementary AI Methods.

### Deep MS^2^ Reasoner: Joint fragmentation-pathway and spectrum generation

At the core of AIMe is the Deep MS^2^ Reasoner (*DeepMS^2^Reasoner*), a structure-informed neuro-symbolic generative model that reasons forward over an implicitly exponential space of bond dissociations to construct fragmentation pathways and derive the corresponding MS^2^ spectrum. The spectrum is obtained by propagating probability mass across the pathway and reading out an intensity from a learned halting probability at each fragment.

A *molecule*, whether the precursor or an intermediate product ion, is represented as a feature-augmented heavy-atom graph, with atoms as nodes and bonds as edges. Hydrogen atoms are not represented as explicit graph nodes; instead, hydrogen counts, hydrogen shifts and charge-related bookkeeping are encoded as atom-level features. Each fragmentation action deterministically updates the molecular graph and its chemical state. *DeepMS^2^Reasoner* considers three structural fragmentation operators: linear-bond cleavage, ring-bond cleavage and excision. Linear-bond cleavage separates a fragment into two complementary pieces, with the two possible charge assignments represented as distinct actions. Ring-bond cleavage opens a ring without immediately detaching atoms and may therefore proceed through multiple consecutive actions. Excision removes an internal substructure as a neutral loss and reconnects the retained boundary atoms, allowing the model to represent product-ion bonds that were not present in the precursor. Because both boundary atoms remain in the retained fragment, which carries the charge, excision does not branch into two charge assignments. Charge localization and hydrogen transfers are represented as part of the evolving fragment state, so subsequent fragmentation decisions are conditioned on the specific state produced by earlier actions. Exact action encodings, hydrogen-transfer states and operator-specific constraints are provided in the Supplementary Information. Fragmentation is formulated as a Markov process^38^ over molecular graphs: each state is a chemically annotated fragment, and each transition is a fragmentation action with a learned probability. *DeepMS^2^Reasoner* does not predict fragment intensities directly. Instead, fragment intensities emerge from transition and halting probabilities through probability-mass propagation, so that the model learns the fragmentation process rather than regressing peak intensities conditioned on the precursor.

The Action Likelihood Estimator Advisor (ALEA), a structure-informed graph transformer^36^, scores every enumerated fragmentation action, including halting, as a single probability distribution conditioned on the current fragment state and experimental configuration. Scoring halting alongside fragmentation actions lets the model balance continuation against termination using the same information. *DeepMS^2^Reasoner* implements pathway construction and generation as a coordinated, event-driven exploration over active fragment states using deterministic symbolic chemical operators and feasibility checkers together with ALEA. Fragmentation pathway generation proceeds by repeatedly expanding actions under their accumulated probability mass, incrementally constructing a probabilistic fragmentation directed acyclic graph (DAG). For a fragment (F), a non-halting action (a) is extended only when its contribution to the overall spectrum exceeds a predefined threshold, *p_in_* (*F*) × *p*( *a* ∣ *F*, ℰ) ≥ *ε* where *p_in_* (*F*) is the accumulated probability of reaching (F) through all constructed pathways and *p*( *a* ∣ *F*, ℰ) is ALEA’s action likelihood under experimental conditions ℰ . This mass-based selection leaves low-probability branches unexpanded while permitting fragmentation cascades without a predetermined maximum depth when supported by sufficient probability mass.

A ring-bond break can produce a transient ring-opening intermediate in which one or more bonds have been broken but no atoms have yet been detached. During such intermediate states, halting and excision are disabled, and ring opening is constrained to proceed through actions that make progress toward separating the initially selected parts of the ring. These actions are re-enabled once a fragmentation step detaches a piece and produces a product ion. This prevents intensity from being assigned to a ring-opening state that has not yet formed a fragment and ensures that ring opening proceeds as a coherent sequence. The exact charge- commitment, connectivity and action-enumeration rules are described in Supplementary Information.

For each selected non-halting action, the child fragment is constructed deterministically and subjected to hard symbolic chemical-feasibility constraints, including hydrogen-transfer constraints. Valid actions are added as directed transitions. Canonical graph hashing over fragment connectivity and chemical state merges previously encountered fragments, allowing distinct pathways that reach the same state to share a single DAG node. When additional probability mass later reaches an existing node, that fragment is reactivated and its outgoing actions are reconsidered under the updated incoming mass. Actions that were previously below the expansion threshold may therefore be extended later, so the DAG is refined as probability mass accumulates rather than in a single pass. The complete queue-based construction and reactivation procedure is given in Supplementary AI Methods, Algorithm 1.

An action that fails the feasibility checks is not added to the DAG, and its probability mass is not redistributed to the surviving actions. Instead, it is retained as unexplained mass, making probability placed on chemically invalid fragmentation visible to the training objective.

Together, these steps couple ALEA’s learned action selection, deterministic chemical operators, hard chemical constraints, canonical state merging and probability-mass–based expansion, allowing DeepMS^2^Reasoner to model chemically plausible fragmentation and predict spectra within a unified reasoning framework while explicitly exploring only a small fraction of the combinatorial pathway space and pruning highly unlikely regions.

### Deep MS^2^ Reasoner Training

*DeepMS^2^Reasoner* is trained by optimizing ALEA’s parameters using the same probabilistic fragmentation-pathway construction procedure employed during inference, supplemented during training with sampling of residual fragmentation trajectories as described below and in Supplementary Information. This creates a bidirectional coupling between symbolic DAG construction and neural learning: ALEA assigns action and halting probabilities that guide pathway expansion and determine path weights within the fragmentation DAG, while the resulting DAG and sampled residual trajectories define the predicted spectrum, fragment intensities, and spectral loss through which ALEA is updated. Within DeepMS^2^Reasoner’s neuro-symbolic architecture, ALEA is the only learned neural-network component. All other components, including action enumeration, chemical-feasibility checks, fragment-graph edits, canonical state merging, and threshold-based expansion, are implemented through deterministic symbolic operators and constraints. Neural learning therefore shapes fragmentation exclusively through ALEA’s action and halting probabilities, which guide DAG construction and are, in turn, learned from the resulting spectral error.

ALEA is implemented as a ***graph-based transformer (Graphormer***^36^) that operates directly on a feature-augmented molecular graph of the current fragment state. Node- and edge-level features encode local chemical properties including atom identity, bond type, ring context, hydrogen bookkeeping together with global experimental metadata including collision energy, instrument, adduct. Learned structural bias terms derived from pairwise graph features are incorporated into the self-attention mechanism^67^, enabling ALEA to reason over both local chemical environments and longer-range structural dependencies within the fragment. Action-specific prediction heads produce logits for linear-bond cleavage, ring opening, excision, and halting. Complete feature definitions, architectural specifications, and decoder inputs are provided in Supplementary AI Methods.

For each training example, consisting of a molecular structure, experimental configuration, and observed MS^2^ spectrum, *DeepMS^2^Reasoner* executes the threshold-based fragmentation- pathway construction procedure described above. Predicted fragment contributions are aligned to retained experimental peaks within mass tolerance. Low-intensity experimental peaks removed during preprocessing are aggregated into an additional *other* target bin. Predicted fragments that do not match a retained experimental peak, together with probability assigned to chemically invalid actions, are assigned to the same bin. Invalid actions are not inserted into the fragmentation DAG, and their probability is not redistributed among valid actions.

Actions whose probability-mass contribution falls below the expansion threshold are not included in the realized DAG. Although each omitted action carries little mass individually, their combined residual probability can be non-negligible. During training, this residual mass is accounted for by sampling additional fragmentation trajectories from the unexpanded action distribution. Each trajectory is continued under ALEA’s local action probabilities until halting or an invalid action, and is weighted by its share of the residual probability mass. These sampled trajectories are used only during training and do not alter the threshold-based inference procedure. Exact probability contributions from the realized DAG and weighted contributions from the sampled residual trajectories are aggregated into the aligned predicted spectrum, which is normalized before loss computation.

The model is trained using a ***KL-divergence–based loss***^37^ that measures agreement between the normalized predicted and experimentally observed spectra. The loss includes additional regularization terms that penalize the expected use of ring-opening and excision actions, softly favoring simpler linear-bond explanations when multiple fragmentation mechanisms account for the same observed peaks. The complete objective, regularization weights, spectral alignment procedure, and residual-trajectory estimator are provided in Supplementary Information.

Conditioned on the realized fragmentation DAG and sampled residual trajectories, gradients with respect to ALEA’s local action probabilities could be obtained by automatic differentiation through the complete probability-propagation computation. In our implementation, the initial derivatives of the spectral and regularization losses with respect to the local log action probabilities are instead computed analytically, which we found to be faster and more numerically stable. These derivatives are supplied as upstream gradients, and automatic differentiation then propagates them through ALEA’s action softmax, action-specific prediction heads, and graph-transformer encoder to its learnable parameters. The discrete pathway topology, chemical-feasibility decisions, canonical state merging, threshold comparisons, and sampled trajectory outcomes are treated as fixed during differentiation.

To stabilize optimization over the highly combinatorial fragmentation space, *DeepMS^2^Reasoner* is trained using a ***curriculum strategy***^39^ that gradually introduces higher-mass molecules. Training begins with lower-mass compounds, where action spaces are smaller and fragmentation pathways are shallower, allowing ALEA to learn transferable local fragmentation preferences. As training progresses and the learned action probabilities become more selective, larger and more complex molecules are introduced, enabling stable learning of deep, multi-step fragmentation behavior. Stochastic augmentation of experimental metadata is additionally used to improve robustness to instrument variability and metadata uncertainty.

### Training and test data processing

We processed the NIST20^41^ tandem mass-spectrometry dataset by retaining only MS^2^ spectra for protonated ([M+H]^+^) and deprotonated ([M-H]^-^) precursor ions, excluding compounds containing metals, and requiring each spectrum to have a valid molecular structure whose formula and InChIKey connectivity matched the NIST20 annotation. Spectra were split by full InChIKey into training, validation, and held-out test sets before peak processing, using an 80%, 10%, 10% split, respectively. Peaks were merged within 20 ppm mass tolerance and filtered to retain signals explainable by chemically feasible molecular subformulas within 5 ppm; spectra with no explainable peaks or large unexplained peaks were discarded. Molecular structures were canonicalized with RDKit^68^ tautomer enumeration. For final evaluation we removed held-out target molecules that were identical to any training molecule under Morgan fingerprint^69,70^ Tanimoto similarity^48^ of 1.0. CASMI and MassSpecGym^47^ were used only as external test benchmarks. CASMI^46^ spectra were standardized and filtered to retain formula-supported peaks before scoring. MassSpecGym [M+H]^+^ spectra were evaluated using the provided test spectra and annotations. For both benchmarks, targets identical to NIST training molecules by Morgan-fingerprint Tanimoto similarity of 1.0 were removed. For MassSpecGym, we additionally removed exact two- dimensional matches and test molecules with a Maximum Common Edge Subgraph (MCES^71^) distance below 10 to any NIST training molecule, applying the benchmark’s original structural- separation criterion relative to our NIST-trained model.

For inverse structure-identification evaluations, we followed the candidate-selection procedure used by FraGNNet^40^. For each test spectrum, we identified candidate molecules whose precursor m/z was within 10 ppm of the query neutral monoisotopic exact mass and selected up to 49 candidates with the highest molecular-fingerprint Tanimoto similarity to the true structure as decoys. Because selection is by precursor m/z rather than exact formula (following FraGNNet), this set can include near-isobaric non-isomers. Together with the true molecule, this yielded at most 50 candidates per query for the NIST, CASMI, and MassSpecGym inverse evaluations. For CASMI, we additionally evaluated a broader PubChem^30^-isomer setting in which candidates were restricted to the dataset-provided true molecular formula, but all PubChem structures with that formula were included rather than limiting the set to 49 decoys; this yielded almost 2.5 million candidate spectra in total.

### MS^2^KOSMOSGenerator: constructing MS^2^KOSMOS, a Map of Known Organic Small- Molecule Space

MS^2^KOSMOS extends chemically grounded fragmentation modeling from individual molecules to the known chemical space captured by PubChem. *MS^2^KOSMOSGenerator* builds this resource by orchestrating *DeepMS^2^Reasoner* across the PubChem-derived small-molecule space and organizing the resulting predicted MS^2^ spectra and fragmentation pathways into a unified, efficiently queryable representation, which will be made publicly available. From predictions generated at four collision energies and two ionization modes, *MS^2^KOSMOSGenerator* derives molecular-formula spectra, in which fragment intensities are aggregated in discrete molecular-formula space rather than continuous, noisy *m/z* space.

Each spectrum is represented as a sparse vector over a mode-specific vocabulary of fragment and neutral-loss formulas defining the Formula Coordinate System. Equivalently, the collection can be represented as a sparse spectra-by-formula matrix whose nonzero entries connect predicted spectra to their fragment formulas with the corresponding intensities. Assigning each peak to a formula coordinate once removes the need to repeat peak alignment for every pairwise spectrum comparison.

MS^2^KOSMOSGenerator stores this sparse representation as an inverted index that maps each fragment formula to the molecules whose predicted spectra contain it. A query therefore visits only formula coordinates populated by the query and scores only molecules sharing at least one such coordinate. Molecule pairs with no shared fragment formula have zero similarity and are never explicitly compared, making similarity search across MS^2^KOSMOS computationally feasible. Exact PubChem filtering, formula-vocabulary construction, spectrum transformation, sparse storage, and index-construction procedures are described in Supplementary AI Methods.

### MS^2^KOSMOSMapper: contextualization of small-molecule MS^2^ spectra in MS^2^KOSMOS

*MS^2^KOSMOSMapper* performs on-demand inference over MS^2^KOSMOS to contextualize experimental MS^2^ spectra through a coarse-to-fine workflow. Because predicted spectra in MS^2^KOSMOS carry exact fragment formulas whereas experimental peaks do not, *MS^2^KOSMOSMapper* first assigns a formula to each experimental peak. Candidate formulas must fall within the specified mass tolerance and be consistent with the precursor formula, with the count of each element no greater than that of the precursor. This places the query in the same Formula Coordinate System as MS^2^KOSMOS while suppressing matches caused only by coincident m/z values. Formula-enumeration, ambiguous-formula assignment, uncertain- precursor handling, and query-cleaning procedures are provided Supplementary AI Methods.

The resulting sparse query vector is searched against the inverted index to retrieve a coarse neighborhood of candidate molecules that share fragment formulas with the query. When several spectra of the same feature are available, for example at different collision energies, their formula representations are combined for coarse retrieval; when only one spectrum is available, it is searched against the nearest precomputed collision-energy condition. The exact aggregation and normalization procedures are described in Supplementary AI Methods.

The coarse candidates are then re-ranked using fresh *DeepMS^2^Reasoner* predictions generated on demand under the query’s exact experimental conditions. Each available query spectrum is compared with the corresponding condition-matched prediction using fragment-formula- informed spectral similarity, and the resulting scores are aggregated across experimental conditions. *MS^2^KOSMOSMapper* returns a ranked candidate list, together with condition-matched predicted spectra and fully annotated fragmentation DAGs, allowing shared peaks to be inspected at the level of putative fragment formulas, structures, and pathways. Because retrieval is based on shared fragment formulas rather than exact whole-molecule identity, the returned neighborhood can remain informative when the query structure itself is absent from PubChem. Complete coarse-retrieval, re-ranking, scoring, and interface procedures are provided in Supplementary AI Methods.

### AIMe annotation of GNPS repository

We used the data made available in Bittremieux et al.^42^, consisting of 1,335 publicly available datasets derived from the GNPS repository. In this previous study, a total of 520,823,130 MS^2^ spectra were processed and 168,193,526 non-singleton spectra were grouped into 8,543,020 non-singleton spectra clusters, of which 454,091 were annotated by GNPS spectral-library searching using a precursor ion tolerance of 2.0 *m/z*, a fragment ion tolerance of 0.5 *m/z*, a minimum cosine similarity of 0.7, and minimum six matched peaks. From this reported dataset, we linked cluster representatives to the GNPS Living Data^42,58^ raw spectra using exact raw-filenames and scan numbers. Consensus spectra were retained when at least one exact match was available, and no contributing spectrum had an explicit label of charge greater than one. A total of 168,193,526 non-singleton spectra were used for AIMe search. We further filtered this dataset for precursors *m/z*<1000, excluding 199,960 spectra. Searches across MS^2^KOSMOS take into account ionization mode, and therefore spectra were categorized as derived from positive ionization (6,321,796 spectra) or negative ionization mode (818,578 spectra), leaving 4,106 spectra with unknown or conflicting ionization labels. The search covered all 7,144,480 analyzed cohort spectra, with zero missing spectra. Spectra with known ionization mode were evaluated against the corresponding predicted ionization mode, whereas spectra with unknown ionization were evaluated against both modes and the higher cosine similarity was retained. To obtain annotations representing likely isomers, the MS^2^KOSMOS search space was restricted to a precursor-mass window of 20 ppm with a minimum absolute window of 0.01 Da. For both the unrestricted MS^2^KOSMOS search and the precursor-restricted likely-isomer analysis, candidates retrieved across the four MS^2^KOSMOS collision energies (20, 30, 50, and 80 NCE) were pooled, and the top 50 candidates per query were retained. Annotations were counted as unique consensus spectra at cosine similarity thresholds of 0.7, 0.8, and 0.9, and previously annotated GNPS spectra were excluded when reporting new AIMe annotations (see Supplementary AI Methods for more details).

### Mice

C57BL/6 (Jax, 000664) and Nr1h4-/- (Jax, 007214) mice were originally purchased from The Jackson Laboratories and bred at Weill Cornell Medicine (WCM). Vnn1-/-mice60 were bred at the Yale School of Medicine (gift of Dr. Phillipe Nasquet (CIML, France) and Dr. Ruslan Medzhitov (Yale University, USA). GF C57BL/6J mice were bred and housed in flexible PVC isolators (Park Bioservices) at WCM. Other mice were maintained under specific pathogen-free condition. All mice used were between 6 and 12 weeks old, age- and sex-matched for each experiment, maintained on a 12-h light–dark cycle, an average ambient temperature of 21 °C and an average humidity of 48%, and provided food and water *ad libitum*. All protocols were approved by the Weill Cornell Medicine Institutional Animal Care and Use Committees (IACUC), and all mice were used in accordance with governmental and institutional guidelines for animal welfare.

### Metabolite extraction from mouse feces

600 µL of methanol were added to 30 mg of feces in 1.7 mL Eppendorf tubes. The tubes were sonicated for 1 min, followed by another 10 min of vigorous stirring. Extracts were pelleted at 5,000 g for 5 min, and supernatants were transferred to 2 mL HPLC vials. Remaining pellets were further extracted with another 10 min of vigorous stirring in 600 µL ethanol. Extracts were pelleted at 5, 000 g for 5 min, and the combined supernatants were then dried in a SpeedVac (ThermoFisher Scientific) vacuum concentrator. Samples were then resuspended in 100 µL of methanol and pelleted at 5,000 g for 5 min, and the clarified extracts were transferred to fresh HPLC vials and stored at −20 °C until analysis.

### MS methods and equipment overview

High-resolution LC-MS was performed on a ThermoFischer Vanquish UHPLC system coupled with a Q-Exactive HF hybrid quadrupole-orbitrap high-resolution mass spectrometer equipped with a HESI ion source. Metabolites were separated using acetonitrile containing 0.1% formic acid (organic phase) and 0.1% formic acid in water (aqueous phase) as solvents on a ThermoFisher Hypersil GOLD C18 column (150 mm × 2.1 mm, particle size 1.8 µm). The gradient started at 1% organic for 3 min after injection and increased linearly to 100% organic over 20 min, then 100% organic for 5 min, and down to 1% organic for 3 min at a flow rate of 0.5 mL/min. Mass spectrometer parameters: spray voltage 3.5 kV, capillary temperature 380 °C, probe heater temperature 400 °C; 60 sheath flow rate, 20 auxiliary flow rate, and one spare gas; S-lens RF level 50, resolution 240,000, AGC target 3 × 106. The instrument was calibrated weekly with positive and negative ion calibration solutions (ThermoFisher). Each sample was analyzed in negative and positive ionization modes using a *m/z* range of 100 to 1000.

### Feature detection and characterization

LC-MS RAW files for all mouse fecal samples were converted to mzXML format (centroid mode) using MSconvert (ProteoWizard version 3.0.18250- 994311be0), followed by analysis using the XCMS analysis feature in Metaboseek version 0.9.7 (metaboseek.com)^72^ based on the centWave XCMS algorithm^73,74^ to extract features. Peak detection values were set as: 4 ppm, 3 to 20 peakwidth, 3 snthresh, 3 and 100 prefilter, FALSE fitgauss, 1 integrate, TRUE firstBaselineCheck, 0 noise, wMean mzCenterFun, -0.005 mzdiff. XCMS feature grouping values were set as: 0.2 minfrac, 2 bw, 0.002 mzwid, 500 max, 1 minsamp, FALSE usegroup. Metaboseek peak filling values set as: 5 ppm 5 rtw, TRUE rtrange. Resulting tables of all detected features were then processed with the Metaboseek data explorer. To remove background derived features, we first applied filters that only retained entries with a retention time window of 1 to 20 min, and then applied maximum intensity (at least one repeat > 10,000), and Peak Quality (>0.98) thresholds, as calculated by Metaboseek^72^. To select differential features, we applied a filter retaining entries with peak area ratios more than 4-fold reduced or 4-fold increased in GF mice compared to SPF mice, as calculated by Metaboseek. We manually curated the resulting list to remove false positive entries, i.e., features that upon manual inspection of raw data were not differential. For verified differential features, we examined elution profiles, isotope patterns, and MS^1^ spectra to find molecular ions and remove adducts, fragments, and isotope peaks. Remaining masses were put on the inclusion list for MS^2^ (ddMS2) characterization. To acquire MS^2^ spectra, we ran a top-10 data dependent MS^2^ method on the Thermo Q-exactive-HF mass spectrometer with MS^1^ resolution 60,000, AGC target 1 × 10^6^, maximum IT (injection time) 50 ms, MS^2^ resolution 45,000, AGC target 5 × 10^5^, maximum IT 80 ms, isolation window 1.0 *m/z*, NCE (normalized collision energy) 20, 35, 50 and 80 for positive and negative ionization mode, dynamic exclusion 3 s.

## Data availability

Source data for Figs. 1-3,5 and Extended Data Figs. 4-8 are provided with this paper. MS data for all mouse metabolome samples analyzed in this study are available at the GNPS website (massive.ucsd.edu) under MassIVE ID number MSV000102710.

## Code availability

During review, AIMe is accessible at https://www.cs.cornell.edu/gomes/udiscoverit/kosmos/ Upon acceptance of the manuscript, AIMe will be accessible as a community resource https://www.cs.cornell.edu/gomes/udiscoverit/kosmos/search.html.

The source code will be made publicly available at https://github.com/gomes-lab/aime and will also be archived in a public long-term repository.

## Supporting information

Supplementary User Guide

## Acknowledgements

We thank all members of the Gomes and Schroeder laboratories for discussion and critical reading of the manuscript, Christian Belardi, Yuanqi Du, and Alexandros Polyzois for assistance with literature searches and helpful discussions and Daniel N. Salter for assistance with chemical syntheses. The figure schematics were created using Adobe Illustrator 2026 30.2.1. All chemical structures were created with ChemDraw 25.5.0. This work was partly supported by an AI2050 Senior Fellowship (to C.P.G.), the use of a Schmidt Sciences high-performance computing cluster, whose GPU resources partly supported the large-scale computations presented in this work (to C.P.G.), and an Eric and Wendy Schmidt AI in Science Postdoctoral Fellowship (to T.J.S.), Schmidt Sciences programs; the National Institute of Food and Agriculture (USDA/NIFA; 2023-67021-39829, to C.P.G); the Air Force Office of Scientific Research (AFOSR; FA9550-23-1-0322, to C.P.G.); the NIH (R35 GM131877 to F.C.S). D.A. and F.C.S. were supported by a grant from the Biocodex Microbiota Foundation. C.N.P. was supported by a Brain and Behavior Research Foundation (NARSAD) Young Investigator Award, NIMH K08MH130773. M.A.F. was supported by an NIH Predoctoral Training Grant (T32GM138826).

## Author contributions

C.P.G. and F.C.S. conceived and supervised the study. U.U.A., D.C. D.F., A.M.F. designed and implemented AIMe under the guidance of C.P.G. and F.C.S., G.J.G. developed a metabolomics framework compatible with AI-based analyses and G.J.G. T.J.S., T.H.W. and M.A.F carried out the metabolomics, and analyzed and curated newly acquired MS^2^ data under the guidance of F.C.S. U.U.A., R.A.B., and G.J.G designed and developed the user interface under the guidance of C.P.G. and F.C.S. H.W. conducted chemical syntheses. C.N.P. and D.A. generated GF and SPF mouse samples and contributed to design of the comparative metabolomics study. U.U.A., G.J.G., T.J.S., F.C.S. and C.P.G. wrote the manuscript with input from all co-authors.

## Competing interests

F.C.S. is a cofounder of Ascribe Bioscience and Holoclara Inc., and a member of the scientific advisory board of Hexagon Bio. D.A. has contributed to scientific advisory boards at Pfizer, Takeda, Nemagene and the Kenneth Rainin Foundation. The other authors declare no competing interests.

## Additional information

Correspondence and requests for materials should be addressed to Carla Gomes or Frank Schroeder.

## Extended Data Figures

**Extended Data Figure 1.**
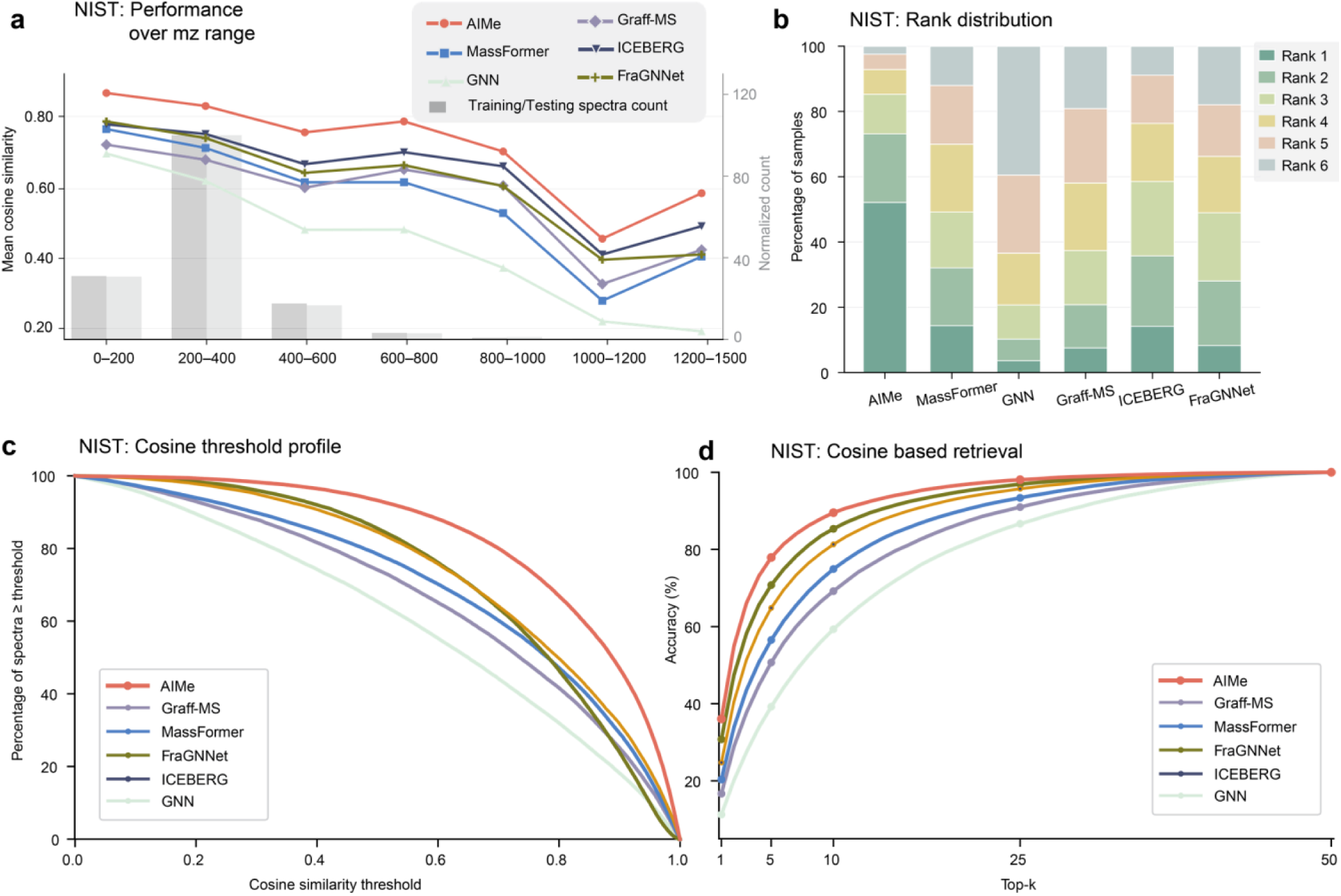
: **Additional metrics for AIMe performance on the NIST test set. a**, Entropy similarity for spectra predicted by all evaluated models compared with experimental spectra. The average cosine was calculated per bin of 200 *m/z*. Grey bars correspond to the normalized number of spectra per bin for both training and testing datasets. **b**, For each experimental spectrum we ranked which model predicted the most similar spectra. AIMe achieved the highest ranked prediction for more than 50% of the NIST test set. Different colors in the stacked bars represent percentages of predicted spectra from a specific model that, when compared with the experimental data by cosine, are most similar, second-most similar etc. among the six evaluated models. **c**, Cosine similarity profile of all tested models, representing the number of spectra above or equal to any given cosine threshold. **d**, NIST test set retrieval using cosine similarity. The top 50 candidates from PubChem, ranked by Tanimoto similarity, were predicted for each model and compared against the corresponding experimental spectrum in the NIST test set, generating top-k retrieval of the correct matching structure.

**Extended Data Figure 2.**
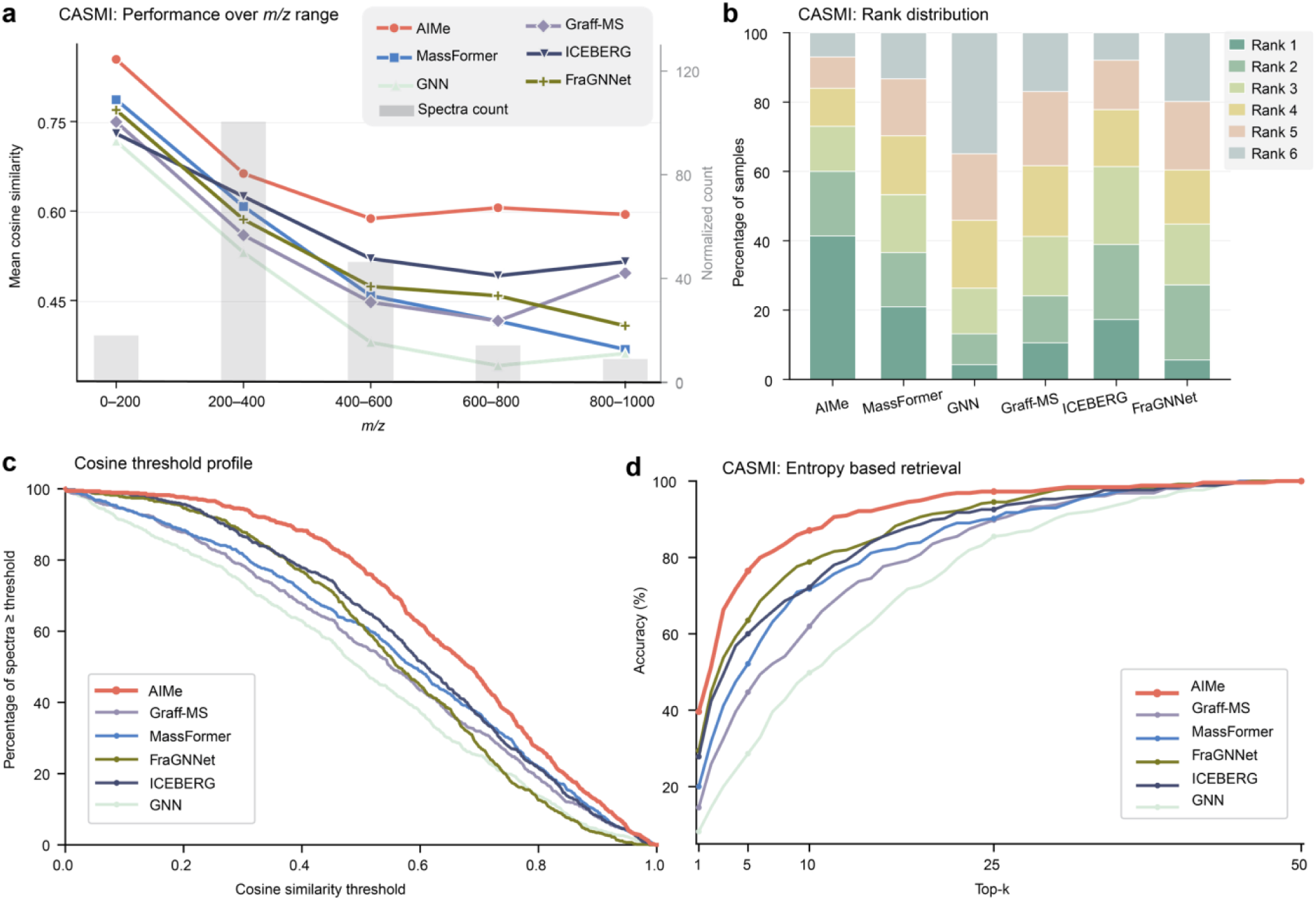
: **Additional metrics for AIMe performance on the CASMI data set. a**, Cosine similarity for spectra predicted by all evaluated models compared with experimental spectra. The average cosine was calculated per bin of 200 *m/z*. Grey bars correspond to the normalized number of spectra per bin in the testing dataset. **b**, For each experimental spectrum we ranked which model predicted the most similar spectra. AIMe achieved the highest ranked prediction for 40% of the spectra in the CASMI test set. Different colors in the stacked bars represent percentages of predicted spectra from a specific model that, when compared with the experimental data by cosine, are most similar, second-most similar etc. among the six evaluated models. **c**, Cosine similarity profile of all tested models, representing the number of spectra above or equal to any given cosine threshold. **d**, CASMI data set retrieval using entropy. The top 50 candidates from PubChem, ranked by Tanimoto similarity, were predicted for each model and compared against the corresponding experimental spectrum in the CASMI data set, generating top-k retrieval of the correct matching structure.

**Extended Data Figure 3.**
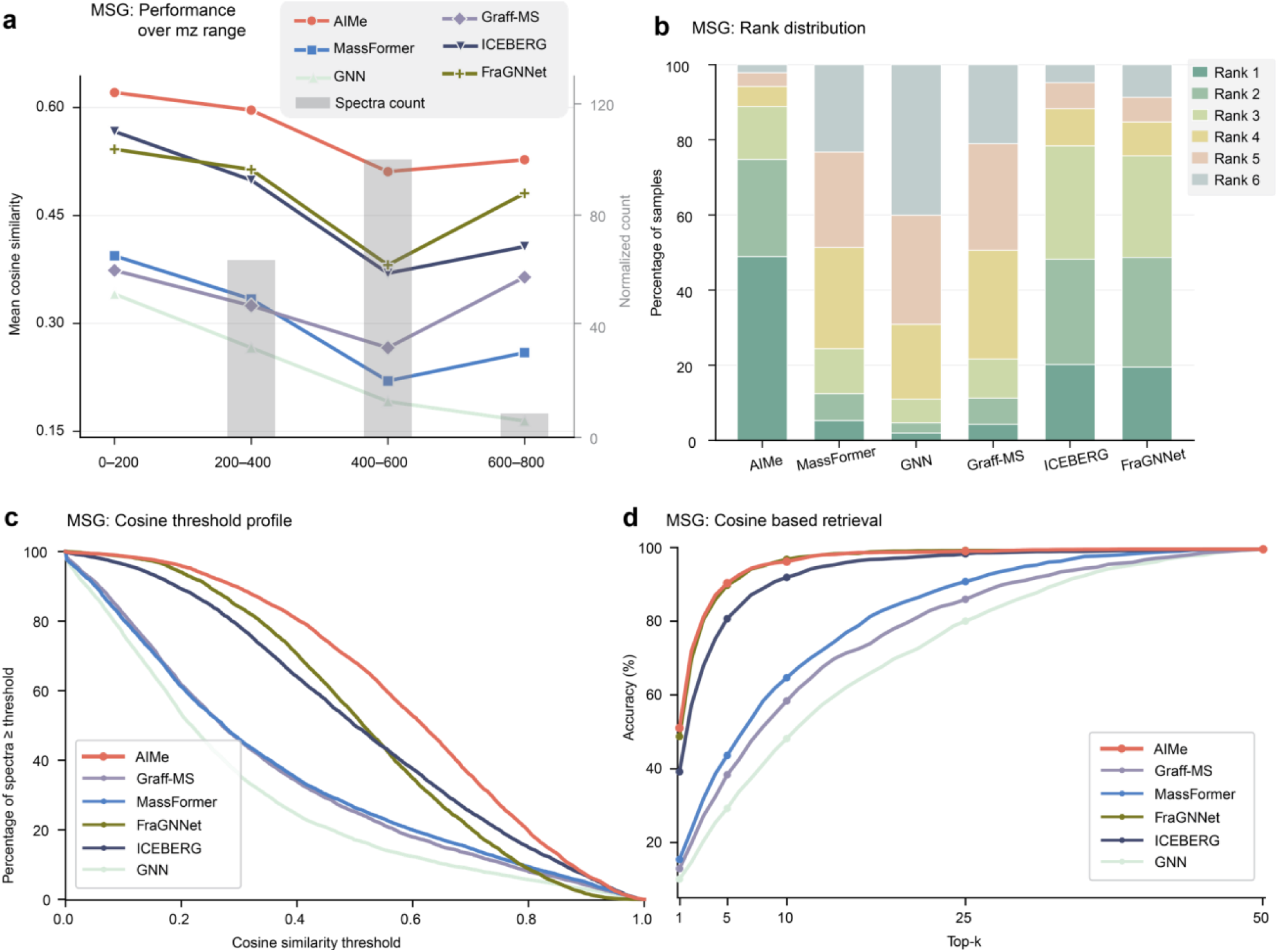
: **Additional metrics for AIMe performance on the MSG data set. a**, Cosine similarity for spectra predicted by all evaluated models compared with experimental spectra. The average cosine was calculated per bin of 200 *m/z*. Grey bars correspond to the normalized number of spectra per bin in the testing dataset. **b**, For each experimental spectrum we ranked which model predicted the most similar spectra. AIMe achieved the highest ranked prediction for near 50% of the MSG test set. Different colors in the stacked bars represent percentages of predicted spectra from a specific model that, when compared with the experimental data by cosine, are most similar, second-most similar etc. among the six evaluated models. **c**, Cosine similarity profile of all tested models, representing the number of spectra above or equal to any given cosine threshold. **d**, MSG data set retrieval using cosine. The top 50 candidates from PubChem, ranked by Tanimoto similarity, were predicted for each model and compared against the corresponding experimental spectrum in the MSG data set, generating top-k retrieval of the correct matching structure.

**Extended Data Figure 4.**
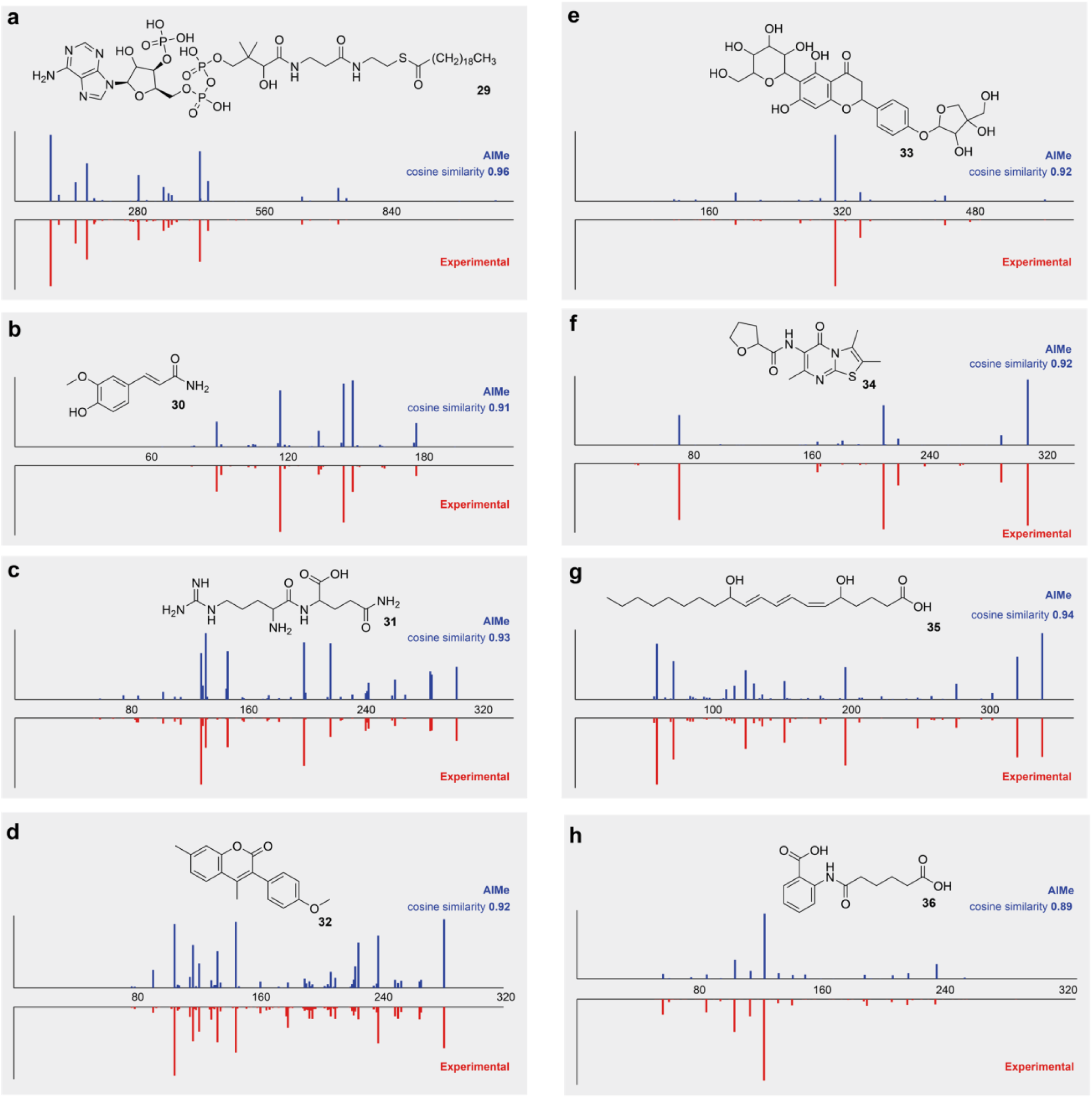
: Additional examples of AIMe-predicted MS^2^ spectra with a collision energy matched to each experimental spectrum. The cosine similarity was calculated for each predicted/experimental pair. a, Arachidoyl-CoA, [M-H]^-^, NCE 35%, cosine similarity 0.96. b, Ferulamide, [M+H]^+^, NCE 65%, cosine similarity 0.91. c, Arg-Gln, [M-H]^-^, NCE 45%, cosine similarity 0.93. d, 4,7- Dimethyl-3(4’-methoxyphenyl)coumarin, [M+H]^+^, NCE 60%, cosine similarity 0.92. e, Naringenin 6-C- glucoside 4′-O-apiofuranoside, [M-H]^-^, NCE 35%, cosine similarity 0.92. f, N-(2,3,7-trimethyl-5-oxo-5H- thiazolo[3,2-a]pyrimidin-6-yl)tetrahydrofuran-2-carboxamide, [M+H]^+^, NCE 30%, cosine similarity 0.92. g, Leukotriene B3, [M-H]^-^, NCE 40%, cosine similarity 0.94. h, 2-(5-carboxypentanoylamino)benzoic acid, [M+H]^+^, NCE 45%, cosine similarity 0.89.

**Extended Data Figure 5.**
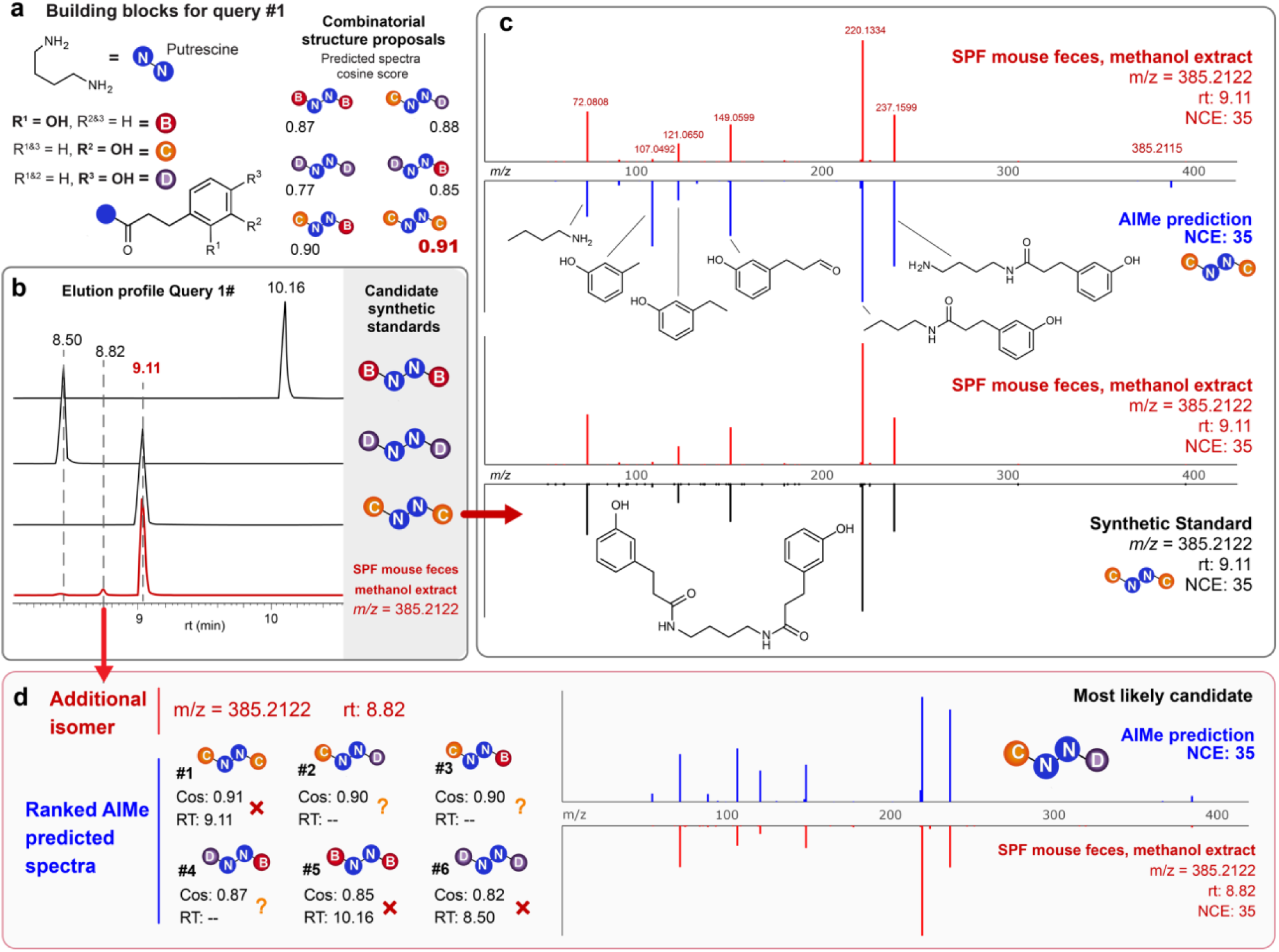
: **Structural validation of query#1 proposed structure. a,** Building blocks derived from MS^2^ KOSMOS retrieved candidates allowed to propose a combinatorial space of possible structures. Each structure was predicted and compared against the experimental spectra, with the R^2^ (C) positional isomer on both terminal ends of the putrescine scaffold producing the highest cosine similarity score. **b**, The elution profile for the synthetic standard of the highest-ranking candidate matched exactly to the mouse metabolome extract. **c**, MS^2^ Mirror plots of the synthetic standard, predicted and SPF mouse metabolome extract, further illustrating AIMe structural annotations for each matching fragment. **d**, An additional isomer at 8.82 min lies between the R^2^ (C) and R^3^ (D) isomers of the synthetic standards, suggesting a different conformation on the terminal building blocks. Comparing this experimental MS^2^ spectrum against the predicted candidates and the elution profiles suggests C and D arrangement as the most likely structure.

**Extended Data Figure 6.**
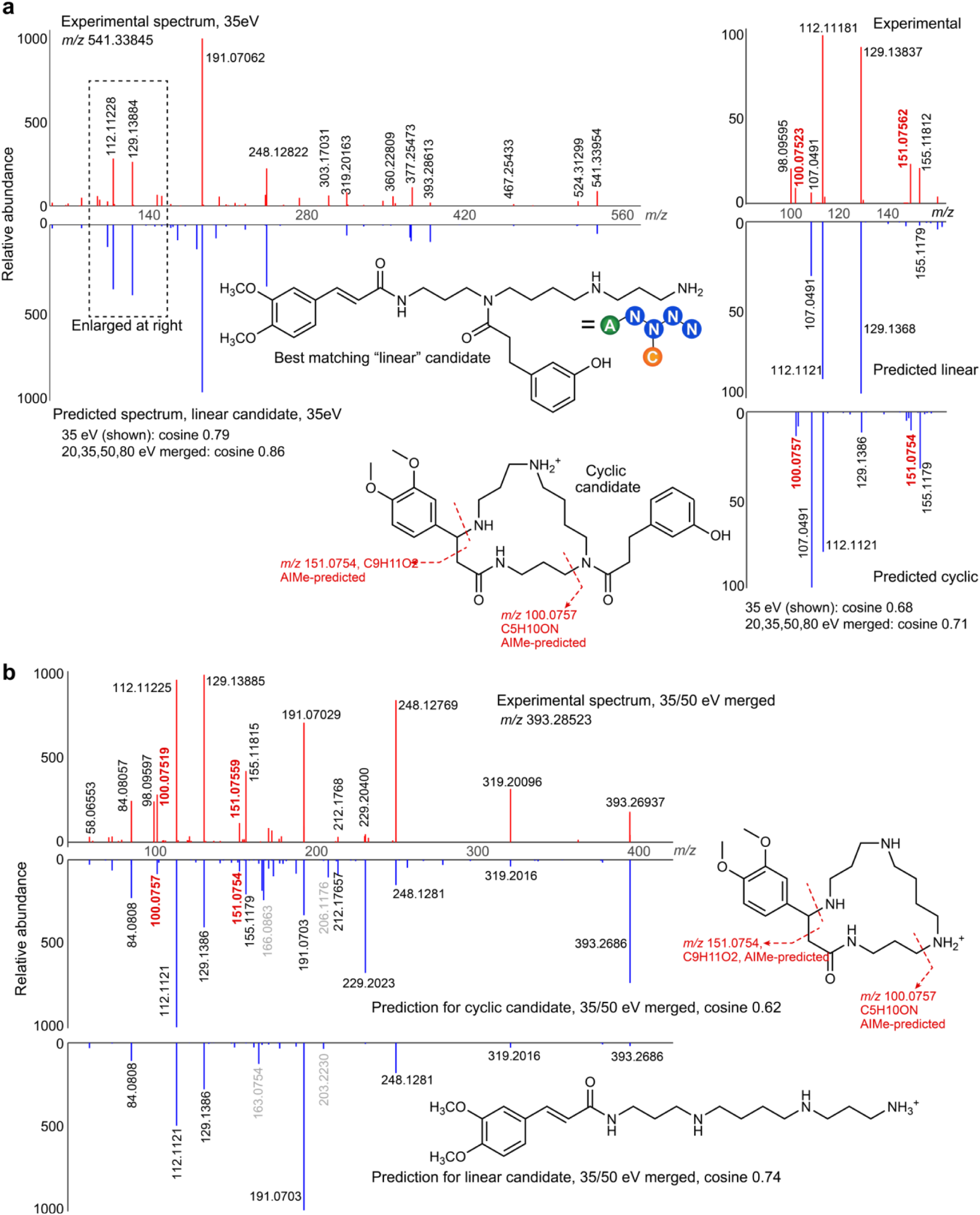
: Predictions for candidates identified for query#2. **a,** Comparison of experimental MS^2^ spectrum of *m/z* 541.33854 (C_30_H_45_N_4_O_5_^+^) and predicted spectra for the best-matching candidate shown in Figure 5 (“linear” isomer, merged cosine 0.86). Despite otherwise excellent fit, fragments at *m/z* 100.0757 and 151.0754 (highlighted in red) are absent in the prediction for this linear isomer (and any other predicted linear candidate). This mismatch led to expanding our search to consider additional variants for moiety A (i.e., isomers of 3,4-dimethoxycinnamoyl) as well as other connection variants. Within this search space, only the shown cyclic candidate was predicted to generate *m/z* 100.0757 and 151.0754. A parallel search for other microbiota-dependent compounds in our data set whose MS^2^ spectra showed prominent fragments at *m/z* 100.0757 and 151.0754 revealed several putatively related metabolites (see Supplementary Fig. 3), including a metabolite represented by *m/z* 393.28523 (C_21_H_37_N_4_O_3_^+^). **b,** Comparison of experimental MS^2^ spectrum of *m/z* 393.28523 (C_21_H_37_N_4_O_3_^+^) and predicted spectra for cyclic and linear candidates (measured at and predicted for 35 eV). As in **a**, key fragments *m/z* 100.0757 and 151.0754 appear only in the prediction for the cyclic candidate.

**Extended Data Figure 7.**
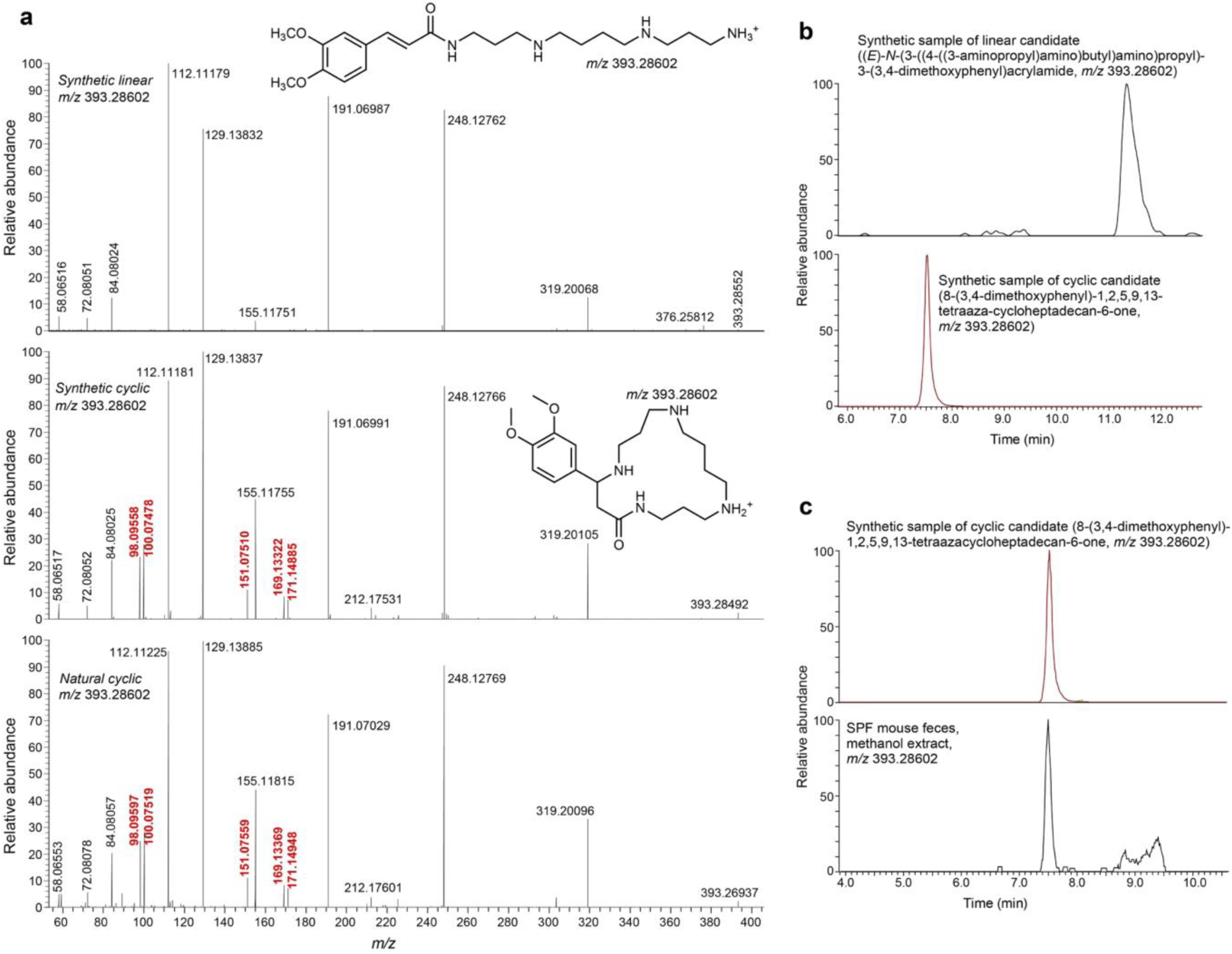
: Comparison of the MS^2^ spectra and retention times of synthetic samples obtained for the linear and cyclic candidates for *m/z* 393.28523 (C_21_H_37_N_4_O_3_^+^). **a,** Similar to the corresponding predicted spectrum (**Extended Data** Fig. 6), the MS^2^ spectrum of the linear isomer (top) lacks key fragments (highlighted in red) present in the MS^2^ spectra of the cyclic isomer (middle), which matches the MS^2^ spectrum (bottom) of the natural compound. **b,** HPLC (using a HILIC column) retention times of synthetic linear (top) and cyclic isomers differ greatly. **c,** Retention times of the natural compound (top) and the synthetic sample of the cyclic isomer (bottom) match.

**Extended Data Figure 8.**
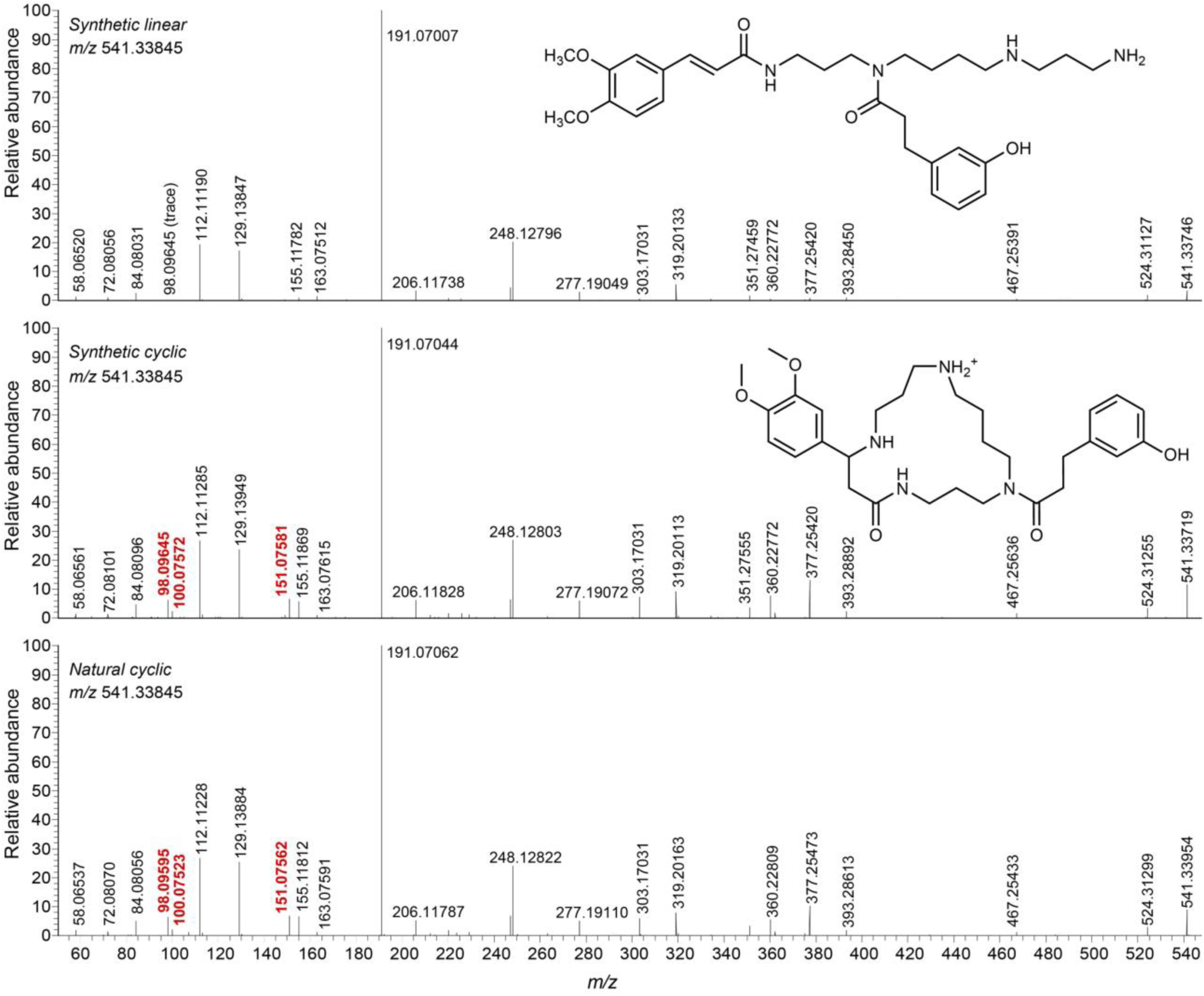
: Comparison of the MS^2^ spectra of synthetic samples obtained for the linear and cyclic candidates for *m/z* 541.33854 (C_30_H_45_N_4_O_5_^+^). Similar to the corresponding predicted spectrum (**Extended Data** Fig. 6), the experimental MS^2^ spectrum of the linear isomer (top) lacks key fragments (highlighted in red) present in the experimental MS^2^ spectra of a cyclic isomer (middle), which matches the MS^2^ spectrum (bottom) of the natural compound.

**Extended Data Figure 9.**
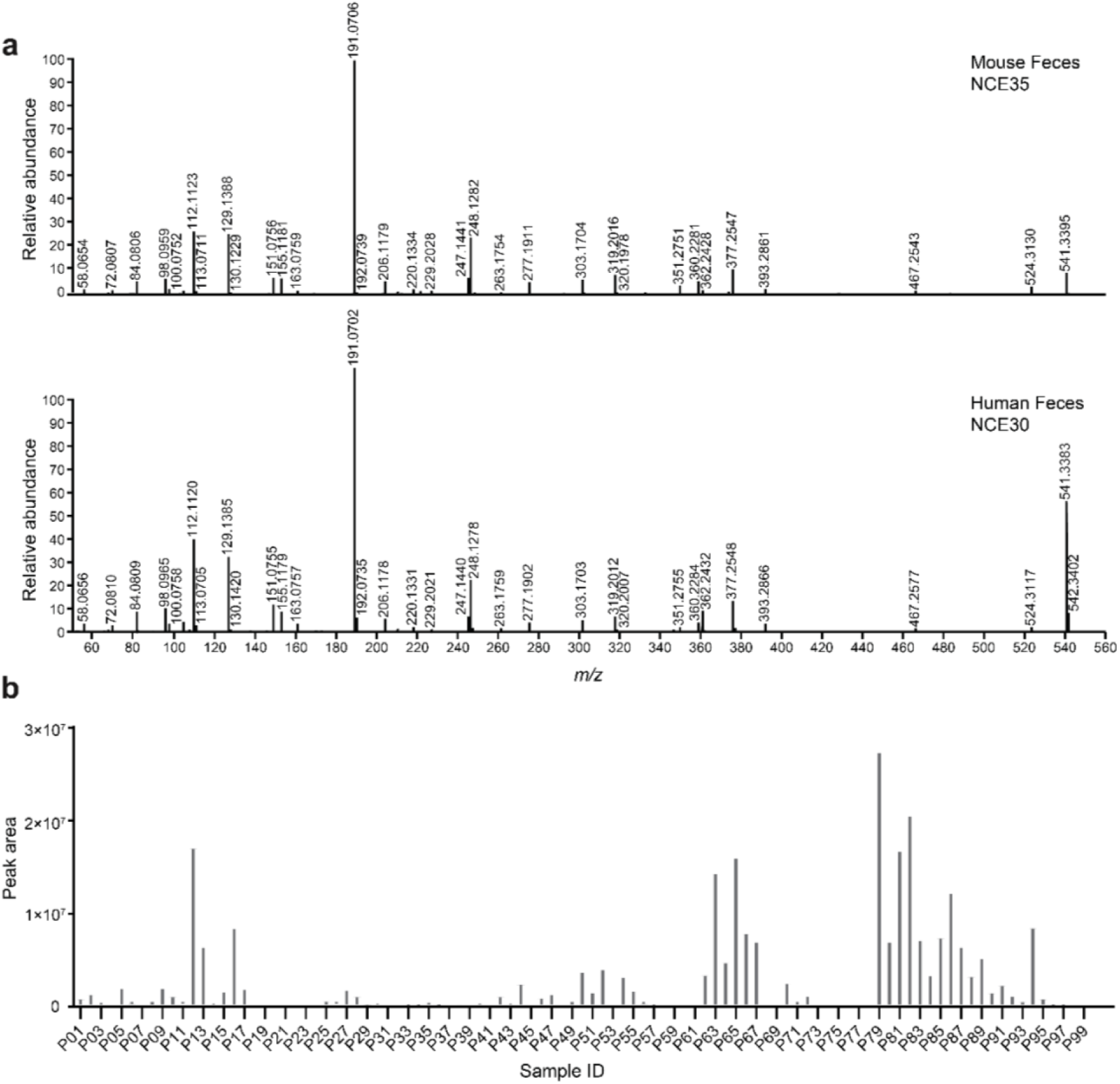
Detection of query#2 (*N*,*N*’-(butane-1,4-diyl)bis(3-(3-hydroxyphenyl) propanamide)) in human fecal metabolomics data (MSV000082629). **a**, Mirror plot comparing the MS^2^ spectrum of query#2 detected in mouse feces with a spectrum acquired from human feces. The mouse and human spectra were acquired at normalized collision energies (NCE) of 35 and 30, respectively, and had a cosine similarity of 0.93. **b**, Blank-subtracted peak areas of the feature corresponding to query#2 across 99 human fecal samples from the publicly available MassIVE dataset MSV000082629. Each bar represents an individual human fecal sample. The feature corresponding to query#2 was detected in 57 out of 99 human fecal samples using a peak area threshold of 3E^5^.

