## Supplementary for "Charting the small-molecule universe from mass spectra with neuro-symbolic AI"

### Table of Contents

Supplementary Tables:

Supplementary Table 1. NIST test set forward prediction metrics 3

Supplementary Table 2: CASMI forward prediction metrics 3

Supplementary Table 3: MassSpecGym forward prediction metrics 3

Supplementary Table 4: NIST retrieval metrics 4

Supplementary Table 5: CASMI retrieval metrics 5

Supplementary Table 6: MassSpecGym retrieval metrics 6

Supplementary Table 7: MS features of the 111 most abundant unidentified microbiota-dependent metabolites 6

Supplementary Table 8: Additional information for the numbered structures in the figures 6

Supplementary Figures:

Supplementary Figure 1: AIMe predicted spectra for structures in Fig. 5b 7

Supplementary Figure 2: AIMe predicted spectra for structures in Fig. 5h 8

Supplementary Figure 3: $MS^2$ spectra of other putatively related microbiota-dependent metabolites 9

Supplementary Synthesis Methods 10

Supplementary References 14

### Supplementary Tables

**Supplementary Table 1.** AIMe forward prediction performance for the NIST test set. Reported similarity of the predicted to the experimental spectra, calculated for both positive and negative ionization modes and averaged using cosine and entropy.

| | Cosine similarity | | | Entropy similarity | | |
|---|---|---|---|---|---|---|
| **Model** | **Average** | **ESI+** | **ESI-** | **Average** | **ESI+** | **ESI -** |
| ***AIMe*** | 0.83 | 0.84 | 0.81 | 0.73 | 0.74 | 0.72 |
| **ICEBERG** | 0.74 | 0.75 | 0.73 | 0.67 | 0.67 | 0.66 |
| **FraGNNet** | 0.73 | 0.74 | 0.72 | 0.55 | 0.56 | 0.53 |
| **MassFormer** | 0.70 | 0.71 | 0.69 | 0.64 | 0.64 | 0.62 |
| **Graff-MS** | 0.67 | 0.68 | 0.66 | 0.60 | 0.60 | 0.58 |
| **GNN** | 0.61 | 0.62 | 0.59 | 0.53 | 0.54 | 0.52 |

**Supplementary Table 2.** AIMe forward prediction performance for CASMI. Reported similarity of the predicted to the experimental spectra, calculated for both positive and negative ionization modes and averaged using cosine and entropy.

| | Cosine similarity | | | Entropy similarity | | |
|---|---|---|---|---|---|---|
| **Model** | **Average** | **ESI+** | **ESI-** | **Average** | **ESI+** | **ESI -** |
| ***AIMe*** | 0.66 | 0.67 | 0.637 | 0.56 | 0.58 | 0.52 |
| **ICEBERG** | 0.59 | 0.63 | 0.546 | 0.56 | 0.59 | 0.51 |
| **MassFormer** | 0.56 | 0.60 | 0.513 | 0.54 | 0.59 | 0.48 |
| **FraGNNet** | 0.56 | 0.566 | 0.546 | 0.42 | 0.44 | 0.39 |
| **Graff-MS** | 0.54 | 0.57 | 0.481 | 0.52 | 0.56 | 0.46 |
| **GNN** | 0.49 | 0.53 | 0.435 | 0.48 | 0.53 | 0.42 |

**Supplementary Table 3.** AIMe forward prediction performance on the MassSpecGym dataset. Reported cosine and entropy similarity metrics of the predicted to the experimental spectra, calculated for positive ionization mode and averaged. The MassSpecGym data set does not contain spectra acquired in negative ionization mode.

| | Cosine similarity | | | Entropy similarity | | |
|---|---|---|---|---|---|---|
| **Model** | **Average** | **ESI+** | **ESI-** | **Average** | **ESI+** | **ESI -** |
| ***AIMe*** | 0.594 | 0.594 | -/- | 0.506 | 0.506 | -/- |
| **ICEBERG** | 0.515 | 0.515 | -/- | 0.462 | 0.462 | -/- |
| **FraGNNet** | 0.514 | 0.514 | -/- | 0.311 | 0.311 | -/- |
| **MassFormer** | 0.383 | 0.383 | -/- | 0.331 | 0.331 | -/- |
| **Graff-MS** | 0.363 | 0.363 | -/- | 0.309 | 0.309 | -/- |
| **GNN** | 0.305 | 0.305 | -/- | 0.265 | 0.265 | -/- |

**Supplementary Table 4.** AIMe retrieval performance on the NIST test set using cosine and entropy. For every compound in the test set, the 50 most similar isomers in PubChem, measured by Tanimoto similarity to the query structure, were predicted. Results represent the percentage of the test set for which the correct predicted spectra were among the top 1 to 50. Spectra retrieval was performed using two methods: single-spectrum, where each individual spectrum was paired and their similarity was compared, and multi-spectra, where spectra for different collision energies of the same compound were compared and the results were aggregated to provide a final Top-*k* rank.

| NIST test set retrieval - top-k (%) | | | | | | | | | |
|---|---|---|---|---|---|---|---|---|---|
| **Metric** | **Model** | **Single spectrum** | | | | **Multi-spectra** | | | |
| | | ***k-1*** | ***k-5*** | ***k-10*** | ***k-50*** | ***k-1*** | ***k-5*** | ***k-10*** | ***k-50*** |
| *Cosine* | ***AIMe*** | 34.63 | 74.83 | 87.43 | 100 | 42.140 | 81.40 | 91.05 | 100 |
| *Cosine* | **FraGNNet** | 27.29 | 67.12 | 82.10 | 100 | 36.13 | 76.44 | 88.18 | 100 |
| *Cosine* | **ICEBERG** | 20.66 | 57.51 | 75.27 | 100 | 27.27 | 66.45 | 81.28 | 100 |
| *Cosine* | **MassFormer** | 17.05 | 50.02 | 68.43 | 100 | 22.64 | 58.78 | 75.7 | 100 |
| *Cosine* | **Graff-MS** | 14.79 | 45.69 | 63.56 | 100 | 19.35 | 52.76 | 70.33 | 100 |
| *Cosine* | **GNN** | 9.35 | 35.44 | 53.74 | 100 | 11.94 | 41.40 | 60.88 | 100 |
| *Entropy* | ***AIMe*** | 36.07 | 77.91 | 89.54 | 100 | 42.81 | 82.36 | 91.89 | 100 |
| *Entropy* | **FraGNNet** | 30.78 | 70.76 | 85.34 | 100 | 36.70 | 76.32 | 89.00 | 100 |
| *Entropy* | **ICEBERG** | 24.74 | 64.83 | 81.31 | 100 | 30.35 | 70.89 | 85.40 | 100 |
| *Entropy* | **MassFormer** | 20.35 | 56.52 | 74.92 | 100 | 24.58 | 63.39 | 79.99 | 100 |
| *Entropy* | **Graff-MS** | 16.68 | 50.68 | 69.19 | 100 | 20.65 | 57.49 | 75.23 | 100 |
| *Entropy* | **GNN** | 11.27 | 39.21 | 59.33 | 100 | 14.97 | 45.49 | 64.95 | 100 |

**Supplementary Table 5.** AIMe retrieval performance on the CASMI data set using **a)** single spectrum retrieval and **b)** multi-spectra retrieval. For each compound in the test set, retrieval is reported for the 50 most similar structures in PubChem to the query structure, and is also calculated across all isomers found in PubChem. Results represent the percentage of the CASMI set for which the correct predicted spectra were among the top 1 to 50 using both entropy and cosine metrics.

| **a)** | | **Single spectrum retrieval (%)** | | | | | | | |
|---|---|---|---|---|---|---|---|---|---|
| | | **Top-1** | | **Top-5** | | **Top-10** | | **Top-50** | |
| **Metric** | **Model** | **All CASMI isomers** | **50 PubChem isomers** | **All CASMI isomers** | **50 PubChem isomers** | **All CASMI isomers** | **50 PubChem isomers** | **All CASMI isomers** | **50 PubChem isomers** |
| *Cosine* | ***AIMe*** | 29.32 | 35.46 | 50.14 | 68.39 | 58.49 | 81.94 | 73.15 | 100 |
| *Cosine* | **ICEBERG** | 13.01 | 18.33 | 30.00 | 52.72 | 39.73 | 69.32 | 60.82 | 100 |
| *Cosine* | **MassFormer** | 10.55 | 13.15 | 24.11 | 44.75 | 31.78 | 64.28 | 55.62 | 100 |
| *Cosine* | **FraGNNet** | 22.88 | 30.41 | 43.29 | 61.75 | 52.74 | 76.10 | 68.49 | 100 |
| *Cosine* | **Graff-MS** | 8.90 | 11.16 | 20.96 | 38.51 | 29.18 | 56.71 | 51.51 | 100 |
| *Cosine* | **GNN** | 8.08 | 7.57 | 17.53 | 28.29 | 23.15 | 46.61 | 43.42 | 100 |
| *Entropy* | ***AIMe*** | 32.60 | 36.52 | 52.60 | 72.24 | 60.82 | 84.86 | 77.81 | 100 |
| *Entropy* | **FraGNNet** | 21.51 | 27.09 | 39.73 | 60.56 | 49.18 | 76.36 | 66.16 | 100 |
| *Entropy* | **ICEBERG** | 15.89 | 21.91 | 35.62 | 55.11 | 45.75 | 70.65 | 65.34 | 100 |
| *Entropy* | **MassFormer** | 14.38 | 18.33 | 30.41 | 50.60 | 39.73 | 68.13 | 62.05 | 100 |
| *Entropy* | **Graff-MS** | 10.68 | 13.81 | 24.93 | 41.04 | 34.66 | 61.89 | 56.16 | 100 |
| *Entropy* | **GNN** | 6.58 | 7.04 | 17.12 | 29.08 | 25.34 | 49.54 | 46.58 | 100 |

| **b)** | | **Multi-spectra retrieval (%)** | | | | | | | |
|---|---|---|---|---|---|---|---|---|---|
| | | **Top-1** | | **Top-5** | | **Top-10** | | **Top-50** | |
| **Metric** | **Model** | **All CASMI isomers** | **50 PubChem isomers** | **All CASMI isomers** | **50 PubChem isomers** | **All CASMI isomers** | **50 PubChem isomers** | **All CASMI isomers** | **50 PubChem isomers** |
| *Cosine* | ***AIMe*** | 33.20 | 38.82 | 54.66 | 70.20 | 61.13 | 84.31 | 74.49 | 100 |
| *Cosine* | **ICEBERG** | 13.77 | 20.00 | 31.98 | 56.08 | 42.11 | 71.37 | 65.18 | 100 |
| *Cosine* | **MassFormer** | 12.55 | 12.94 | 27.13 | 49.02 | 35.22 | 68.24 | 58.70 | 100 |
| *Cosine* | **FraGNNet** | 27.13 | 36.08 | 46.15 | 66.27 | 57.09 | 78.43 | 73.28 | 100 |
| *Cosine* | **Graff-MS** | 10.93 | 13.73 | 23.48 | 40.39 | 32.39 | 58.04 | 54.66 | 100 |
| *Cosine* | **GNN** | 7.69 | 8.24 | 18.62 | 31.76 | 23.48 | 47.45 | 46.96 | 100 |
| *Entropy* | ***AIMe*** | 35.63 | 39.61 | 57.09 | 76.47 | 64.37 | 87.06 | 79.76 | 100 |
| *Entropy* | **FraGNNet** | 21.46 | 29.41 | 43.32 | 63.53 | 53.04 | 78.82 | 70.85 | 100 |
| *Entropy* | **ICEBERG** | 19.43 | 27.84 | 39.27 | 60.00 | 51.82 | 72.16 | 69.64 | 100 |
| *Entropy* | **MassFormer** | 15.79 | 20.00 | 33.20 | 52.16 | 42.11 | 71.76 | 63.56 | 100 |
| *Entropy* | **Graff-MS** | 12.55 | 14.51 | 27.94 | 44.71 | 35.63 | 61.96 | 58.70 | 100 |
| *Entropy* | **GNN** | 9.31 | 8.24 | 18.62 | 28.63 | 27.53 | 49.80 | 48.58 | 100 |

**Supplementary Table 6.** AIMe retrieval performance on the MassSpecGym data set using **a)** single spectrum retrieval and **b)** multi-spectra retrieval. For each compound in the test set, retrieval is calculated across all isomers in PubChem and across the 250 structures with the smallest Maximum Common Edge Subgraph distance to the query structure. Results represent the percentage of the CASMI set for which the correct predicted spectra were among the top 1 to 50 using both entropy and cosine metrics for comparing spectra.

| **a)** | | **Single spectrum match (%)** | | | | | | | |
|---|---|---|---|---|---|---|---|---|---|
| | | **Top-1** | | **Top-5** | | **Top-10** | | **Top-50** | |
| **Metric** | **Model** | **All MSG isomers** | **MCES filter** | **All MSG isomers** | **MCES filter** | **All MSG isomers** | **MCES filter** | **All MSG isomers** | **MCES filter** |
| *Cosine* | ***AIMe*** | 36.84 | 40.58 | 79.32 | 83.62 | 90.73 | 93.64 | 100 | 100 |
| *Cosine* | **FraGNNet** | 36.57 | 38.70 | 78.75 | 81.54 | 89.89 | 92.04 | 100 | 100 |
| *Cosine* | **ICEBERG** | 25.30 | 28.74 | 65.27 | 69.33 | 82.19 | 85.24 | 100 | 100 |
| *Cosine* | **MassFormer** | 8.67 | 8.45 | 35.47 | 33.29 | 57.25 | 55.68 | 100 | 100 |
| *Cosine* | **Graff-MS** | 7.59 | 7.48 | 30.62 | 30.43 | 50.41 | 50.06 | 100 | 100 |
| *Cosine* | **GNN** | 4.96 | 4.68 | 22.22 | 21.95 | 40.48 | 39.61 | 100 | 100 |
| *Entropy* | ***AIMe*** | 43.02 | 47.44 | 83.38 | 87.56 | 92.84 | 95.43 | 100 | 100 |
| *Entropy* | **FraGNNet** | 39.51 | 42.98 | 80.28 | 83.57 | 91.32 | 93.47 | 100 | 100 |
| *Entropy* | **ICEBERG** | 32.26 | 36.10 | 73.14 | 77.60 | 87.15 | 90.68 | 100 | 100 |
| *Entropy* | **MassFormer** | 10.58 | 10.68 | 39.54 | 37.96 | 60.52 | 58.81 | 100 | 100 |
| *Entropy* | **Graff-MS** | 9.02 | 9.24 | 34.07 | 33.29 | 53.91 | 52.82 | 100 | 100 |
| *Entropy* | **GNN** | 6.02 | 6.30 | 25.67 | 25.38 | 43.23 | 42.77 | 100 | 100 |

| **b)** | | **Multi-spectra match (%)** | | | | | | | |
|---|---|---|---|---|---|---|---|---|---|
| | | **Top-1** | | **Top-5** | | **Top-10** | | **Top-50** | |
| **Metric** | **Model** | **All MSG isomers** | **MCES filter** | **All MSG isomers** | **MCES filter** | **All MSG isomers** | **MCES filter** | **All MSG isomers** | **MCES filter** |
| *Cosine* | ***AIMe*** | 47.87 | 49.08 | 89.29 | 90.35 | 96.29 | 96.48 | 100 | 100 |
| *Cosine* | **FraGNNet** | 46.47 | 46.64 | 89.24 | 89.75 | 96.85 | 97.13 | 100 | 100 |
| *Cosine* | **ICEBERG** | 34.93 | 36.61 | 78.94 | 80.21 | 91.51 | 91.97 | 100 | 100 |
| *Cosine* | **MassFormer** | 12.52 | 11.61 | 43.41 | 41.21 | 64.94 | 63.39 | 100 | 100 |
| *Cosine* | **Graff-MS** | 10.16 | 9.00 | 37.29 | 35.74 | 58.53 | 56.78 | 100 | 100 |
| *Cosine* | **GNN** | 6.45 | 6.02 | 26.72 | 26.14 | 46.99 | 46.04 | 100 | 100 |
| *Entropy* | ***AIMe*** | 54.73 | 56.51 | 92.39 | 93.06 | 97.22 | 97.29 | 100 | 100 |
| *Entropy* | **FraGNNet** | 50.42 | 51.25 | 90.07 | 90.56 | 97.26 | 97.51 | 100 | 100 |
| *Entropy* | **ICEBERG** | 43.27 | 44.69 | 85.34 | 86.71 | 94.85 | 95.39 | 100 | 100 |
| *Entropy* | **MassFormer** | 14.61 | 13.94 | 48.47 | 47.02 | 68.88 | 67.25 | 100 | 100 |
| *Entropy* | **Graff-MS** | 12.29 | 11.33 | 42.35 | 40.84 | 62.48 | 60.57 | 100 | 100 |
| *Entropy* | **GNN** | 7.93 | 7.65 | 31.45 | 31.02 | 50.74 | 49.84 | 100 | 100 |

**Supplementary Table 7:** MS data for the 111 most abundant unidentified microbiota-dependent metabolites. This table is provided as a separate file.

**Supplementary Table 8:** Additional information for the numbered structures in the main document figures detailing SMILES strings, monoisotopic mass, molecular formula, ionization mode, collision energy and experimental spectra source. This table is provided as a separate file.

### Supplementary Figures

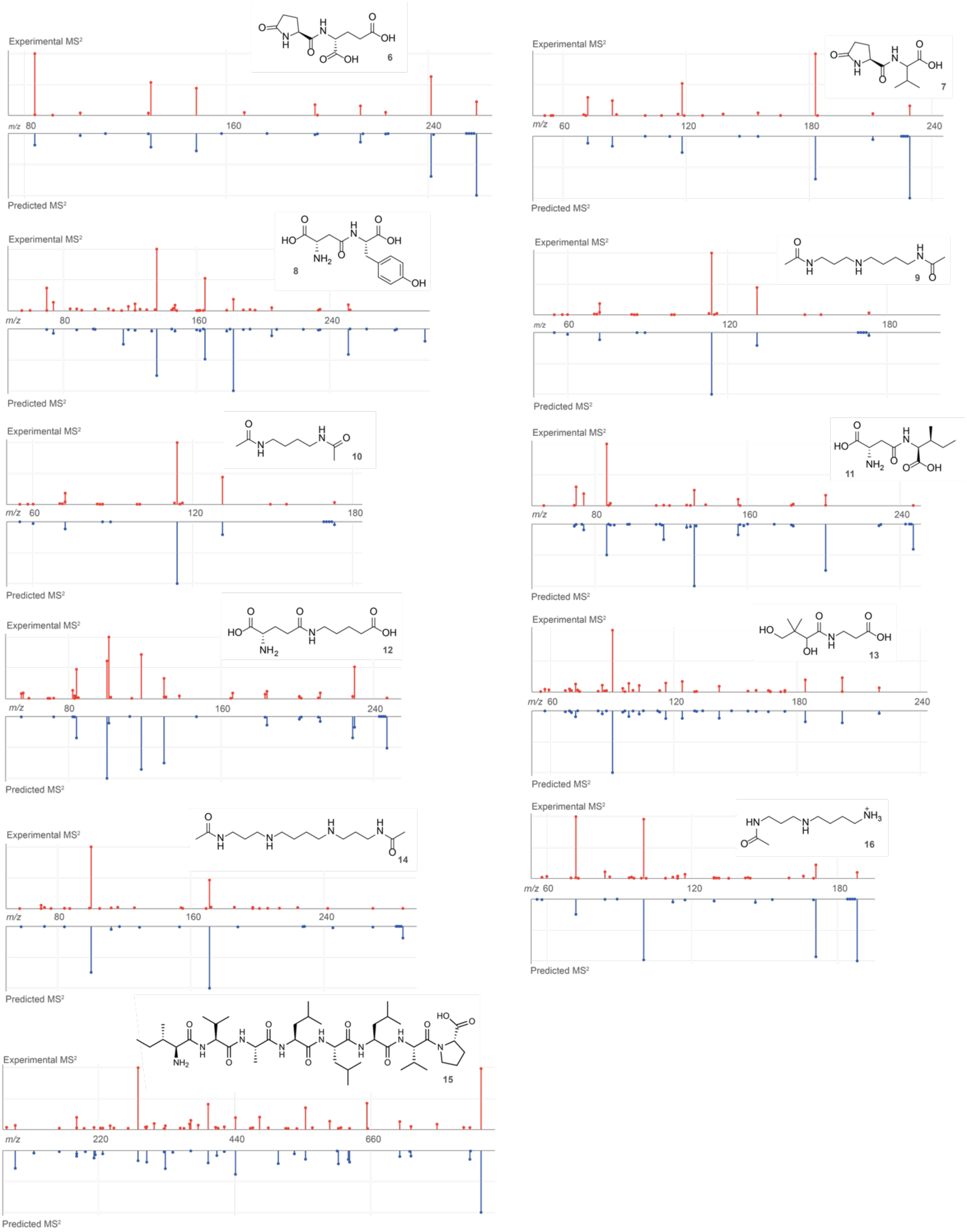


**Supplementary Figure 1.** AIMe-predicted spectra for metabolites represented in Fig. 5b that had close matches with MS²KOSMOS spectra, suggesting that they represent known compounds or closely related isomers. Structure numbers shown in bold. Further details for each structure can be found in Supplementary Table 8.

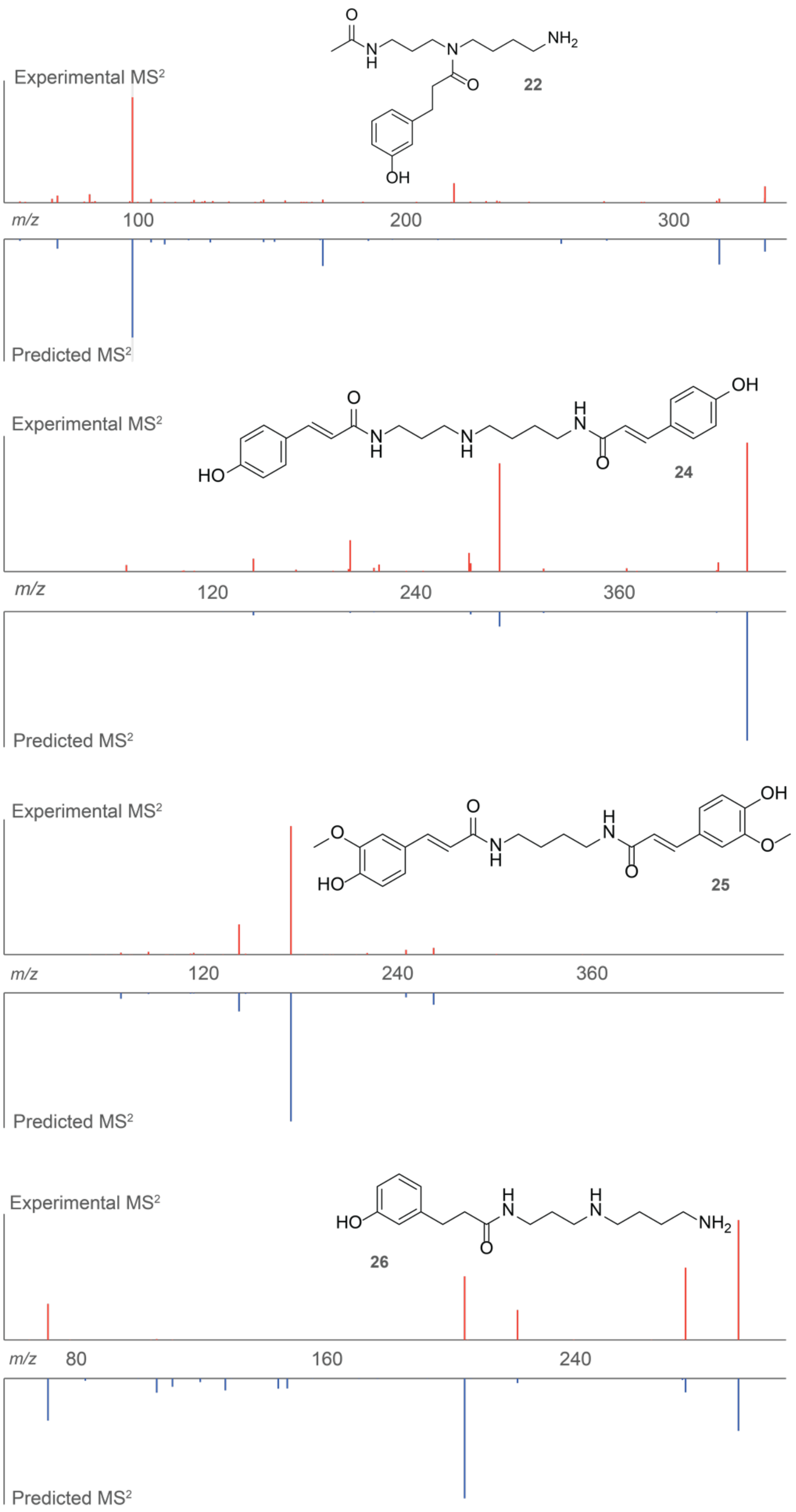


**Supplementary Figure 2**. AIMe predicted $MS^2$ spectra for additional previously undescribed polyamine derivatives listed in Fig. 5h, compared against the experimental spectra. Structure numbers are represented in bold. Further details of each structure can be found in Supplementary Table 8.

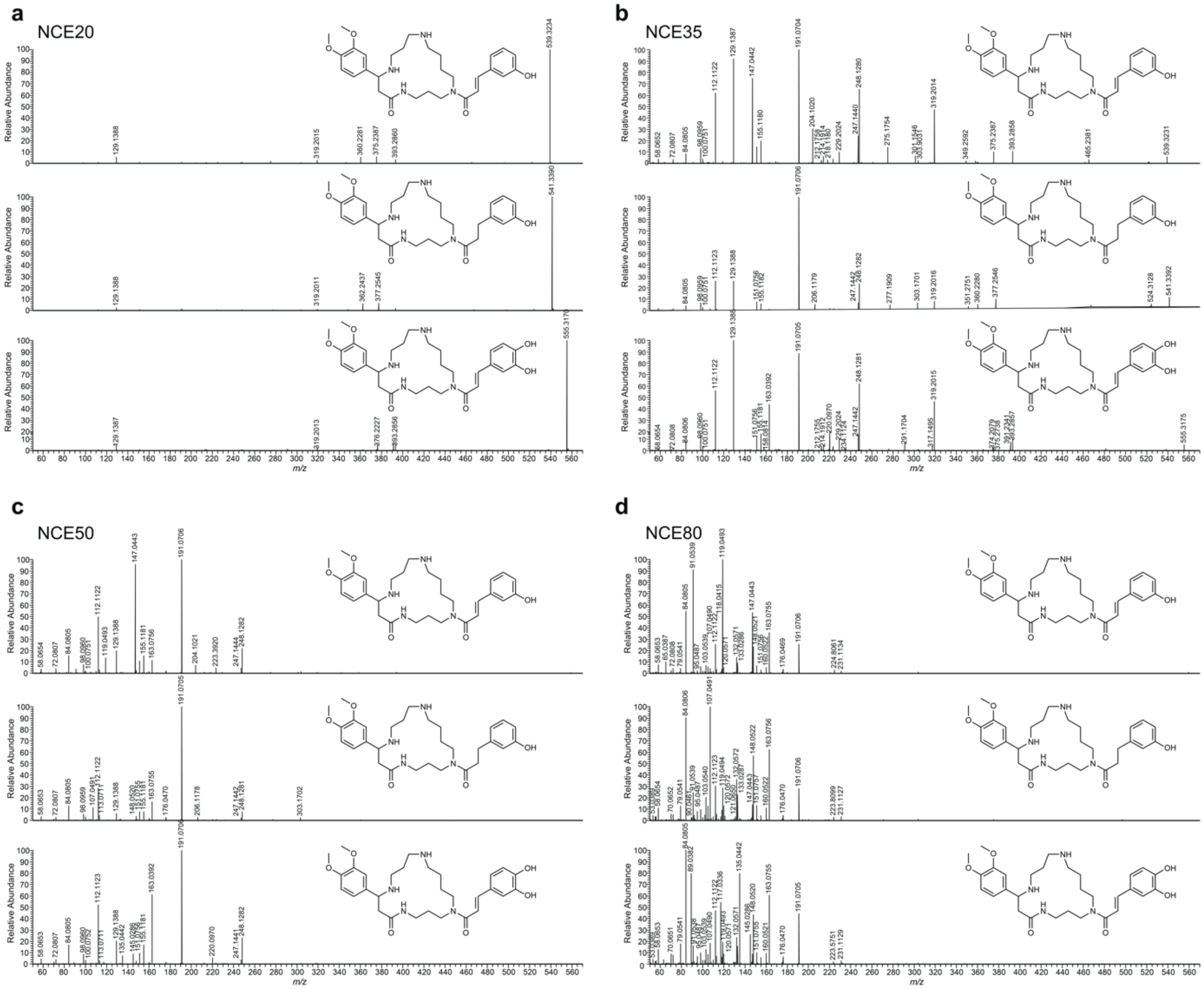


**Supplementary Figure 3**. $MS^2$ spectra and putative structures of metabolites related to the newly identified cyclic polyamide metabolite shown in Fig.5 c, Query#2. **a-d**, spectra collected at NCE: 20, 35, 50 and 80 in positive mode ionization.

### Supplementary Synthesis Methods

**General procedures for chemical syntheses.** Unless noted otherwise, all chemicals and reagents were purchased from Sigma-Aldrich. Solutions and solvents sensitive to moisture and oxygen were transferred via standard syringe and cannula techniques. Acetic acid, acetonitrile, dichloromethane, and methanol used for chromatography and as a reagent or solvent were purchased from Fisher Scientific. Flash chromatography was performed using Teledyne Isco CombiFlash systems using Teledyne Isco RediSep Rf silica and C18 columns. Nuclear Magnetic Resonance (NMR) spectra were recorded on a Varian INOVA 600 MHz NMR spectrometer (600 MHz $^{1}$H reference frequency, 151 MHz for $^{13}$C) equipped with an HCN indirect-detection probe or a Bruker AV 500 (500 MHz) spectrometer at Cornell University's NMR facility. $^{1}$H NMR chemical shifts are reported in ppm (δ) relative to residual solvent peaks (7.26 ppm for $CDCl_3$ and 3.31 ppm for $CD_3OD$). $^{1}$H NMR chemical shifts are reported as follows: chemical shift, multiplicity (s, singlet; d, doublet; t, triplet; m, multiplet), coupling constants (Hz) and integration. $^{13}$C NMR chemical shifts are reported in ppm (δ) relative to residual solvent peaks (77.16 ppm for $CDCl_3$ and 49.00 ppm for $CD_3OD$). All synthetic products were characterized by 2D NMR, including either gCOSY or non-gradient dqfCOSY spectra as well as HSQC and HMBC. Non-gradient phase-cycled dqfCOSY spectra were acquired using the following parameters: 0.6 s acquisition time; 400-600 complex increments; 8, 16 or 32 scans per increment. HSQC and HMBC spectra were acquired with these parameters: 0.25 s acquisition time, 200-500 increments, 8–64 scans per increment. $^{1}$H, $^{13}$C-HMBC spectra were optimized for $J_{H,C}$ = 6 Hz. NMR spectra were processed and baseline corrected using MestreLabs MNOVA Mnova v.14.2.3.

#### Synthesis of (*E*)-*N*-(3-((4-((3-aminopropyl)amino)butyl)amino)propyl)-3-(3,4-dimethoxyphenyl)acrylamide

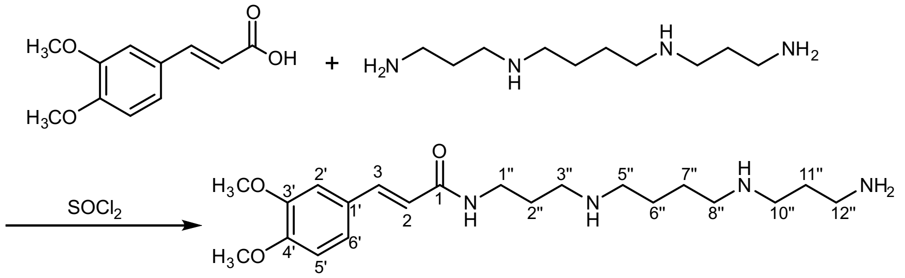


60 mg (0.28 mmol) of (*E*)-3-(3,4-dimethoxyphenyl)acrylic acid were treated with 1 mL of thionyl chloride for 2 h at 20 °C. The excess thionyl chloride was evaporated *in vacuo* without heating and the residue was dissolved in 15 mL of dry dichloromethane (DCM) under argon. The resulting slightly yellow solution was added to a stirred suspension of spermine (0.25 mmol) in 5 mL of DCM over a time period of 5 min. The solution was stirred for an additional 30 min and evaporated to dryness. The residue was analyzed by HPLC-MS and 2D NMR without further purification.

$^{1}$H NMR ($CD_3OD$, 500 MHz): δ [ppm] 7.49 (d, 1H, *J* = 16 Hz, 3-H), 7.25-7.2 (m, 2H, 2'- and 6'-H), 7.03 (d, 1H, *J* = 8.3 Hz, 5'-H), 6.54 (d, 1H, *J* = 16 Hz, 2-H), 3.89 (s, 3H, $CH_3O$), 3.88 (s, 3H, $CH_3O$), 3.44 (t, 1H, *J* = 6.6 Hz, 1"-H), 3.09-2.98 (m, 10H), 1.96-2.07 (m, 4H, 2"-H and 11"-H), 1.85-1.74 (m, 4H, 6"-H and 7"-H).

$^{13}C$ NMR ($CD_3OD$, 126 MHz): δ [ppm] 170.31 (C-1), 150.20, 151.89 (C-3' and C-4'), 142.58 (C-3), 128.77 (C-1'), 123.85 (C-6'), 119.10 (C-2), 112.86 (C-5'), 111.32 (C-2'), 56.7-56.8 (2C, $CH_3O$), 46.5-46.6 (2C, C-3" and C10"), 48.6-48.7 (2C, C-5" and C-8"), 38.6 (C-12"), 37.48 (C-1"), 27.66 (C-2"), 26.45 (C-11"), 25.1-25.3 (2C, C-6" and C-7").

### Synthesis[1,2] of *N*-(4-aminobutyl)-3-(hydroxyphenyl)propanamide

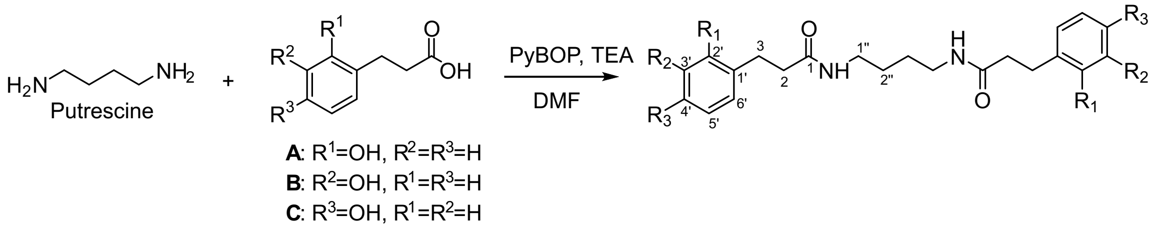


Putrescine dihydrochloride (30 mg, 0.186 mmol, 1 eq) and hydroxyphenyl propionic acid (62 mg, 0.372 mmol, 2 eq) were dissolved in dry dimethylformamide (DMF, 1 mL) in an 8 mL via. Then triethylamine (TEA, 103 µL, 0.744 mmol, 4 eq) and PyBOP (193.6 mg, 0.372 mmol, 2 eq) were added, and mixture was stirred for 3 h and concentrated to dryness. Three reactions were performed, using the 2-, 3-, or 4-hydroxypropionic acid (**A**, **B**, or **C**, respectively). The resulting products were used to prepare 50 µM samples in MeOH for HPLC-MS analysis. Following confirmation that the *meta*-isomer (derived from 3-hydroxycinnamic acid, C) corresponds to the major natural isomer, this compound was isolated via silica gel column chromatography, using a DCM/MeOH solvent gradient with 0.5% TEA. Yield: 50 mg (69.9 %) of a yellow oil, containing small amounts of HOBt and pyrrolidine as impurities.

$^{1}H$ NMR ($CD_3OD$, 500 MHz): δ [ppm] 7.06 (t, *J* = 8 Hz, 2H, 5'-H), 6.66 (m, 2H, 6'-H), 6.64 (m, 2H, 2'-H), 6.60 (m, 2H, 4'-H), 3.12 (m, 4H, 1"-H), 2.82 (t, *J* = 7.6 Hz, 4H, 3-H), 2.43 (t, *J* = 7.6 Hz, 4H, 2-H), 1.82 (m, 4H, 2"-H).

$^{13}C$ NMR ($CD_3OD$, 126 MHz): δ [ppm] 174.85 (2C, C-1), 158.18 (2C, C-3'), 143.35 (2C, C-1'), 130.11 (2C, C-5'), 120.32 (2C, C-6'), 116.03 (2C, C-2'), 113.80 (2C, C-4'), 47.07 (2C, C-1"), 38.65 (2C, C-2), 32.67 (2C, C-3), 27.04 (2C, C-2").

### Synthesis of *N*-(3-aminopropyl)-*N*-(4-((3-aminopropyl)amino)butyl)-3-(3-hydroxyphenyl)propanamide

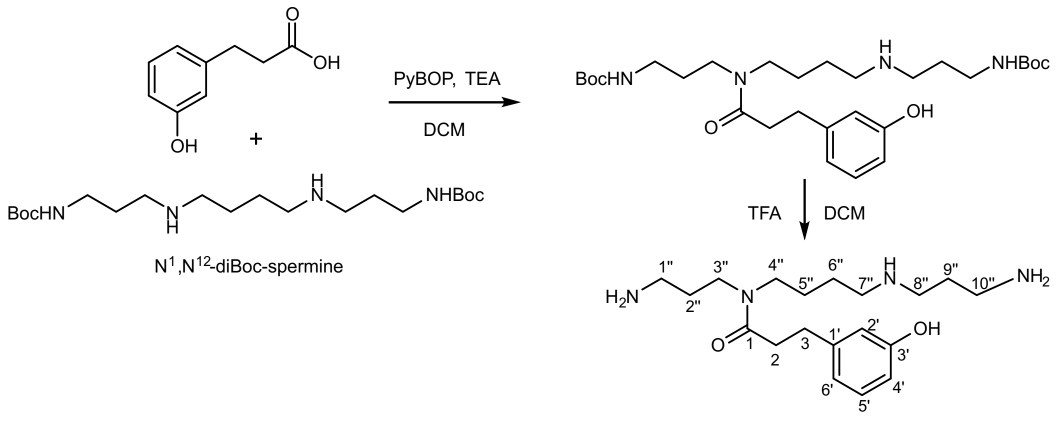

$N^1$,$N^{12}$-diBoc-spermine (500 mg, 1.24 mmol, 1 eq) and 3-(3-hydroxyphenyl)propionic acid (185 mg, 1.12 mmol, 0.9 eq) were dissolved in a dry round bottom flask charged with stir bar in 20 mL of dry DCM. TEA (691 µL, 4.96 mmol, 4 eq) and PyBOP (645 mg, 1.24 mmol, 1 eq) were then added sequentially. The mixture was then stirred at room temperature overnight, concentrated to dryness, and purified via silica gel combiflash chromatography (MeOH/DCM +0.5% TEA) to afford 515 mg Boc-protected product, which was then dissolved in 5 mL TFA/DCM = 1:1 and stirred at room temperature for 1 hour. The resulting mixture was concentrated to dryness and purified using C18 reverse phase combiflash chromatography (MeCN/$H_2O$ +0.1% formic acid) to afford 150 mg product.

$^1$H NMR ($CD_3OD$, 500 MHz): δ [ppm] 7.08 (t, $J$ = 8 Hz, 1H, 5-H), 6.68 (m, 1H, 6-H), 6.65 (m, 1H, 2-H), 6.62 (m, 1H, 4-H), 3.36 (m, 2H, 4"-H), 3.35 (m, 2H, 7"-H, 3.26 (m, 2H, 3"-H). 3.164 (m, 2H, 10"-H). 3.02 (m, 2H, 1"-H), 3.00 (m, 2H, 8"-H), 2.85 (t, $J$ = 7.6 Hz, 2H, 3-H), 2.64 (t, $J$ = 7.6 Hz, 2H, 2-H), 1.85 (m, 2H, 9"-H), 1.67 (m, 2H, 2"-H), 1.66-1.59 (4H, 5"-H and 6"-H).

$^{13}$C NMR ($CD_3OD$, 126 MHz): δ [ppm] 174.52 (C-1), 158.29 (C-3'), 143.52 (C-1'), 130.24 (C-5'), 120.44 (C-6'), 116.04 (C-2'), 113.81 (C-4'), 46.61 (C-3"), 46.29 (C-8"), 45.85 (C-4"), 44.44 (C-7"), 38.3 (C-1"), 37.56 (C-10"), 35.42 (C-2), 32.45 (C-3), 30.11 (C-2"), 27.80 (C-9"), 25.41 and 24.20 (C-5" and C-6").

**Synthesis of (*E*)-*N*-(3-(*N*-(4-((3-aminopropyl)amino)butyl)-3-(3-hydroxyphenyl)-propanamido)propyl)-3-(3,4-dimethoxyphenyl)acrylamide and (*E*)-*N*-(3-((4-(*N*-(3-aminopropyl)-3-(3-hydroxyphenyl)propanamido)butyl)amino)propyl)-3-(3,4-dimethoxyphenyl)acrylamide**

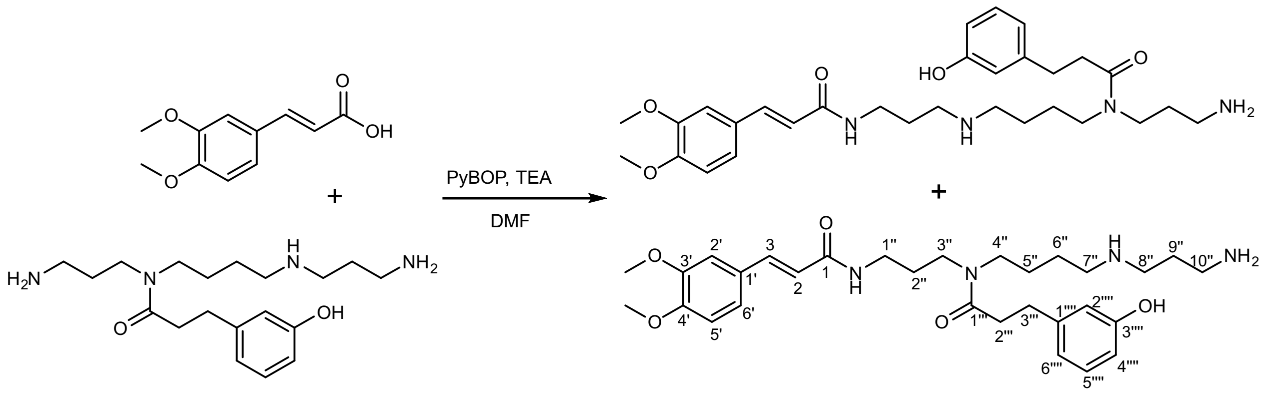


*N*-(3-aminopropyl)-*N*-(4-((3-aminopropyl)amino)butyl)-3-(3-hydroxyphenyl)propanamide (97 mg, 0.276 mmol, 1 eq) was dissolved in 2 mL dry DMF in a 20 mL vial charged with stir bar. In a separate dry 4 mL vial charged with a stir bar, (*E*)-3,4-dimethoxycinnamic acid (57.6 mg, 0.276 mmol, 1 eq) was dissolved in 2 mL dry DMF, then TEA (131.2 µL, 0.941 mmol, 3.4 eq) and PyBOP (158 mg, 0.304 mmol, 1.1 eq) were added sequentially. This mixture was stirred for 10 min before being taken up in a total of 3 mL dry DMF in a 3 mL syringe. Then a syringe pump was used to add the activated acid to the stirring solution of *N*-(3-aminopropyl)-*N*-(4-((3-aminopropyl)amino)butyl)-3-(3-hydroxyphenyl)propanamide over a span of 2 hours. The reaction was then concentrated to dryness and purified via C18 reverse phase combiflash chromatography (MeCN/$H_2O$ +0.1% formic acid) to afford a mixture of the two desired products (13.5 mg).

(*E*)-*N*-(3-((4-(*N*-(3-aminopropyl)-3-(3-hydroxyphenyl)propanamido)butyl)amino)propyl)-3-(3,4-dimethoxyphenyl)acrylamide

$^1$H NMR ($CD_3OD$, 600 MHz): δ [ppm] 7.50 (d, 1H, *J* = 16 Hz, 3-H), 7.17-7.11 (m, 2H, 2'- and 6'-H), 6.97 (d, 1H, *J* = 8.3 Hz, 5'-H), 6.50 (d, 1H, *J* = 16 Hz, 2-H), 3.86 (2s, 6H, $CH_3O$), 3.42 (m, 2H, 1''-H), 1.94 (m, 2H, 2''-H), 3.03 (m, 2H, 3''-H), 2.98 (m, 2H, 4''-H), 1.66 (m, 2H, 5''-H), 1.55 (m, 2H, 6''-H), 3.29 (m, 2H, 7''-H, HMBC to 175.38), 3.44 (m, 2H, 8''-H, HMBC to 175.38), 1.86 (m, 2H, 9''-H), 2.82 (m, 2H, 10''-H), 7.08 (t, *J* = 8 Hz, 1H, 5-H''''), 6.69 (m, 1H, 6-H''''), 6.67 (m, 1H, 2-H''''), 6.62 (m, 1H, 4-H''''), 2.87 (t, *J* = 7.6 Hz, 4H, 3-H'''), 2.69 (t, *J* = 7.6 Hz, 4H, 2-H''').

$^{13}$C NMR ($CD_3OD$, 151 MHz): δ [ppm] 169.56 (C-1), 150.31, 151.92 (C-3' and C-4'), 142.06 (C-3), 128.77 (C-1'), 123.07 (C-6'), 118.58 (C-2), 112.21 (C-5'), 111.00 (C-2'), 56.0-56.2 (2C, $CH_3O$), 36.65 (C-1''), 27.42 (C-2''), 45.92 (C-3''), 48.10 (C-4''), 24.17 (C-5''), 26.55 (C-6''), 48.32 (C-7''), 43.13 (C-8''), 26.46 (C-9''), 37.53 (C-10''), 175.38 (C-1'''), 158.07 (C-3''''), 143.38 (C-1''''), 130.20 (C-5''''), 120.15 (C-6''''), 115.94 (C-2''''), 113.81 (C-4''''), 32.20 (C-3'''), 35.19 (C-2''').

(*E*)-*N*-(3-(*N*-(4-((3-aminopropyl)amino)butyl)-3-(3-hydroxyphenyl)propanamido)propyl)-3-(3,4-dimethoxyphenyl)acrylamide

$^1$H NMR ($CD_3OD$, 600 MHz): δ [ppm] 7.46 (d, 1H, *J* = 16 Hz, 3-H), 7.17-7.11 (m, 2H, 2'- and 6'-H), 6.97 (d, 1H, *J* = 8.3 Hz, 5'-H), 6.47 (d, 1H, *J* = 16 Hz, 2-H), 3.86 (2s, 6H, $CH_3O$), 3.28 (m, 2H, 1''-H), 1.77 (m, 2H, 2''-H), 3.30, (m, 2H, 3''-H, HMBC to 174.4), 3.37 (m, 2H, 4''-H, HMBC to 174.4), 1.61 (m, 2H, 5''-H),1.65 (m, 2H, 6''-H), 3.03 (m, 2H, 7''-H), 3.09, m, 2H, 8''-H), 2.06 (m, 2H, 9''-H), 3.05 (m, 2H, 10''-H), 7.08 (t, *J* = 8 Hz, 1H, 5-H''''), 6.69 (m, 1H, 6-H''''), 6.67 (m, 1H, 2-H''''), 6.62 (m, 1H, 4-H''''), 2.83 (t, *J* = 7.6 Hz, 4H, 3-H'''), 2.64(t, *J* = 7.6 Hz, 4H, 2-H''').

$^{13}$C NMR ($CD_3OD$, 151 MHz): δ [ppm] 168.60 (C-1), 150.31, 151.92 (C-3' and C-4'), 141.43 (C-3), 128.77 (C-1'), 123.07 (C-6'), 118.96 (C-2), 112.21 (C-5'), 111.00 (C-2'), 56.0-56.2 (2C, $CH_3O$), 37.56 (C-1''),29.44 (C-2''),46.61 (C-3''), 45.81 (C-4''), 25.25 (C-5''), 24.10 (C-6''), 48.30 (C-7''), 45.36 (C-8''), 25.11 (C-9''), 37.43 (C-10''), 174.39 (C-1'''), 158.07 (C-3''''), 143.38 (C-1''''), 130.20 (C-5''''), 120.15 (C-6''''), 115.94 (C-2''''), 113.81 (C-4''''), 32.44 (C-3'''), 35.37 (C-2''').

**Synthesis[3] of 8-(3,4-dimethoxyphenyl)-1,5,9,13-tetraazacycloheptadecan-6-one**

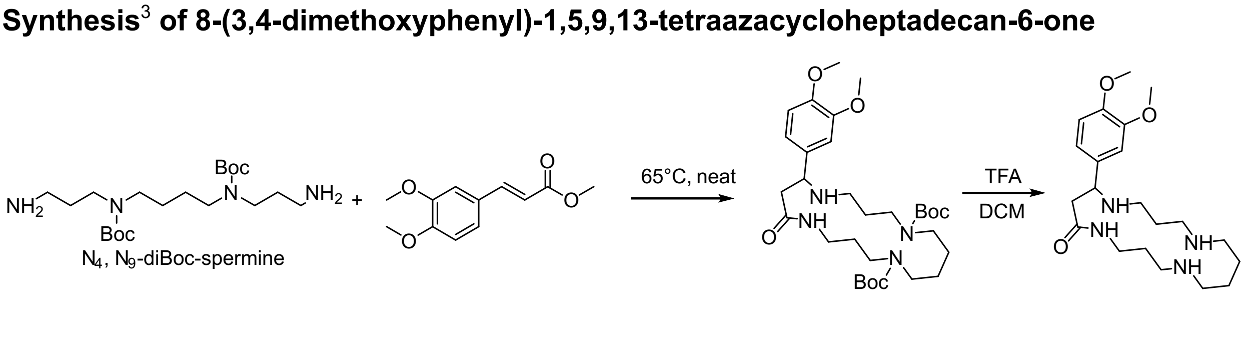


*N*[4],*N*[9]-diBoc-spermine (17 mg, 0.042 mmol, 1 eq) and 3,4-dimethoxy-3-(phenyl)propionic acid methyl ester (4.76 mg, 0.021 mmol, 0.5 eq) were transferred to a dry 4 mL vial as

MeOH stocks ($N^4$,$N^9$-diBoc-spermine 94.5 mg/mL; 3,4-dimethoxy methyl ester 4.76 mg/mL). MeOH was then removed via rotatory evaporation to yield a dry mixture of the two starting materials. Said mixture was heated to 65 °C in a sealed 4 mL vial in a copper block for 48 hours. Subsequently the mixture was allowed to cool to room temperature and dissolved in 1 mL TFA/DCM = 1:1 and stirred at room temperature for 30 minutes. The solution was subsequently concentrated to dryness and the products were redissolved into MeOH at a concentration of 500 μM for HPLC and NMR spectroscopic analysis.
