## Supplementary User Guide for "Charting the small-molecule universe from mass spectra with neuro-symbolic AI"

**Table of Contents**

Supplementary User Interface Guide
Search Page Guide 4
Visualizer Guide 10

### Supplementary User Interface Guide

AIMe user interface step-by-step guide detailing each field, inputs and outputs.

There are two main use-case scenarios:

1- From the Home page, you can select the “Search” icon that redirects to a page where you can:
   a. upload a .mgf file for a single compound (multiple spectra for different collision energies are allowed)
   b. manually input a peak list and respective parameters.

2- Select the “Visualize” button. This allows for .json files previously downloaded from AIMe UI to be loaded, bypassing the search page.

- Home: https://www.cs.cornell.edu/gomes/udiscoverit/kosmos/
- Search: https://www.cs.cornell.edu/gomes/udiscoverit/kosmos/search.html
- Visualize: https://www.cs.cornell.edu/gomes/udiscoverit/kosmos/visualize.html

For a list of .json files available, see the “Source data” section.

#### Quick “Search page” overview:

| Step | Description | Additional information |
|---|---|---|
| 1 | Fill in the experiment metadata manually or upload an MGF/MSP/TXT file | Precursor *m/z* or molecular formula are mandatory inputs and are automatically populated if imported from a file. The Experiment name and InChiKey are optional inputs. |
| 2 | Enter one or more $MS^2$ spectra, including peak list, instrument, ionization mode and collision energy. | The search request requires ionization mode (M+H and M-H only), suitable for any orbitrap/High resolution MS instrument and collision energy |
| 3 | Optional setting:<br>“*User-defined candidates*”.<br>Enter one SMILES string per line. No adduct modification or ionization unless the compound is constitutively charged | The tool predicts spectra only for the supplied candidate molecules, rather than running a search over PubChem-predicted spectra.<br>The button at the end will change to *“Validate candidates”* |
| 4 | Select search settings:<br>• “Isomer/Identity search” or “Full search”<br>• “Precomputed” or “Predict spectra”<br>• Top K | The 6 parameters at the end of this section define criteria to clean up and process spectra. In addition, a check of every peak is made to assess consistency with the precursor molecular formula. |
| 5 | Click “Predict spectra” | An initial candidate list is displayed with links to PubChem entries. |
| 6 | Click “Open Visualizer” | The search results can also be downloaded as a .json file |

**Search Page:** Auto-Fill and Metadata

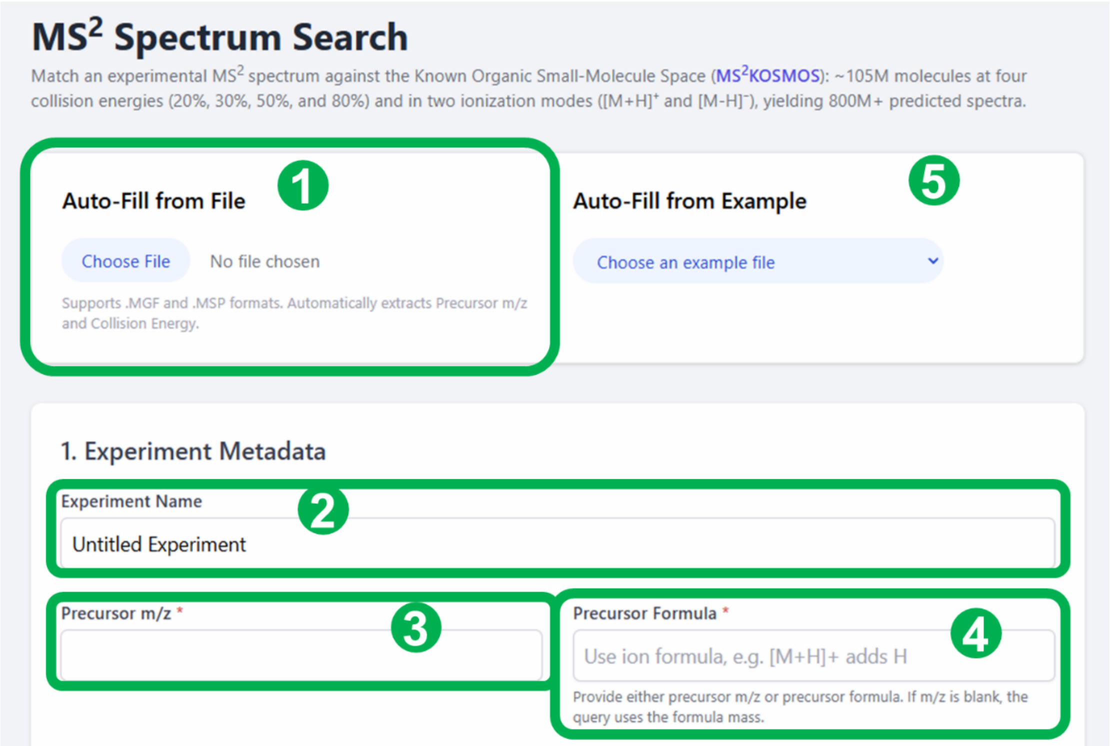


| # | Field | Description | Use/info |
|---|---|---|---|
| 1 | Auto-Fill from File (optional) | File picker for .MGF, .MSP, and .TXT spectrum files. | Use this when a supported spectrum file is available. The page can populate precursor m/z and collision energy from compatible files. Files can be downloaded from Mass Bank of North America, GNPS etc |
| 2 | Experiment Name (optional) | Human-readable name for the query. | -- |
| 3 | Precursor *m/z* | Observed precursor ion mass-to-charge ratio. | -- |
| 4 | Precursor Formula | Molecular formula consistent with precursor *m/z* | Enter the molecular formula for the precursor ion, not the neutral molecule. The placeholder reminds you that [M+H]+ adds H. |
| 5 | Auto-fill | Already predefined fields that get automatically populated | Click on the drop-down menu and select one of the examples |

**Search Page:** Spectra input and Candidate Validation

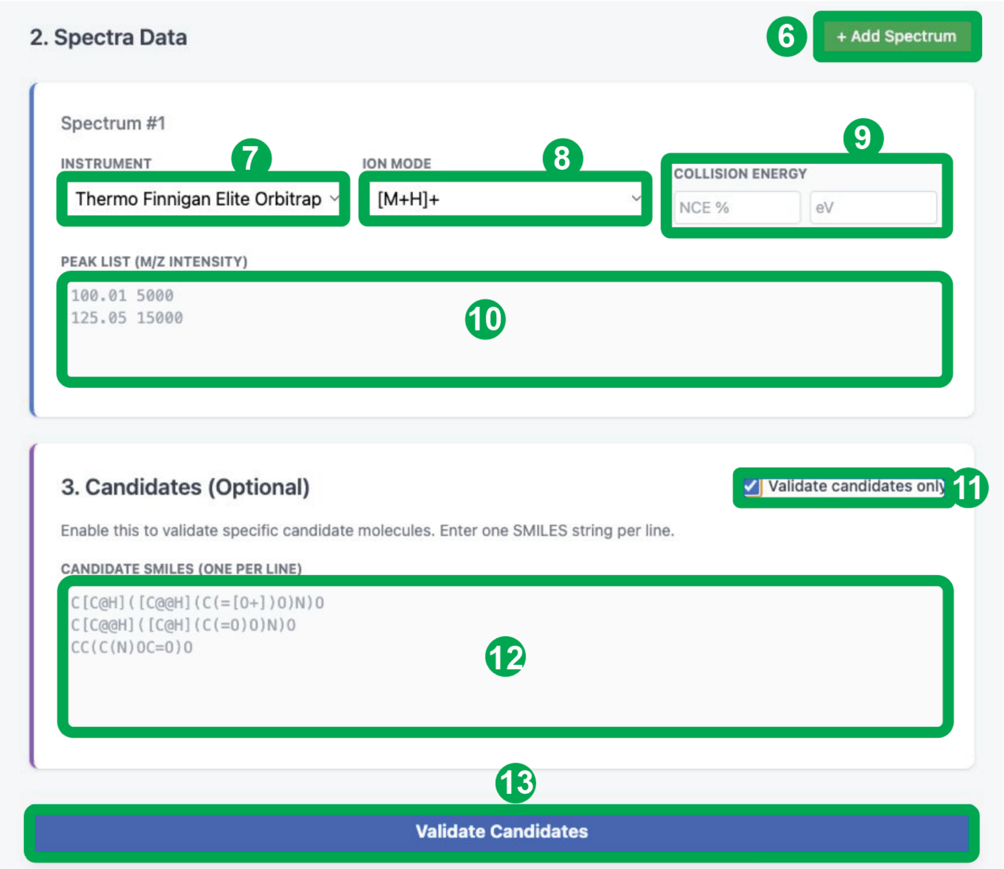


| # | Field | Description | Use/info |
|---|---|---|---|
| 6 | Add Spectrum | Adds another spectrum block for cases where multiple collision energies are available | -- |
| 7 | Instrument | Instrument selection is the default Orbitrap defined by the training data. | The current version does not support instrument-specific parameters. However, in the future, this may extend to other High Resolution Mass Spectrometer options. |
| 8 | Ion Mode | Adduct/ion mode selector. | Choose $[M+H]^+$ for positive mode protonated ions or $[M-H]^-$ for negative mode deprotonated ions. |
| 9 | Collision Energy | Two numeric fields: NCE percent and eV. | Enter NCE or eV matching those of the experimental spectra. Stepped collision energy is not yet supported. Integer values are expected. |
| 10 | Peak List | Experimental $MS^2$ peaks as *m/z* intensity pairs. | Enter one peak per line. |
| 11 | Validate candidates only | Switches the workflow from broad search to a validation of user input molecules. | Enable to test specific candidate SMILES. This setting enables de-novo prediction of candidate molecules, even if absent from MS2KOSMoS. |
| 12 | Candidate SMILES | One candidate molecule per line. | This field is disabled until validation is enabled. Paste plain SMILES strings only. One string per line. The SMILES string should be the neutral/unionized form unless the compound is constitutively charged. |
| 13 | Validate Candidates | Runs the workflow according to the set parameters | Appears when validation mode is enabled; search options are hidden in this mode. |

#### **Search Page:** Search Options

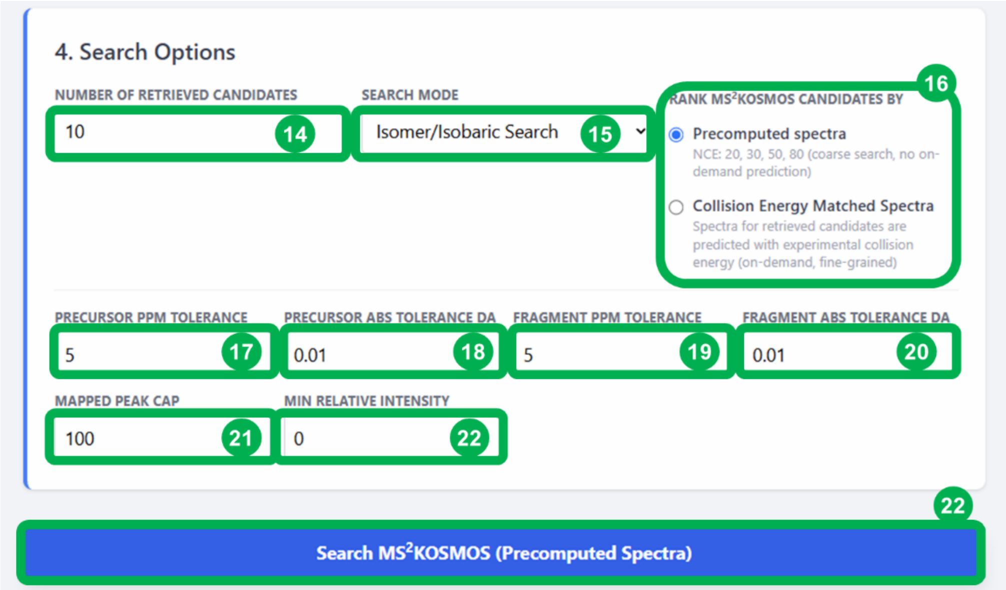


| # | Field | Description | Use/info |
|---|---|---|---|
| 14 | Number of retrieved candidates | Maximum number of candidates to return. | Use a small number for quick review or increase it when you need broader candidate coverage. |
| 15 | Search Mode | Select “*Isomer/Isobaric Search”* or *“Full Search”* | Use Isomer Search for precursor-constrained or Molecular formula constrained lookup. Use Full Search when a broader search is desired. |
| 16 | Rank candidates mode | Chooses precomputed database spectra or model-predicted spectra. **Precomputed search does not contain fragment annotations or fragmentation tree**. | “precomputed” mode uses precomputed NCE 20%, 30%, 50%, 80%, spectra for the entire predicted $MS^2$ KOSMoS. “Predict” mode predicts the candidate spectra at custom collision energy after a behind the scenes pre-computed search and recalculates cosine similarities and re-ranks the candidates. The pre-computed search is always done to the closest experimental collision energy input. |
| 17 | Precursor ppm tolerance | Relative mass tolerance around precursor *m/z.* | Lower values are stricter; raise only when instrument mass accuracy requires it. |
| 18 | Precursor abs tolerance Da | Absolute precursor mass tolerance. | Used together with ppm tolerance. Whichever value is largest is the defined tolerance (ppm/abs). |
| 19 | Fragment ppm tolerance | Relative tolerance for matching fragment peaks. | Controls how tightly fragment *m/z* values are matched during preprocessing/scoring. |
| 20 | Fragment abs tolerance Da | Absolute fragment mass tolerance. | Used together with ppm tolerance. Whichever value is largest is the defined tolerance (ppm/abs). |
| 21 | Mapped peak cap | Maximum number of processed peaks used for search/scoring. | Keeps scoring focused on the top N intensity peaks. |
| 22 | Min relative intensity | Minimum relative intensity filter from 0 to 1. | Raise this to ignore very minor peaks; leave at 0 to keep all processed peaks. |
| 23 | “Predict Spectra” | Starts the selected workflow. | The label changes based on mode: Search Precomputed Database, Predict Spectra, or Validate Candidates. |

**Search Page:** Run-progress visualization

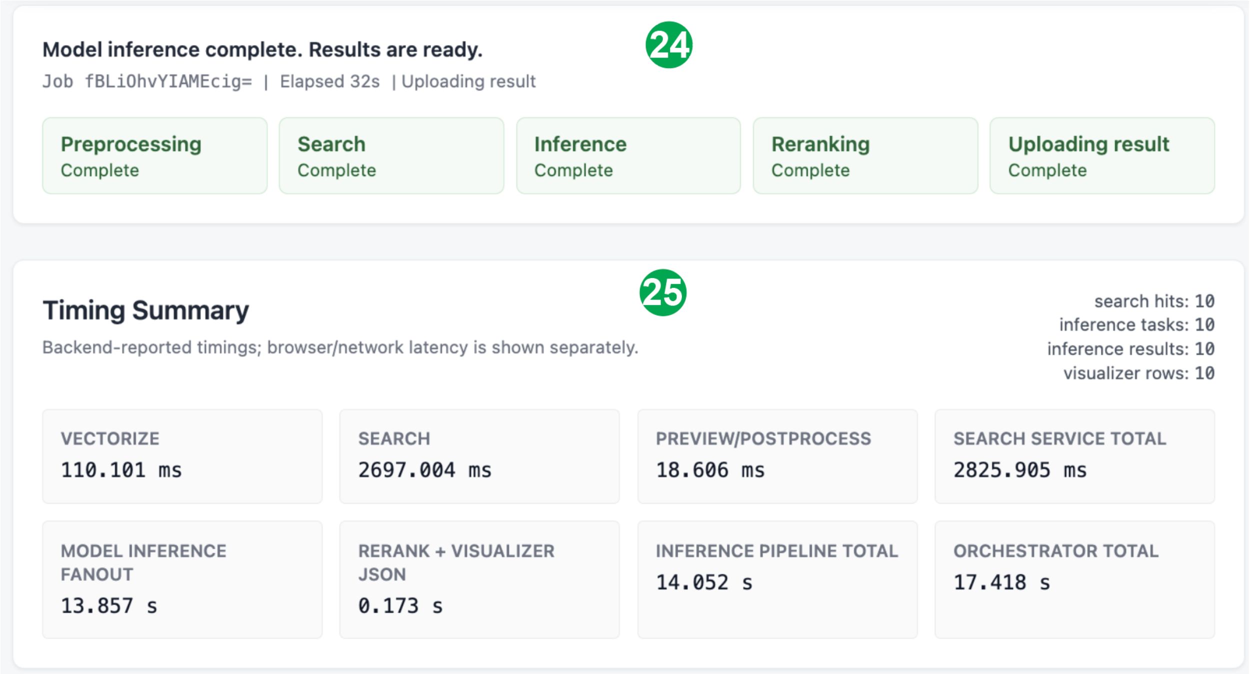


| # | Field | Description | Use/info |
|---|---|---|---|
| 24 | Step-by-step progress | Informs the user of the current step in progress with specific timings | -- |
| 25 | Timing summary | Displays the timings for each step as well and search counts. | -- |

#### **Search Page:** Processing output and data summary

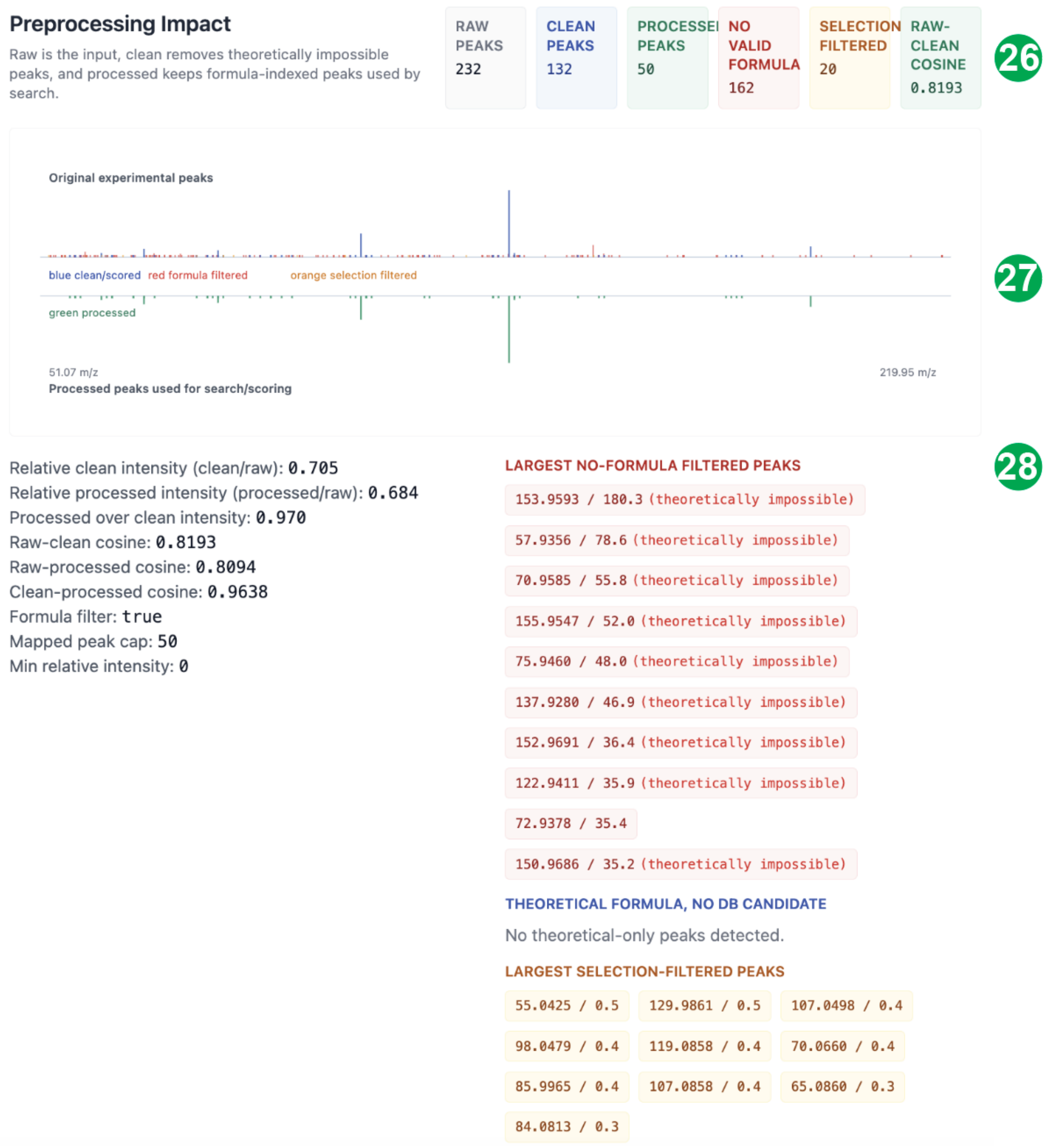


| # | Field | Description | Use/info |
|---|---|---|---|
| 26 | Processing summary | Describe number of input peaks, filtered out peaks, and final cosine score to the input spectrum | Additionally defines the number of peaks considered from the above parameter, and total number of selected peaks. |
| 27 | Spectral overview of the pre- and post-processing | Color coded spectrum of the input spectrum and processing effects | Top input bottom after processing |
| 28 | Peaks excluded | Separates the excluded peaks based on the types of filters applied | Reports each excluded peak intensity and *m/z*. |

**Search Page:** Candidate list summary

29

##### Model Inference Results

Candidates ranked after predicted-spectrum reranking. The vector-rank column shows where the same candidate appeared in the precomputed search.

10 candidates | 10 visible rows

| RANK | VECTOR RANK | CID | FORMULA | SMILES | MODEL COSINE |
|---|---|---|---|---|---|
| 1 | 7 | 66125590 | C8H10N4O2 | CC1=NN(C(=C1N)C(=O)OCC#N)C | 0.846739 |
| 2 | 3 | 89155469 | C6H7N3O | CC1=NC(=CN=C1N)C=O | 0.755665 |
| 3 | 9 | 122407817 | C6H9N3O2 | CNC1=CN(N=C1C(=O)O)C | 0.773858 |
| 4 | 10 | 66124983 | C11H17N7O | CC1=NN(C(=C1N)C(=O)NCCC2=NN=CN2C)C | 0.834701 |
| 5 | 6 | 158028248 | C6H7N3O | C1C=NC2C(C1=O)NC=N2 | 0.782111 |
| 6 | 5 | 171035350 | C6H7N3O | CC1=NC(=NC=C1N)C=O | 0.767302 |
| 7 | 2 | 161886215 | C8H13N5O | CN1C=CNC(=C1N)C(=O)NC(=C)N | 0.765412 |
| 8 | 1 | 88858035 | C6H7N3O | CC1=NC(=NC(=N1)C=O)C | 0.802713 |
| 9 | 8 | 83913941 | C6H7N3O | CC1=NC(=CN1C)N=C=O | 0.808406 |
| 10 | 4 | 13114149 | C6H7N3O | CC1(C=NN(C1=O)C)C#N | 0.749361 |

30

##### Precomputed Vector Results

ce50 | full | 2825.905 ms

10 candidates | 10 visible rows

| RANK | MODEL RANK | CID | FORMULA | SMILES | PREVIEW COSINE / SCORE |
|---|---|---|---|---|---|
| 1 | 8 | 88858035 | C6H7N3O | CC1=NC(=NC(=N1)C=O)C | 0.827495 |
| 2 | 7 | 161886215 | C8H13N5O | CN1C=CNC(=C1N)C(=O)NC(=C)N | 0.826360 |
| 3 | 2 | 89155469 | C6H7N3O | CC1=NC(=CN=C1N)C=O | 0.819229 |
| 4 | 10 | 13114149 | C6H7N3O | CC1(C=NN(C1=O)C)C#N | 0.813857 |
| 5 | 6 | 171035350 | C6H7N3O | CC1=NC(=NC=C1N)C=O | 0.813680 |
| 6 | 5 | 158028248 | C6H7N3O | C1C=NC2C(C1=O)NC=N2 | 0.809594 |
| 7 | 1 | 66125590 | C8H10N4O2 | CC1=NN(C(=C1N)C(=O)OCC#N)C | 0.810661 |
| 8 | 9 | 83913941 | C6H7N3O | CC1=NC(=CN1C)N=C=O | 0.809559 |
| 9 | 3 | 122407817 | C6H9N3O2 | CNC1=CN(N=C1C(=O)O)C | 0.808676 |
| 10 | 4 | 66124983 | C11H17N7O | CC1=NN(C(=C1N)C(=O)NCCC2=NN=CN2C)C | 0.808440 |

| # | Field | Description | Use/info |
|---|---|---|---|
| 29 | Candidate list from search matched to experimental collision energy | Details the retrieved compounds from PubChem and respective links. Cosine similarity to predicted spectra. | Unique identifiers for each retrieved compound and cosine similarity based on predicted spectra at the specified collision energy. Lists re-ranked Top-k candidates. |
| 30 | Candidate list from search precomputed spectra at specified collision energy | Details the unique identifiers and cosine similarity to the closest precomputed collision energy | Original ranking of precomputed search. Top-k retrieval |

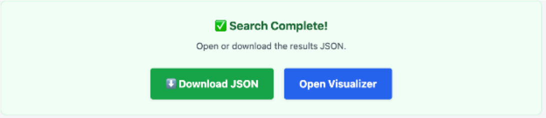


Download search results as .json file and/or open visualizer in a new window

**Visualizer Page:** Overview:

Use case scenarios:

1- Load spectra from pre-loaded examples

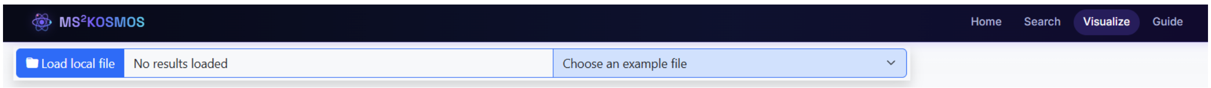


2- Load a .jason file from previously saved session

3- Display after clicking “Open Visualizer” in the search page.

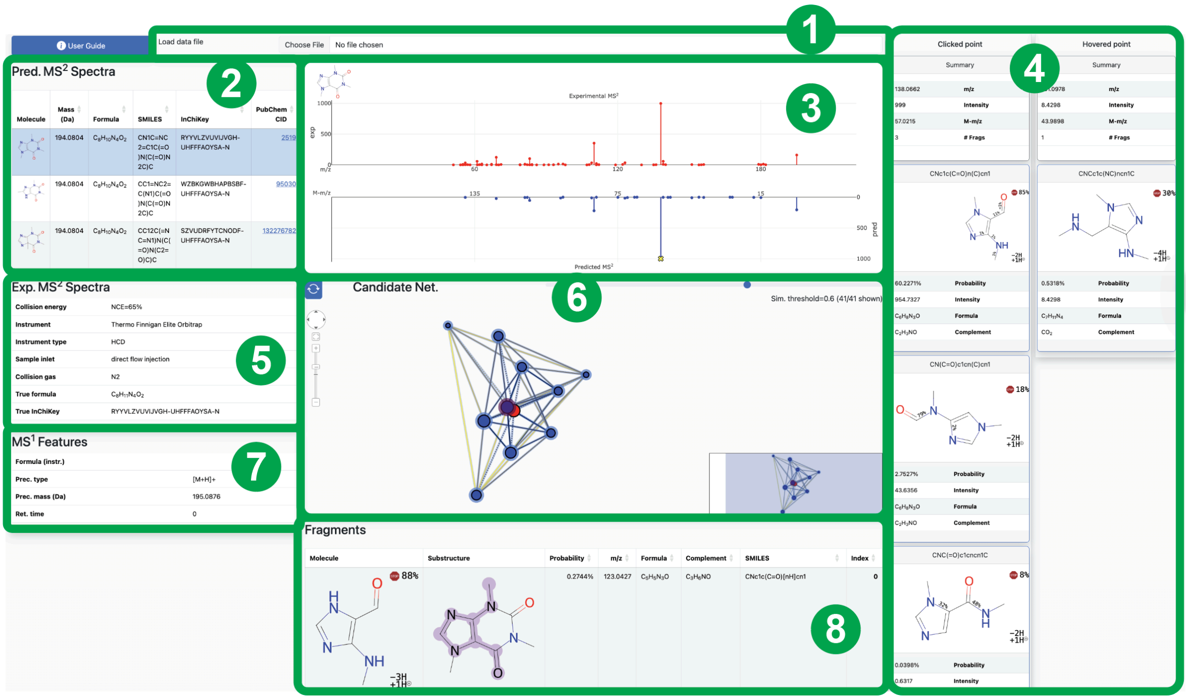


| # | Field | Description | Use/info |
|---|---|---|---|
| 1 | Load data file | Previously save sessions can be loaded and visualized again. | Loads previously saved session in a .json format. |
| 2 | Candidate list | Lists all the details for a given candidate | Rank, chemical identifier, structure, molecular formula, cosine and entropy similarity link to PubChem entry. |
| 3 | Spectra comparison plot | Experimental spectra at the top, predicted at the bottom. | Each predicted fragment is clickable. A yellow “x” denotes a clicked fragment. Tool tips on mouse hover show the predicted m/z and the experimental m/z when they match within 5 ppm. |
| 4 | Clicked and hovered point fragment cards | Left panels indicate the clicked fragment. Panels below show all possible structures that generated a clicked fragment. Right panel displays the same information but on hover. | This display allows comparison of two fragments side-by-side |

| | | | |
|---|---|---|---|
| 5 | Experimental $MS^2$ Spectra details | Experimental acquisition metadata. | Displays collision energy, instrument, etc. for the queried experimental spectrum |
| 6 | Fragmentation tree/candidate network display | Toggle display between fragmentation tree and candidate network. | The interface indicates whether you are currently viewing Candidate Net or Fragment Tree. Clicking the blue button in the top-left corner switches between these two views. Each element of both views is clickable, updating the spectrum viewer accordingly. |
| 7 | $MS^1$ details | Precursor m/z, adduct type and retention time is available | -- |
| 8 | Fragment list | All fragments generated for the selected candidate are displayed, with their respective structure, m/z, intensity, formula and SMILES. | -- |

#### **Visualizer Page:** Candidate table fields

| **Field** | **Description** | **Use/info** |
|---|---|---|
| Molecule | Rendered structure preview. | Quick visual comparison of candidate structures. |
| Mass (Da) | -- | -- |
| Formula | Candidate molecular formula. | -- |
| SMILES | Candidate structure as a SMILES string. | -- |
| InChiKey | Candidate structure as an InChIKey identifier. | -- |
| PubChem CID | PubChem compound identifier. | Click the linked CID to open the PubChem record. |
| Hungarian cosine | Average spectral similarity score. | Higher values generally indicate better experimental/predicted agreement. |
| Entropy | Alternative overall spectral similarity score. | Use alongside cosine when ranking candidates. |
| Hungarian cosine (individual) | Per-spectrum cosine score. | Useful when multiple spectra or energies are present. |
| Entropy (individual) | Per-spectrum entropy score. | Useful when multiple spectra or energies are present. |
| Index | Internal row/candidate index. | -- |

**Visualizer Page:** Mouse click & hover fragment cards

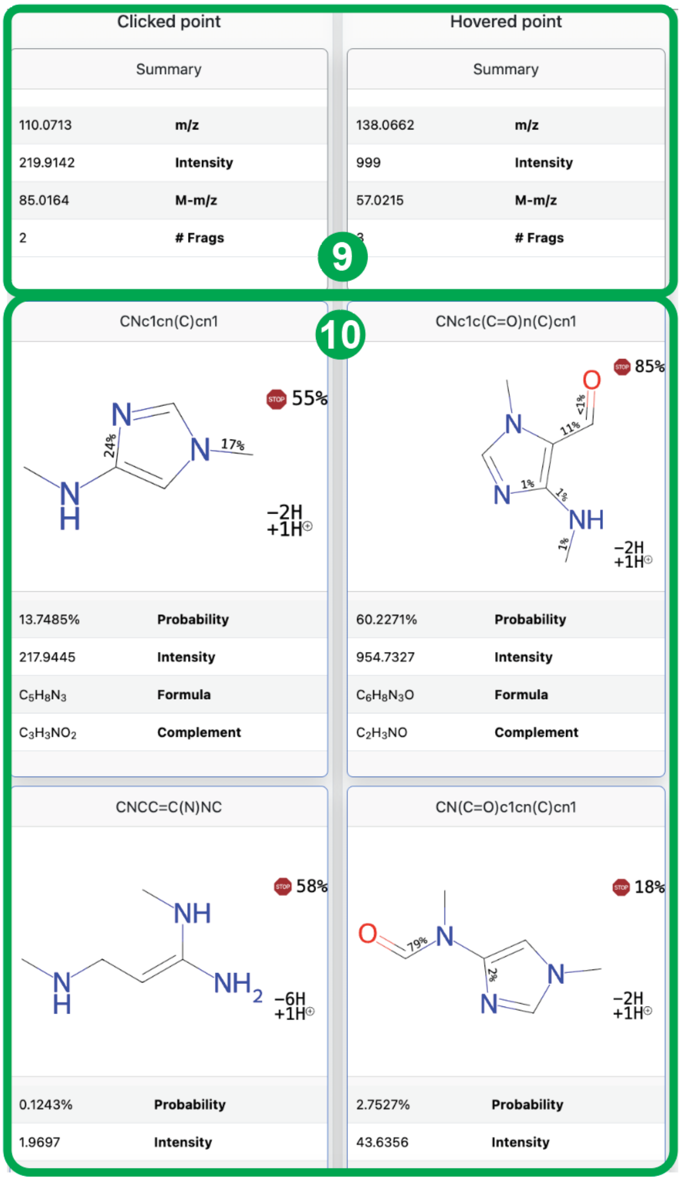


| # | Field | Description | Use/info |
|---|---|---|---|
| 9 | Summary | Pinned peak summary with m/z, intensity, and M-m/z (difference between precursor m/z and fragment m/z | Use after clicking a peak to keep a stable reference while reviewing fragments. |
| 10 | Fragment cards | Details the structure, and sequential bond-breaking probabilities or halting probability. | Number of implicit unsaturations on the bottom right corner. Probability of the fragment and intensity as well as molecular formula predicted by the model and the remaining molecular formula to the precursor (neutral loss-like) |

**Visualizer Page:** Fragmentation DAG and candidate network

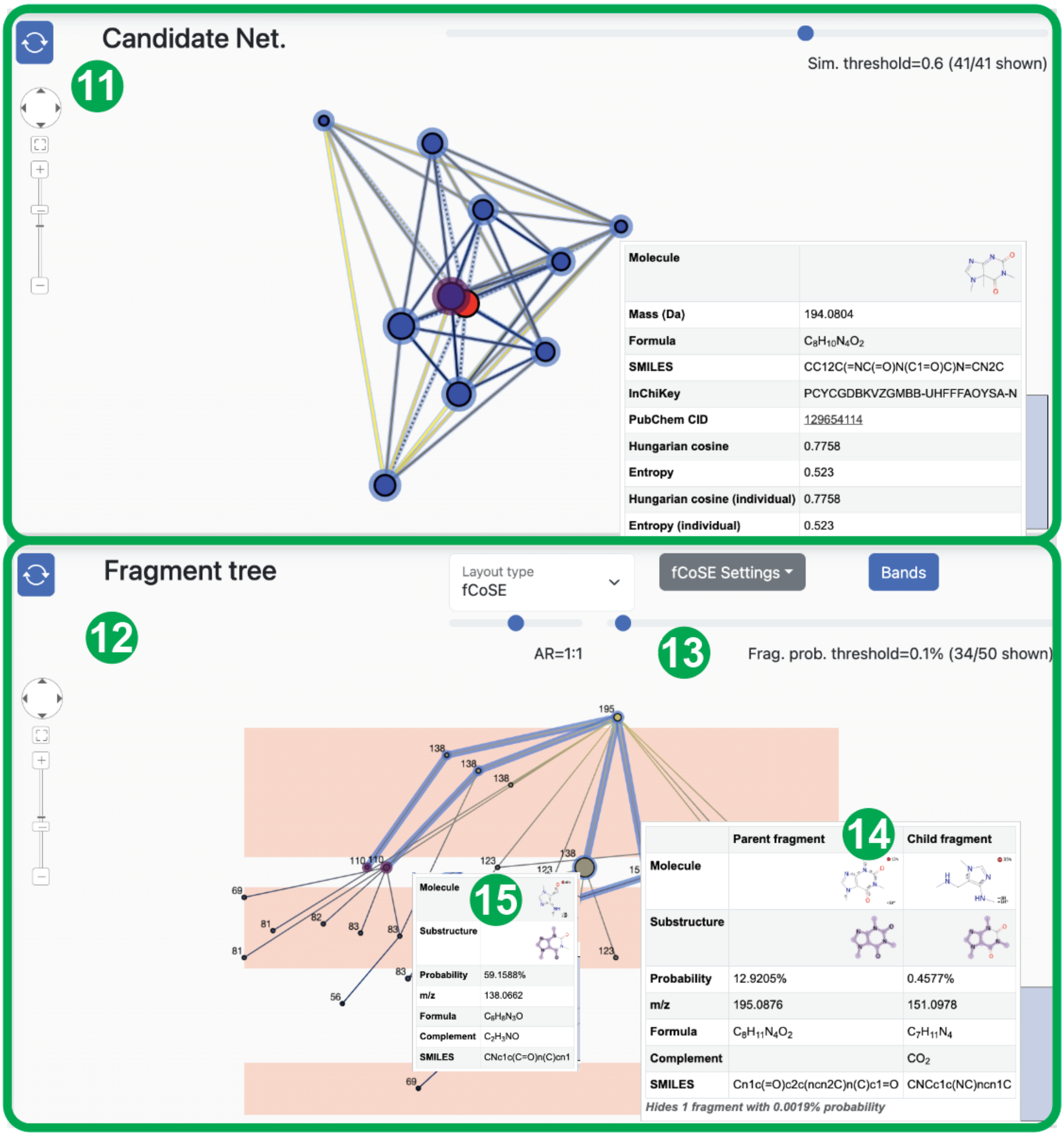


| # | Field | Description | Use/info |
|---|---|---|---|
| 11 | Network display | The center red node corresponds to the experimental spectrum. Remaining nodes are displayed according to their similarity. | Clicking on a node will update the spectrum visualizer to that candidate. Hover tool tip displays candidate information. |
| 12 | Network/Fragmentation DAG toggle | Switch between views | -- |
| 13 | DAG layout options | Slider defines the probability threshold for nodes. | The drop-down menu allows further customization of display options, especially useful for large DAGs |
| 14 | Mouse hover edge tool | Allows for visualization of the | Structure and neutral loss are displayed as |

| | | | |
|---|---|---|---|
| | tip | parent fragment and child fragment in one tool tip | a purple highlighted color alongside fragment details. |
| 15 | Mouse hover node display | Allows for visualization of the structure of the fragment and its corresponding substructure from the parent molecule. | -- |

#### **Visualizer Page:** Fragment table

##### Fragments

| Molecule | Substructure | Probability | m/z | Formula | Complement | SMILES | Index |
|---|---|---|---|---|---|---|---|
| 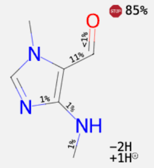 | 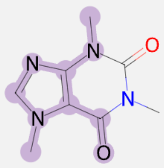 | 59.1588% | 138.0662 | $C_6H_8N_3O$ | $C_2H_3NO$ | CNc1c(C=O)n(C)cn1 | **21** |
| 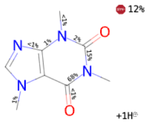 | 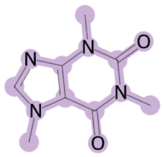 | 12.9205% | 195.0876 | $C_8H_{11}N_4O_2$ | | Cn1c(=O)c2c(ncn2C)n(C)c1=O | **48** |
| 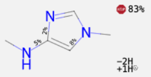 | 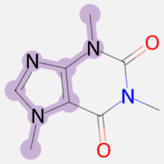 | 9.3379% | 110.0713 | $C_5H_8N_3$ | $C_3H_3NO_2$ | CNc1cn(C)cn1 | **42** |
| 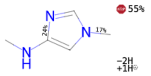 | 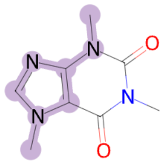 | 4.2436% | 110.0713 | $C_5H_8N_3$ | $C_3H_3NO_2$ | CNc1cn(C)cn1 | **18** |

| # | Field | Description |
|---|---|---|
| All predicted fragments can be visualized in this table | Similar to the fragment cards above, the structure and bond-breaking probabilities are displayed. The shaded area represents one possible substructure from the previous step. | The table can be sorted by *m/z* and probability.<br>For identical fragments that originate from different fragmentation pathways, their molecular formula will be the same, with different assigned probabilities that all contribute to the spectra prediction. |
